# Bivalent Inhibitor Development for a Dynamic Epigenetic Tandem Reader Protein Using a Combinatorial Histone Code

**DOI:** 10.64898/2026.10.09.758058

**Authors:** Joseph E. McPherson, Molly S. Sneddon, Richard K. Begyinah, Kesavan Babu, Kerstin E. Peterson, Ryan F. Friess, Thomas G. Smith, Natalia Y. Tretyakova, Marcus Fischer, William C.K. Pomerantz

## Abstract

Epigenetic readers are a class of proteins that recognize specific chemical modifications on histones to differentially regulate gene expression. Aberrant expression of readers can drive the progression of many diseases and cancers, making this protein class viable therapeutic targets. Understanding the landscape of histone modifications recognized by reader domains, and the structural mechanisms that drive these molecular recognition events, is crucial for guiding the design of pharmaceutically relevant chemical probes. BPTF contains a plant homeodomain (PHD) finger and tandem bromodomain (BRD). Here, we expand upon previous studies that found the BPTF BRD can engage acetyllysine modifications on the N-terminus of histone H3. We uncover H3K4me_3_K14ac as the specific modifications recognized by the PHD-BRD cassette, which is confirmed by crystallography. Using orthogonal methods, our work explores how these modifications can be read cooperatively, including in *cis.* We report the first experimental evidence proposing a ligand-induced conformational change between the two reader domains of BPTF, supporting previous computational approaches that have predicted a macromolecular rearrangement. We leverage these structural insights to rationally design a peptide-small molecule conjugate **MJ-1** that inhibits both domains of BPTF and overcomes ligandability challenges associated with PHD-finger-containing proteins. The results presented here elucidate the molecular basis of the bivalent recognition of multiply modified histone H3 by BPTF and provide structural insight for developing compounds targeting the PHD finger of BPTF for therapeutic applications.

## Introduction

Epigenetic regulation of gene expression is essential for fundamental biological processes including cellular differentiation, embryological development, and aging.^1,2,3^ In contrast, epigenetic dysfunction underlies many diseases, such as cancer and inflammation.^4,5^ Regulation of the epigenome is mediated by the dynamic installation and removal of diverse chemical modifications of nucleic acid bases and amino acid side chains of histone proteins.^2,3^ Histone proteins are the fundamental building blocks of nucleosomes, which are required for the packaging of DNA into chromatin.^6^ Each nucleosome is constructed of two copies of each of the four core histones: H2A, H2B, H3, and H4, around which ∼147 base pairs of DNA are wrapped.^7^ Histone post translational modifications (PTMs) are dynamically regulated by a coordinated network of epigenetic writer, eraser, and reader proteins that deposit, remove, and interpret these marks, respectively.^8^ The functional readout of histone PTMs by reader proteins, which may contain multiple functional domains, offers both specificity and control for regulating gene expression. The combinatorial readout of histone PTMs and how these recognition events elicit downstream biological processes has been described as a histone code.^9,10^ Elucidating the biomolecular mechanisms behind histone code recognition events by epigenetic reader proteins is essential for understanding molecular mechanisms of gene transcription while also offering new structural insights for developing small molecule therapeutics against epigenetically-driven diseases.^11^ One epigenetic reader protein for which the molecular mechanisms underlying chromatin engagement that has yet to be fully resolved is the Bromodomain and PHD Finger Transcription Factor (BPTF), the largest component of the nucleosome remodeling factor (NURF)^12^.

The BPTF protein contains two adjacent c-terminal chromatin binding domains: a plant homeodomain (PHD) finger and a bromodomain (BRD).^13,14^ Canonically, these domains have been proposed to engage single nucleosomes in a *trans* configuration (Figure **1A**), where the BPTF-BRD engages histone H4 – with a preference for acetylation at lysine 16 (H4K16ac; K_d_ ∼ 100 µM) – while the BPTF-PHD binds to tri-methylated lysine 4 on histone H3 (H3K4me_3_; K_d_ ∼ 3 µM).^15^ Specificity for H4K16ac and H3K4Me_3_ was confirmed using cell-based nucleosomal pull down experiments, with cooperativity between both PTMs driving a 2- to 3-fold enhancement in affinity relative to H3K4Me_3_ alone.^15^ Functionally, BPTF binding to histones directs NURF to specific chromatin loci and enables ATP-dependent nucleosome sliding.^14^ BPTF, and thereby NURF function, is essential for altering gene expression pathways in various biological contexts, including stem cell renewal,^16^ T-cell homeostasis,^17^ and cellular differentiation.^18^

**Figure 1.**
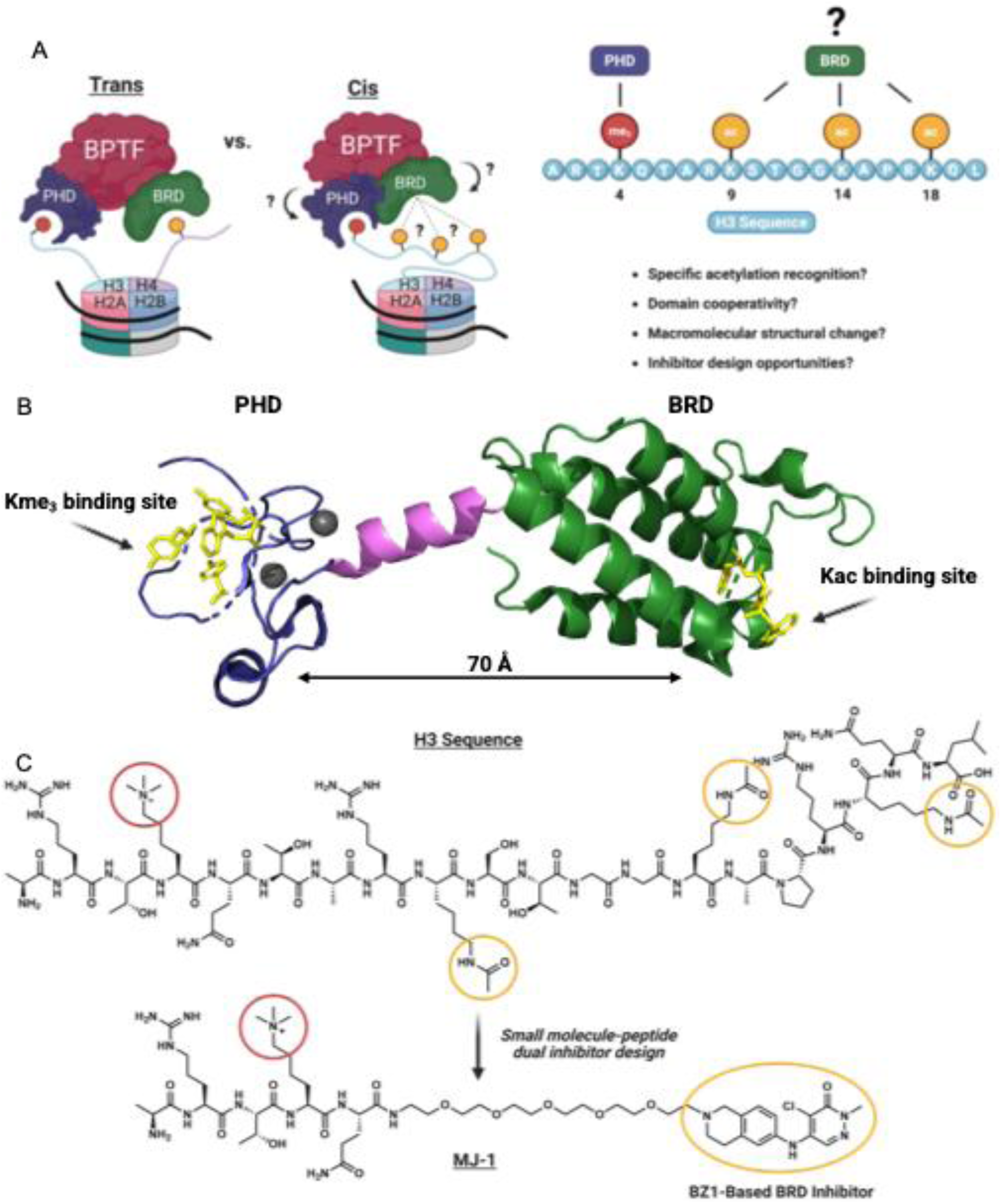
**A)** Comparison of *trans* (left) and *cis* (right) histone engagement mechanisms by BPTF. **B**.) Crystal structure (2F6J) of BPTF-PHD-BRD with chromatin binding domains positioned in orientations not conducive for *cis* engagement. **C**.) Schematic of small-molecule-peptide conjugate design enabled by this work.

BPTF has been identified to have an oncogenic role in many cancers, including triple-negative breast cancer,^19,20^ non-small cell lung cancer,^21,22^ melanoma,^23,24^ neuroblastoma,^25,26^ and colorectal cancer.^27^ Small molecule inhibitors, degraders, and cellular probes targeting BPTF are of interest for further studying the role of BPTF in various disease types. Existing chemical probes targeting BPTF are designed for BRD engagement only, including small molecule inhibitors BZ1,^28^ BZ2,^26^ BI-7190,^29^ and NVS-BPTF-1.^30^ Alternatively, due to their shallow binding pockets, PHDs are a significantly less druggable class of proteins to develop small molecule antagonists against.^31^ The tighter affinity of the BPTF PHD for histones relative to the BRD may support a stronger dependence on PHD function for chromatin binding. As such, PHD inhibition may be an attractive approach considering the absence of significant anticancer activity of bromodomain inhibitors in certain cases.^29,30,32^ We previously reported the first small molecule fragment hits against the BPTF-PHD;^33^ however, attempts to further develop inhibitors from these hits have yet to yield high affinity ligands. Defining the structural basis of the histone interactome of BPTF could enable structure-based drug design efforts for epigenetic readers, like BPTF, that contain poorly ligandable PHDs. Given the multiple epigenetic proteins with adjacent reader domains, such design rules may be more broadly applied.^34^

Although H4K16ac is regarded as the canonical epigenetic mark that the BPTF-BRD recognizes, this reader can interact with additional acetyllysine (Kac) residues on histones. Transient interactions between the BPTF-BRD and the histone H2 variant, H2A.Z, have been characterized previously biophysically and in cell lysates.^35^ Additionally, recognition of histone H3 acetylation by the BPTF-BRD has been proposed. Most recently, Marunde et al.^36^ used peptides derived from the first 20 amino acids of the N-terminus of histone H3 bearing trimethylation at lysine 4 and acetylation at lysines 9, 14, and 18 [H3(1-20)K4me_3_K9acK14acK18ac; see **Table 1**] to test interactions with a BPTF construct containing both PHD and BRD chromatin binding modules (BPTF-PHD-BRD). Peptides engaged the BRD Kac binding pocket by ^1^H,^15^N-HSQC NMR. Peptides and synthetic nucleosomes with H3K4me_3_K9acK14acK18ac modifications were evaluated for direct association with the BPTF-PHD-BRD using an AlphaScreen™-based direct binding interaction assay. No preference for a specific mono-acetylated lysine modification was observed in this assay platform; however, genomic co-localization of BPTF to H3K18ac was observed suggesting this interaction is relevant in a cellular context. Two potential, non-mutually exclusive, explanations have been proposed. First, poly-acetylation is necessary to facilitate PHD engagement of nucleosome-bound histones through neutralizing interactions with DNA;^37,38^ or, second, the BPTF-PHD-BRD engages nucleosomes containing both trimethylation and acetylation on histone H3 simultaneously in a *cis* configuration (**Figure 1A**).^39^ In a separate study, Chen et al.^40^ characterized peptides representative of the first 17 amino acids of the N-terminus of histone H3 to suggest that K14ac on histone H3 can interact with the BRD in vitro and subsequently in cells. However, a direct interaction with K14ac alone was not observed by isothermal titration calorimetry (ITC) and the number of binding sites was ambiguous between a 1:1 (*cis*) binding event versus a 1:2 (*trans*) binding event. Given the short length of peptides used in this study, the contributions of K18ac were not evaluated.

**Table 1.**
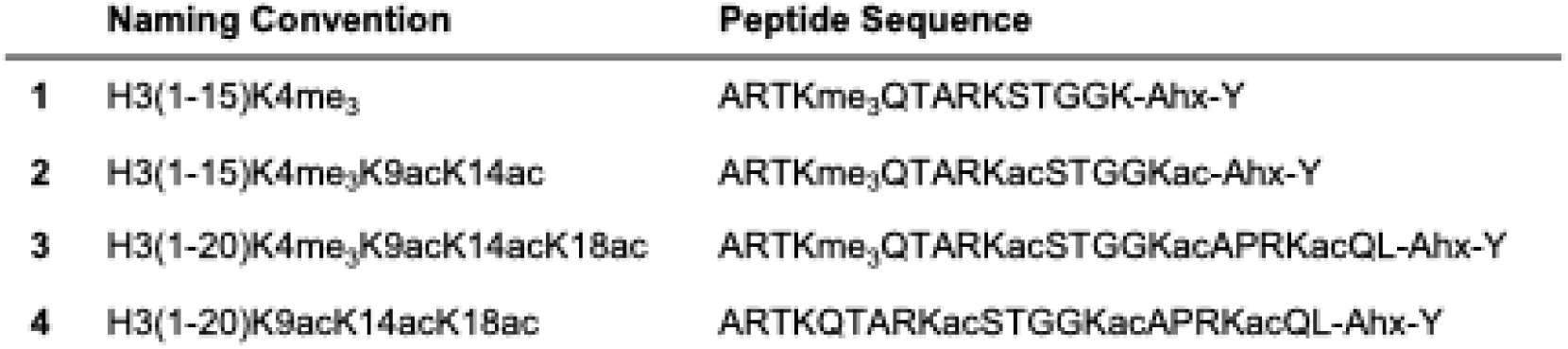
Histone H3 peptide ligands synthesized based on literature precedence (me_3_ = trimethylation; ac = acetylation; Ahx = amino hexanoic acid).

|  | Naming Convention | Peptide Sequence |
| --- | --- | --- |
| **1** | H3(1-15)K4me<sub>3</sub> | ARTKme<sub>3</sub>QTARKSTGGK-Ahx-Y |
| **2** | H3(1-15)K4me<sub>3</sub>K9acK14ac | ARTKme<sub>3</sub>QTARKacSTGGKac-Ahx-Y |
| **3** | H3(1-20)K4me<sub>3</sub>K9acK14acK18ac | ARTKme<sub>3</sub>QTARKacSTGGKacAPRKacQL-Ahx-Y |
| **4** | H3(1-20)K9acK14acK18ac | ARTKQTARKacSTGGKacAPRKacQL-Ahx-Y |

Taken together, these two studies suggest that H3 peptides containing lysine trimethylation and acetylation PTMs can engage both domains of the BPTF-PHD-BRD module in a nucleosomal context. However, the degree of histone H3 Kac necessary for BRD engagement, or the presence of a single predominant Kac binding motif, remain to be elucidated. Additionally, whether both chromatin binding domains of BPTF engage a multiply modified peptide simultaneously in *cis* needs to be verified biophysically. The structural basis of a *cis* combinatorial PTM H3 recognition event by BPTF-PHD-BRD is unknown since existing crystal structures have only resolved ligands in either the PHD or BRD domains, but never both simultaneously.^15,41^ In crystallized structures, the positioning of the reader domains appears incompatible for intramolecular *cis* engagement of methylated and acetylated histone H3 peptides due to a linear orientation and ∼70 Å distance^15^ between the two binding sites (**Figure 1B**). To resolve this structural disconnect, here we used differential scanning fluorimetry (DSF), competition-based AlphaScreen™ assays, and ITC to monitor the binding contributions of each reader domain of BPTF-PHD-BRD to a library of peptides that vary in length, Kac position, and degree of lysine acetylation/methylation. We used inactive domain mutants to confirm the binding location of these H3 peptide ligands within the PHD and BRD binding pockets and obtain the first crystal structure of the BPTF-PHD-BRD bound to a trimethylated and acetylated histone H3 peptide (2.17 Å), along with a shorter PHD-only binding peptide for reference. Finally, we employed size exclusion chromatography coupled small angle X-ray scattering (SEC-SAXS) to verify the oligomeric state and survey the conformational populations of BPTF-PHD-BRD when bound to dually modified H3 peptides, with additional macromolecular structural analyses performed using limited proteolysis mass spectrometry (LiP-MS). We find that BRD engagement is predominated by a single Kac modification at H3K14ac with improved affinity over H4K16ac, and that this PTM is read cooperatively with H3K4me_3_ by the PHD. Our data support that at the peptide level, combinatorial reading by BPTF-PHD-BRD of PTMs on H3 peptides can proceed via a *cis* binding event in addition to the previously reported *trans* engagement, and that this mode of bivalent molecular recognition can be accessed through conformational rearrangement of the domains in solution. These results provide a basis for understanding the biophysical mechanisms behind the cooperative reading ability of BPTF’s tandem chromatin binding domains. We culminate our investigation into this combinatorial mechanism of interaction by reporting the first examples of dual engaging inhibitors of the BPTF-PHD-BRD (**Figure 1C**). These results expand the H3 histone code for BPTF engagement to chromatin and establish a foundation for developing inhibitors of PHD finger-containing proteins. This work further supports prior efforts demonstrating that the BPTF BRD can recognize H3Kac epitopes in addition to H4K16ac. These findings also help resolve a structural conundrum for dual domain engagement to histone H3 by BPTF based on the previously available structural information and provide design rules for bivalent inhibitors that could overcome the longstanding ligandability challenge of PHD finger-containing proteins.

## Results and Discussion

### Combining H3 Methylation and Acetylation Enhances Binding Interactions

To assess the influence of H3 trimethylation and acetylation alone versus when combined, we synthesized a set of four peptides that varied in length, Kme_3_, and Kac inspired by literature precedence. Peptides containing the first 15 amino acids of the N-terminal histone H3 sequence have been used previously to analyze BPTF-PHD finger engagement to K4me_3_ (**1** and **2, Table 1**).^41,42^ Longer peptides **3** and **4** containing another acetylation site at K18 were also synthesized^36^ and the extent of thermal stabilization of BPTF-PHD-BRD by each ligand was evaluated by DSF. DSF relies on a fluorescent reporter dye to monitor temperature-dependent unfolding of proteins. Ligand binding often stabilizes proteins, enabling measurement of the change in melting temperature (ΔT_m_) between bound and unbound states serving as a proxy for binding affinity.^43^ Addition of **1** or **4** resulted in ΔT_m_ values of approximately +2 °C (**Figure 2A**). When these PTM patterns were combined in **3**, a greater than a 3-fold increase in ΔT_m_ (+7.9 °C, **Figure 2A**) was observed. **2** stabilized BPTF-PHD-BRD to a greater extent than peptides **1** and **4** but not to the extent of **3**, suggesting that longer peptides that include lysine 18 may exhibit enhanced binding interactions when compared to shorter counterparts.

**Figure 2.**
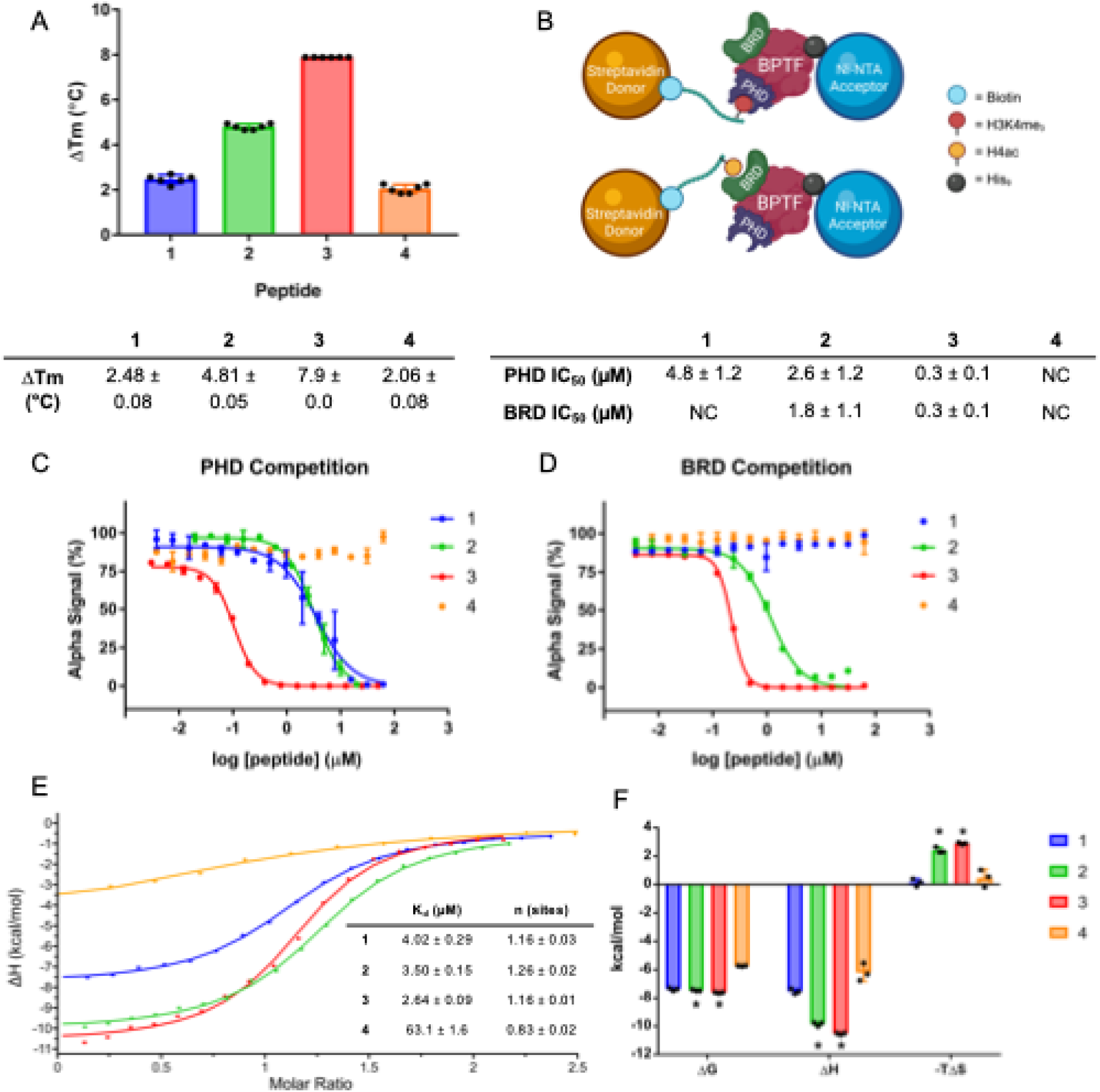
Biophysical Characterization of BPTF-PHD-BRD Histone Interactions. **A**.) Change in melting temperature (ΔT_m_) with peptides bound relative to apo BPTF. Average melting temperature derived from six technical replicates ± standard error of the mean (SEM). **B**.) Overview of Alphascreen™ competitive inhibition platforms used in this study and average IC_50_ values calculated from at least three experimental replicates ± SEM. NC = no competition, defined by <50% maximal inhibition. **C** and **D**.) IC_50_ curves, with each point reflecting the average of two technical replicates. **E**.) ITC binding isotherms with average ± SEM of binding affinities and n values for peptides 1-4. **F**.) Thermodynamic parameters for peptides 1-4, all derived from three experimental replicates. Asterisks = peptides with combined K4me_3_ and K14ac PTMs.

We also used AlphaScreen™, a platform that can detect low-affinity interactions due to avidity^44^ and is commonly used in inhibitor discovery campaigns and PPI profiling.^45,46^ A direct-binding AlphaScreen™ assay has been previously used with BPTF-PHD-BRD to quantify interactions with peptides and nucleosomes.^36^ While apparent affinities can be determined for relative rank ordering of binding interactions, we reasoned that this direct-binding assay might obscure subtle structure-activity relationships among peptide ligands bearing combinations of Kme_3_ and Kac PTMs. Therefore, we employed competition-based Alphascreen™ assays by using biotinylated histone peptides that bind to either the PHD or BRD (**Figure 2B**), which are routinely used for generating domain-specific IC_50_ values with competing small molecule ligands.^28,33^ We observed an IC_50_ of 4.8 µM for **1** in a PHD competition experiment with the identical biotinylated H3(1-15)K4me_3_ probe, which is comparable to the reported binding affinities of 2-8 µM for this interaction (**Figure 2B and 2C**).^41,42^ **3** exhibited an ∼16-fold reduction in PHD IC_50_ relative to **1**, suggesting that distal Kac modifications at lysines 9, 14, or 18 could form interactions with the BPTF-PHD-BRD that drive the observed potency increase. A submicromolar BRD IC_50_ of 0.3 µM of was observed for **3** (**Figure 2B and 2D)**, which suggested that H3Kac interactions occur within the BRD Kac binding pocket. These increased affinities were not as pronounced with **2** (**Figures 2B-2D**) which aligned with trends observed in DSF and highlighted the importance of peptide length, K18ac presence, or a combination of both properties. Finally, we found that enhancement of both ΔT_m_ and BRD IC_50_ was dependent on the PHD-K4me_3_ interaction as **4** elicited weaker thermal shift values than **2** or **3. 4** was also unable to outcompete the tetra acetylated H4 probe from the BRD in AlphaScreen™ (**Figure 2A-D)**, implying that higher concentrations of the peptide may be needed to observe full competition. Together, these data suggest that H3K4me_3_ and H3Kac are read cooperatively and PHD engagement is required for enhanced potency.

We next turned to ITC for generating K_d_ values, thermodynamic parameters (ΔG, ΔH, and ΔS), and binding stoichiometries (n) as it is routinely used to monitor the binding preferences of histone reader proteins.^42,47,48,49^ The stoichiometry and K_d_ of binding for peptide **1** mirrored previous measurements obtained for the PHD (**Figure 2E and 2F**).^42^ Thermodynamic analyses revealed **2** and **3** exhibited acetylation-dependent increased enthalpic responses relative to peptide **1**. However, enhanced enthalpic binding contributions for **2** and **3** were accompanied by a significant entropic penalty, which resulted in similar binding affinities and free energies of binding among **1-3**. All peptides demonstrated near 1:1 binding stoichiometry and the observed enthalpy-entropy compensation was specific to peptides containing combined K4me_3_ and Kac PTMs. Therefore, we reasoned that simultaneous engagement of both the PHD and BRD is enthalpically-driven, yet entropically unfavorable. Prior to elucidating the biomolecular mechanism behind this entropic penalty of binding, we focused our attention on determining if tri-acetylation was necessary for observing enhanced thermal stabilization, reduced IC_50_ values, and larger enthalpies of binding.

### Peptide Structure Activity Relationships Uncover a Predominating Kac Motif

We next sought to determine if a specific pattern of mono- or di-acetylation on the H3(1-20) sequence may predominate BRD interactions given that the BPTF BRD exhibits specificity for mono-acetylated histone H4. To test his hypothesis, we synthesized peptides (**Table 2**) that were 20 amino acids in length to ensure that structure-activity relationship (SAR) trends (**Figure 3**) were dependent on changes in PTM patterns and not peptide length. **5-10** all contained K4me_3_ in combination with di- and mono-Kac sequences. Di-acetylation (**6** and **7**, respectively) showed enhanced ΔT_m_ measurements and reduced IC_50_ values relative to unacetylated **5**, with these trends being slightly more pronounced for **7**. **8** maintained similar thermal stabilization and PHD IC_50_ values to **5** and did not demonstrate competitive binding in the BRD platform, suggesting that acetylation at lysine 9 (K9ac) alone is not sufficient for mediating interactions with the BRD. **9** – containing mono-acetylation at lysine 14 (K14ac) – resulted in a ΔT_m_ that exceeded +10 °C and demonstrated sub-micromolar IC_50_ values in both competition platforms. Although mono-acetylation at lysine 18 (K18ac; **10**) corresponded to a moderately increased ΔT_m_ compared to **5**, inhibition was not observed in the BRD competition platform and the PHD IC_50_ was still in the low-micromolar range. These data suggested that K14ac is the predominant acetylation mark behind the potency improvements observed in DSF and AlphaScreen™ for multiply acetylated ligands. Additionally, these trends were dependent on PHD functionality as peptide **11**, bearing K14ac without K4me_3_, did not induce significant thermal stabilization or elicit dose-dependent inhibition, thereby corroborating our previous findings with **4** and supporting the concept of cooperative binding between the PHD and BRD.

**Figure 3.**
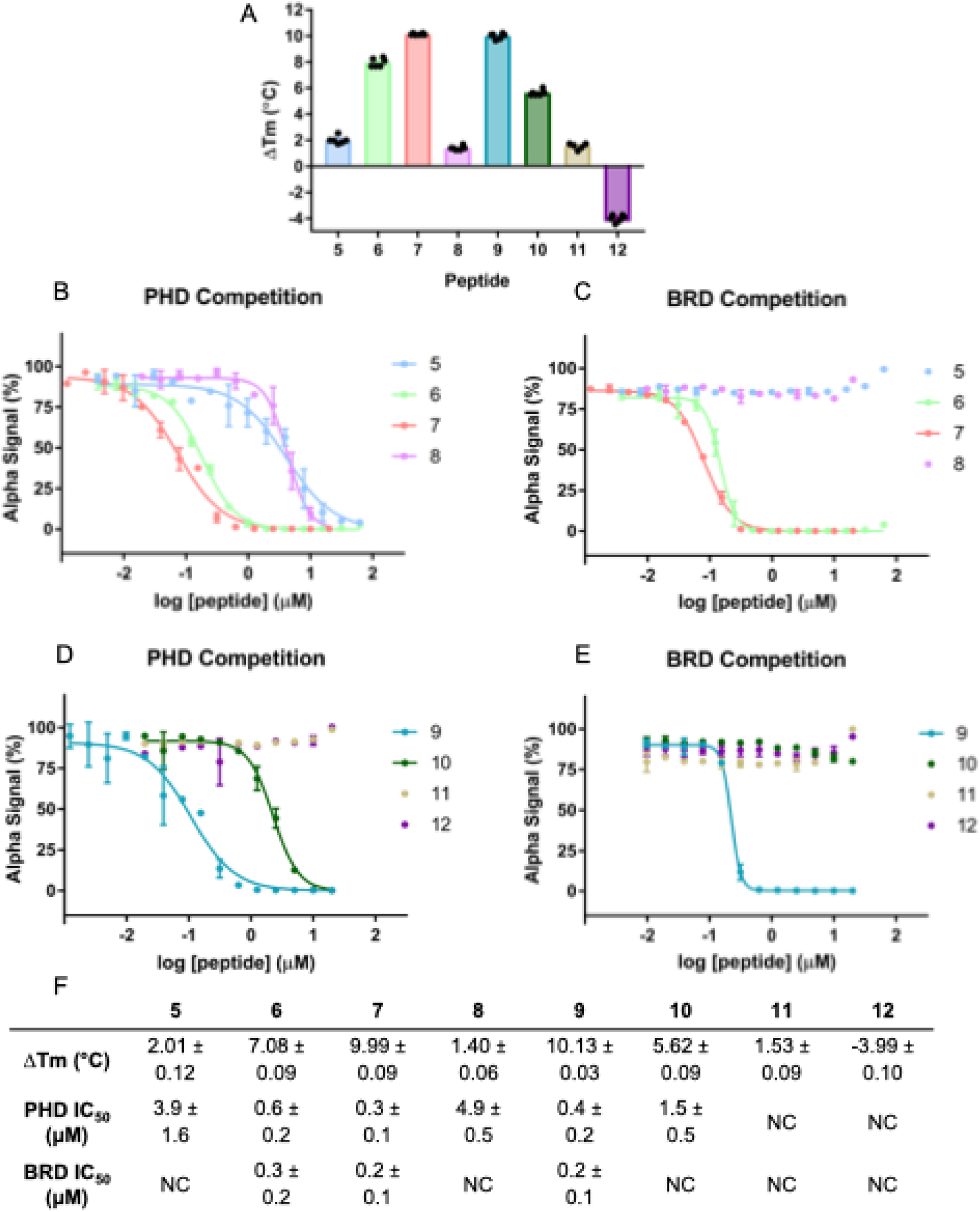
Biophysical Characterization of the BPTF-PHD-BRD with Histone Peptides 5-12. **A**.) DSF ΔT_m_ values for peptides **5**-**12**. **B-E**.) AlphaScreen™ IC_50_ curves, with each point reflecting the average of two technical replicates. Average ± SEM shown in **F**.

**Table 2.** Histone H3 ligands containing 20 amino acids with varying PTMs.

|  | Naming Convention | Peptide Sequence |
| --- | --- | --- |
| **5** | H3(1-20)K4me<sub>3</sub> | ARTKme<sub>3</sub>QTARKSTGGKAPRKQL-Ahx-Y |
| **6** | H3(1-20)K4me<sub>3</sub>K9acK14ac | ARTKme<sub>3</sub>QTARKacSTGGKacAPRKQL-Ahx-Y |
| **7** | H3(1-20)K4me<sub>3</sub>K14acK18ac | ARTKme<sub>3</sub>QTARKSTGGKacAPRKacQL-Ahx-Y |
| **8** | H3(1-20)K4me<sub>3</sub>K9ac | ARTKme<sub>3</sub>QTARKacSTGGKAPRKQL-Ahx-Y |
| **9** | H3(1-20)K4me<sub>3</sub>K14ac | ARTKme<sub>3</sub>QTARKSTGGKacAPRKQL-Ahx-Y |
| **10** | H3(1-20)K4me<sub>3</sub>K18ac | ARTKme<sub>3</sub>QTARKSTGGKAPRKacQL-Ahx-Y |
| **11** | H3(1-20)K14ac | ARTKQTARKSTGGKacAPRKQL-Ahx-Y |
| **12** | H3(1-20)Unmodified | ARTKQTARKSTGGKAPRKQL-Ahx-Y |

ITC analyses of **5-12** revealed that, generally, peptides containing combined K14ac and K4me_3_ PTMs displayed the largest enthalpic contributions of binding with compensatory entropic penalties. These results aligned with our findings for **2** and **3** and further highlight K14ac as the driving force behind enthalpic gains and a mediator of BRD interactions (**Figure 4**). Although **6** is the exception to K14ac-mediated enthalpic gains, this result aligns with our earlier findings of pronounced ΔT_m_ measurements and slightly enhanced IC_50_ values for **7** and **9** that lack K9ac relative to **6**, suggesting that the presence of K9ac within the histone H3 sequence may have a minor negative effect on the BRD’s ability to recognize neighboring K14ac. This phenomenon has been identified in other bromodomains.^50^ Alternatively, K9ac was recently shown to engage an allosteric region of BPTF PHD, which may explain the unique binding characteristics observed for this ligand.^51^ Although longer peptides displayed tighter binding affinities than their shorter counterparts, minimal differences in binding affinity were observed among **5-10** due to enthalpy-entropy compensation, which is in line with our ITC findings for **3**. As shown with **11**, H3K14ac alone exhibited a K_d_ of 50.9 µM for the BRD, which is stronger than the reported affinities for H4K16ac and greater than 20-fold more potent than mono-acetylation on histone H2A.Z.^15,52^ No direct interaction was observed by unmodified **12** in ITC (see SI), suggesting that the destabilization event observed for this ligand in DSF (**Figure 3A and 3F)** may be due to transient interactions.

**Figure 4.**
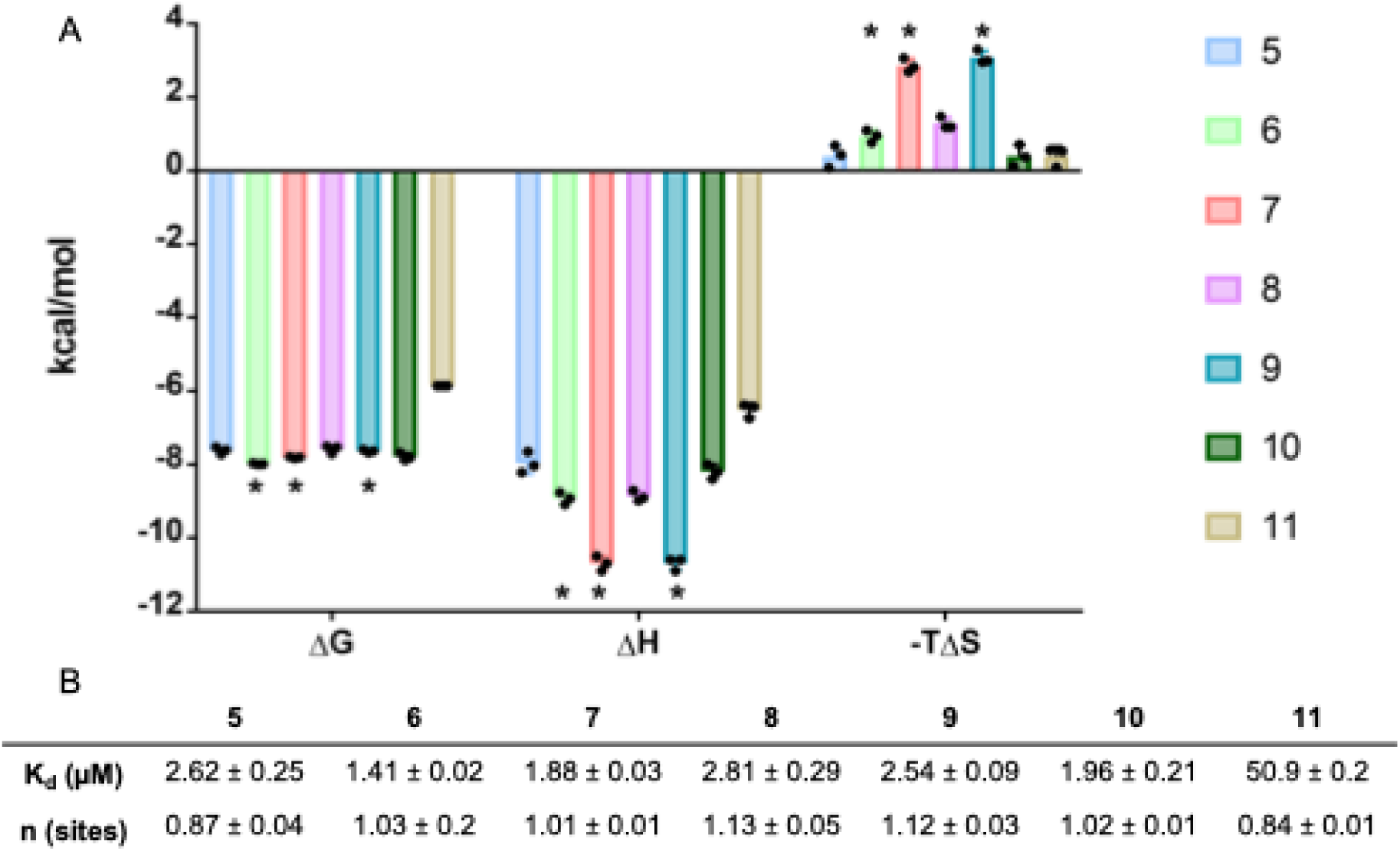
ITC Analysis of Peptides 5-11. **A**.) Thermodynamic parameters for peptides 5-11. **B**.) Associated binding affinities and n-values with peptide **12** omitted due to lack of interaction. Asterisks = peptides with combined K4me_3_ and K14ac PTMs.

### Mutagenesis Studies Confirm Binding Sites

Next, we set out to confirm the binding location of K4me_3_ and K14ac side chains to the PHD and BRD. We expressed BPTF-PHD-BRD mutant constructs harboring single amino acid substitutions previously demonstrated to eliminate binding ability to either Kme_3_ (PHD_mut_, W32E) or Kac (BRD_mut_, F154A).^15,41^ As expected, the PHD_mut_ and BRD_mut_ constructs were incapable of mediating ternary complex formation with biotinylated probes, respectively, confirming domain-specific inactivity (**Supplementary Figure S1**). Importantly, the unmutated domain within each construct maintained functionality in our AlphaScreen™ platforms, confirming that these domain-specific mutants can be used to further interrogate the contributions of individual reader modules within the PHD-BRD protein (**Supplementary Figure S1**). Peptides containing K4me_3_ and K14ac modifications were selected for evaluation in DSF and AlphaScreen™ assays with each mutant protein. These peptides elicited significantly smaller stabilization values relative to the wild-type (WT) BPTF-PHD-BRD, corresponding to at least four-fold reductions in ΔT_m_ (**Figure 5A and 5B**). ΔT_m_ measurements of approximately +2 degrees were observed with the BRD_mut_, which was consistent with PHD engagement alone. However, thermal stabilization events with the PHD_mut_ were negligible. These trends were recapitulated in AlphaScreen™ BRD competition experiments where the PHD_mut_ showed no binding for any of the tested peptides (**Figure 5B and 5D**). The absence of binding with the PHD_mut_ in both platforms emphasizes the reliance of PHD engagement for peptide interactions. Similarly, AlphaScreen™ PHD competition experiments with the BRD_mut_ demonstrated significantly weaker binding (single digit micromolar) for the tested peptides compared to WT BPTF (**Figure 5B and 5C**), further underscoring the necessity of both domains for a cooperative binding event to occur. Together, these data provide direct evidence for molecular recognition of K4me_3_ within the aromatic cage of the PHD and K14ac within the canonical BRD Kac binding pocket. Further, these mutational analyses support a cooperative binding mode between both BPTF chromatin binding domains when reading H3 peptide ligands. Importantly, with this AlphaScreen™ platform we cannot rule out affinity gains being driven by intermolecular peptide interactions between different protein units. Additionally, the multi-valent display may reduce the entropic penalty observed by ITC.^53^

**Figure 5.**
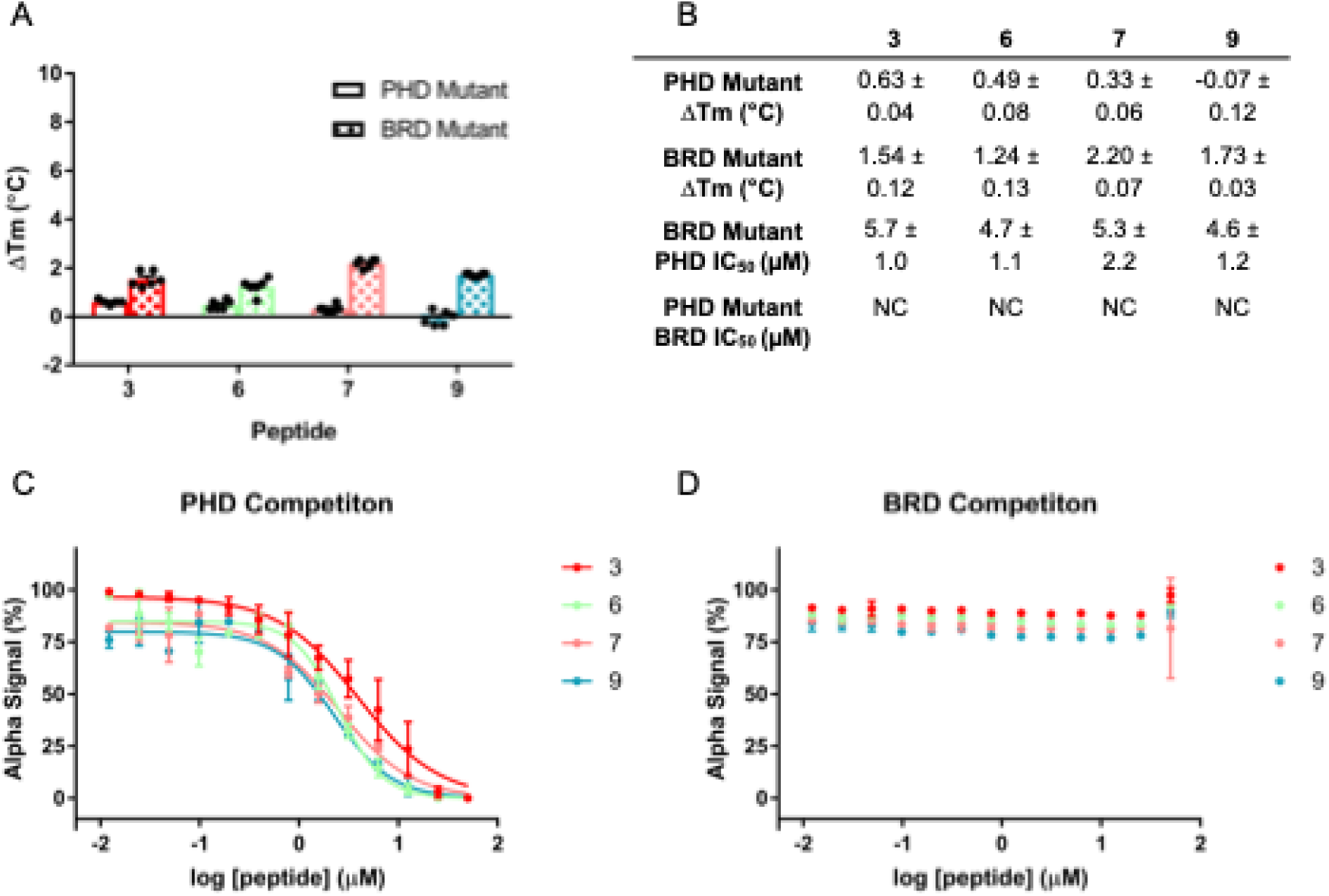
Binding Analysis with BPTF Single-domain Mutants. DSF ΔT_m_ (**A**) and AlphaScreen™ competition plots (**C** and **D**). Average ± SEM shown in **B**.

### Crystallographic Studies Confirm Binding Sites

To further validate the binding location of Kme_3_ and K14ac containing peptides to BPTF-PHD-BRD and visualize the structural basis behind the cooperative binding event, we performed co-crystallization studies. Given that existing crystal structures show that the BPTF-PHD-BRD peptide binding pockets orient away from each other in a linear fashion that spans distances of ∼70 Å,^15^ we expected a conformational change placing the domains in closer proximity to each other was necessary for a bivalent binding mechanism. Such a mechanism is plausible based upon previously conducted molecular dynamics (MD) simulations with BPTF-PHD-BRD bound to peptides and nucleosomes containing K4me_3_ and would explain the enthalpy-entropy compensation observed for dually binding peptides in ITC.^54,55^ To resolve both the binding mode and interactions with the PHD and BRD pocket, we solved the 2.17 Å co-crystal structure of BPTF-PHD-BRD bound to peptide **3**. Under crystallography conditions, our co-crystal structure supports an intermolecular dimerization event whereby one molecule of **3** bridges across two BPTF-PHD-BRD monomers and forming dimers in a 2:2 head-to-tail configuration (**Figure 6A and Supplementary Figure S2A**). To further clarify the observation of the crystallographic dimer, we solved another structure bound with a seven residue H3 peptide which has dimethyl group at 4^th^ position of lysine and lacks any acetylation site that could interact with the BRD. The electron density allowed modeling of the full peptide sequence in the crystal structure (**Supplementary Figure S2B**). To elucidate potential mechanisms driving dimerization, we constructed a dimer based on crystallographic symmetry operations of the monomer in the asymmetric unit. As the head-to-head packing is different from the peptide **3** dimer and other structures, e.g. apo, this argues for the stabilization of a peptide-specific energetically favorable state by **3** that gets preferentially packed during crystal growth (**Supplementary Figure S2**).

**Figure 6:**
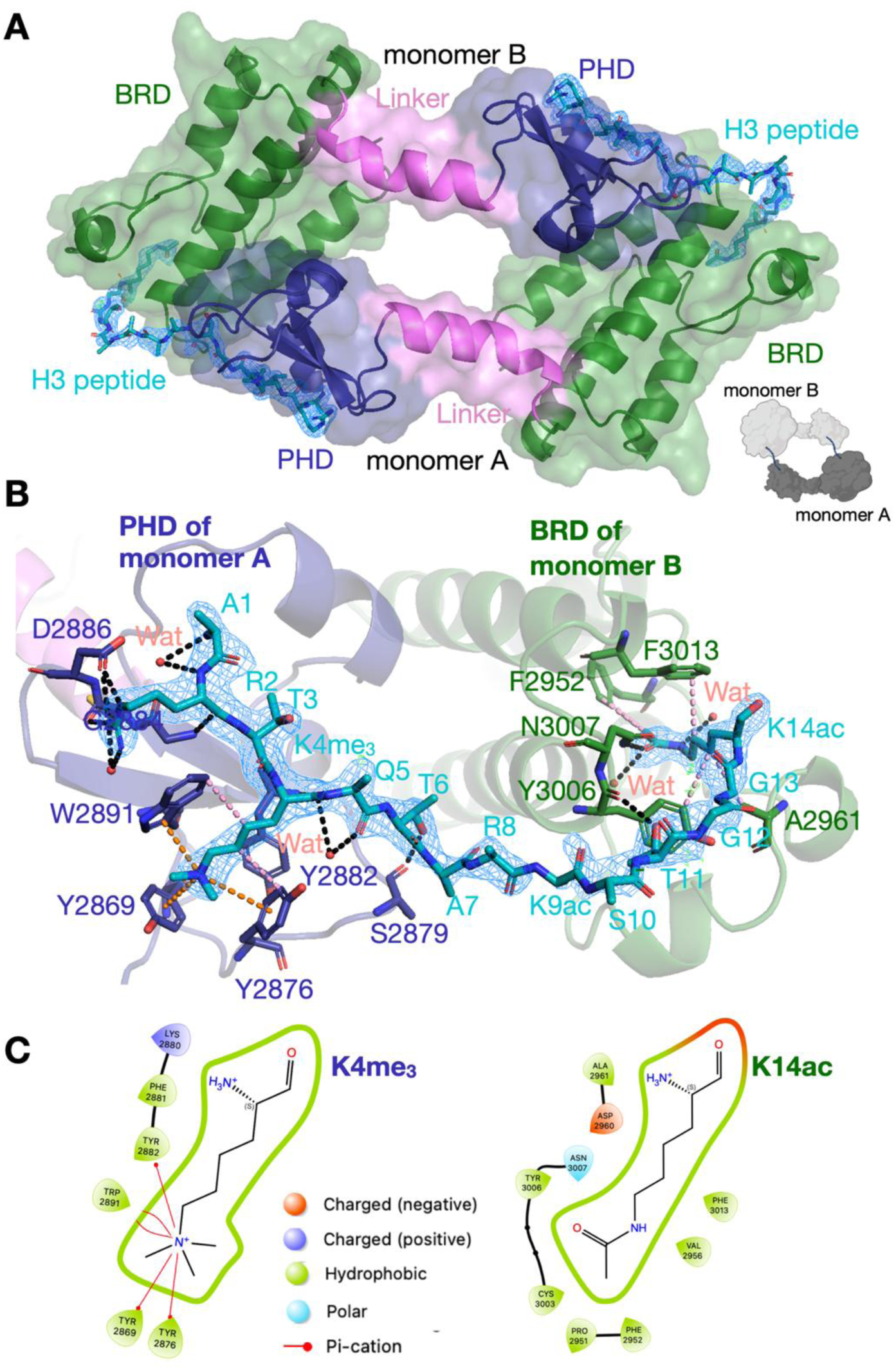
Crystal structure of BPTF-PHD-BRD with 3 reveals binding site and mode. (**A**) The symmetrical 2:2 head-to-tail dimer (see inset) of **3** bound BPTF-PHD-BRD is shown in surface and cartoon representation. 2Fo-Fc maps are shown in blue at 1σ and Fo-Fc maps are shown in green and red at 3σ and −3σ, respectively. Inset is a pictorial representation of BPTF-PHD-BRD-**3** complex in the head-to-tail conformation. (**B**) Bridging interactions of **3** with PHD and BRD domains; 2Fo-Fc and Fo-Fc maps are rendered as same as in panel **A**. Pink lines are hydrophobic interactions. Orange lines are cation-*π* interactions. Black lines are polar interactions. (**C**) Interaction profile of K4me3 and K14ac at the PHD and BRD pockets, generated using Maestro software (Schrödinger, New York).

Another insight we gained from the structure was how the trimethylated lysine binds to the PHD pocket of one monomer, and which acetylated lysine binds to the BRD pocket of another monomer (**Figure 6**). Prominent electron density for the peptide from A1 to K14ac (**Figure 6**) revealed that the PHD pocked binds K4me_3_ via a collection of cation-*π* and van-der-Waals interactions between the trimethyl ammonium group and aromatic cage containing three tyrosine residues and a tryptophan, as seen in other co-crystal structures.^41^ Additional interactions with the PHD pocket are formed by proximal peptide residues A1 and R2 supporting the binding data, showing high affinity binding of **3** to the PHD domain (**Figure 6**). Of the three acetylated lysines, the BRD pocket unambiguously binds K14Ac, forming hydrophobic and water-mediated interactions (**Figure 6**) similar to those observed in other published structures.^15^ The K14Ac binding mode confirms AlphaScreen™, DSF, and ITC experiments that showed that K14ac predominantly drives peptide binding to the BRD pocket over K9ac and K18ac (**Figure 6B**). Except for K14ac and T11, no major BRD engagement is observed, aligning with the loss of binding affinity observed with mono-acetylated **11** that does not contain trimethylated lysine (**Figure 6B**). Electron density for the middle portion of the peptide, from A7 to S10, only traces the peptide backbone, in agreement with a flexible, solvent-exposed region that contributes insignificantly to binding. Notably, this includes acetylated lysine at the 9^th^ position; also, no amino acid side chain after K14ac (A15 to L20) could be resolved in the structure, including K18Ac (**Figure 6B**).

### Dual Engaging Peptides Induce Conformational Changes in Solution

To delineate whether this novel mode of dimerization-based recognition could be observed in other approaches, we turned to solution-based techniques to characterize these protein-peptide complexes. SEC-SAXS is a technique used to study the size, structure, and oligomeric state of biomolecules in solution.^56,57,58^ SEC-SAXS has been used previously on other epigenetic tandem readers to glean insight on domain positioning and the overall molecular shape adopted in solution.^59,60^ We obtained scattering data on the BPTF-PHD-BRD protein alone and in the presence of **3** to assess potential changes in protein structure upon simultaneous engagement of both reader domains (**Figure 7B**). We focused our analysis on the major species present in SEC. Molecular weight calculations resulted in values within 3 kDa of the expected mass of a BPTF monomer (∼22.8 kDa; **Figure 7A**), corresponding to the major species in solution. The atom pair distance distribution (P(*r*)) function provides visualization of distances between all pairs of atoms within the particle.^61^ Globular macromolecules exhibit a Gaussian curve whereas longer, extended structures exhibit asymmetry and peak tailing.^61^ BPTF-PHD-BRD displayed a near-symmetrical curve centered at an interatomic distance (*r*) of approximately 22 Å (**Figure 7A and 7D**) when bound to **3**. In comparison, the P(*r*) function without **3** exhibited a greater degree of asymmetry and peak tailing toward larger *r* values, suggesting that BPTF-PHD-BRD adopts extended conformations in the unbound form (**Figure 7D**). The Kratky plot is another qualitative representation that captures structural globularity and flexibility in biomolecules. Extended structures produce larger scattering intensities that plateau at increased scattering angles (*q*), whereas compact species form ordered, bell-shaped curves.^61^ Both peptide-bound and -unbound samples showed a mixture of compact and elongated species, represented by the presence of both Gaussian distributions and increased disorder at larger *q* values (**Figure 7C**). Interestingly, the peptide-bound sample demonstrated a leftward shift with the curve peaking closer to qR_g_ ∼ 1.73, where the curve of a Kratky plot for an ordered, globular protein should reach a maximum value.^62^ These data suggest that simultaneous engagement of both domains induces a more globular structure relative to the unbound form (**Figure 7C**). This data aligns with the approximate 3 Å reduction in R_g_ we observed in the presence of **3** (ΔR_g_ = −3.0 Å for P(*r*) function derivation; ΔR_g_ = −3.3 Å for Guinier approximation; **Figure 7A**). We thus infer that dually engaging peptides induce a conformational change that orients the PHD and BRD into a more compact, globular configuration. Our SAXS analyses are in good agreement with findings from a MD study that calculated a R_g_ of 25.0 Å for the extended form of BPTF-PHD-BRD, coupled with a ΔR_g_ of −3.1 Å when BPTF-PHD-BRD was bound to a nucleosome core particle.^54^

**Figure 7.**
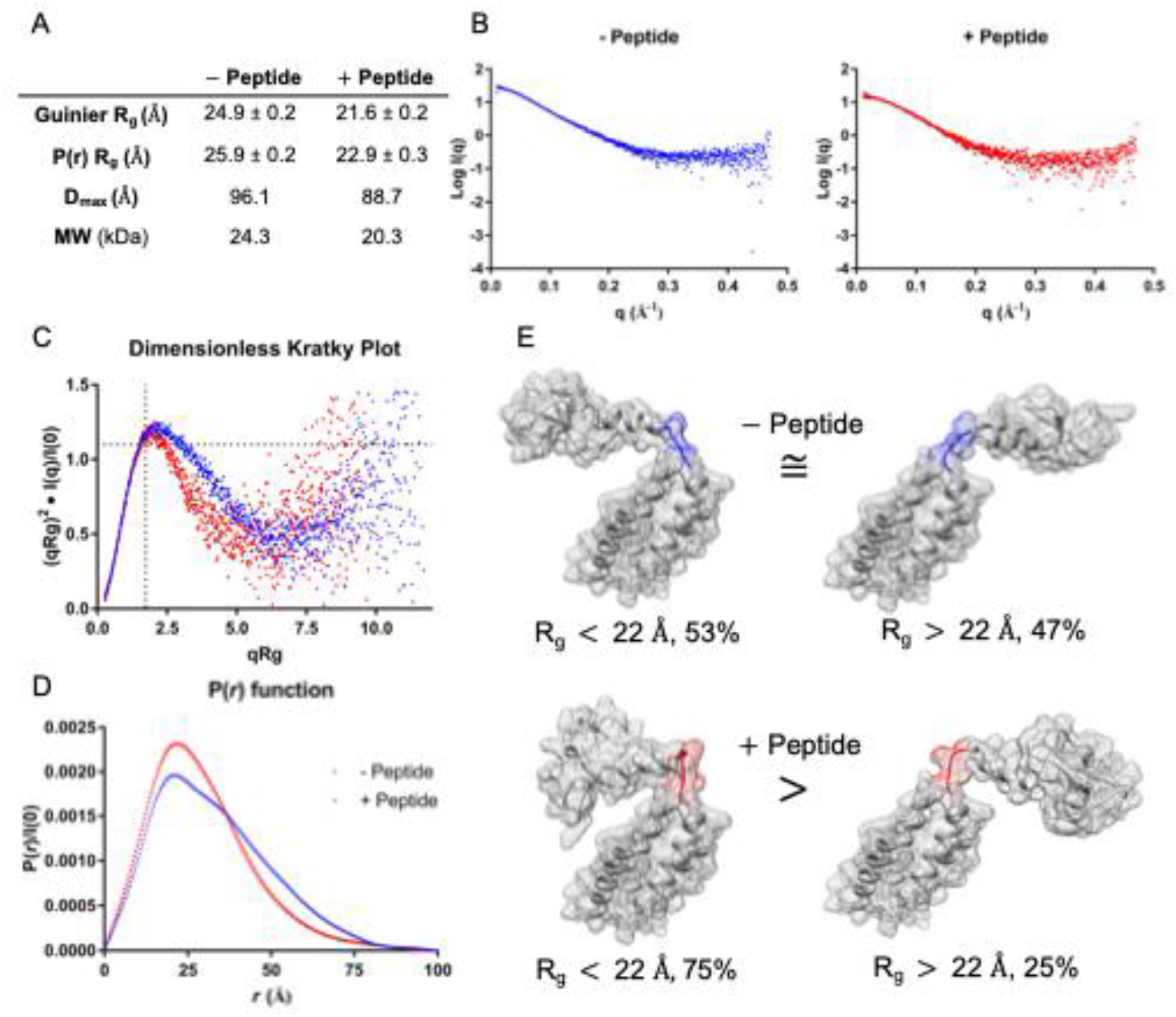
SEC-SAXS Data Analysis of the BPTF-PHD-BRD with and without 3. **A**.) Summary R_g_, Dmax and molecular weight values obtained from SAXS. **B**.) Scattering profiles obtained for unliganded BPTF-PHD-BRD (left, blue) and bound to peptide **3** (right, red). **C** and **D**.) Kratky and P(*r*) representations. **E**.) Representative poses of the multiple conformational populations modeled by MultiFoXS from scattering data between unliganded (top) and bound to **3** (bottom) for the 4-state model. Hinge residues are highlighted in each pose. Poses are shown for populations occupying 20.7 Å and 26.3 Å (unliganded) and 18.2 Å and 28.4 Å (**3** bound), with cumulative population weighted ranges surrounding a 22 Å cut off annotated. Detailed R_g_ distributions for both fitting models can be found in **supplementary Table 3**.

We used the Multi Fast X-ray Scattering (MultiFoXS) server to model the conformational change indicated by SAXS. MultiFoXS can model conformationally heterogeneous samples and estimate the percentage of each conformer that is present by merging experimental SAXS data with a crystal structure.^63,64^ Amino acids 65 through 69 (V65-L69; 2F6J) were selected as flexible residues based upon previous MD simulations that identified this region as a molecular hinge that permits interdomain flexibility.^54^ Both 4-state models generated acceptable *χ* values of 1.57 and 0.85 for the peptide absent and peptide present conditions, respectively.^63^ Modeling BPTF-PHD-BRD without **3** bound showed that the R_g_ distribution split into two distinct, yet roughly equally weighted ranges: 53% of the population adopted radii between 19.2-20.7 Å, and the remaining 47% fell between 24.3-26.2 Å (**Figure 7E** and **Supplementary Table 3**). Once bound to **3**, the R_g_ distribution favored smaller radii. 75% of the population was narrowly centered between 18.2-19.0 Å, whereas the remaining 25% had radii between 24.0-28.4 Å (**Figure 7E** and **Supplementary Table 3**). Although the output structures generated from MultiFoXS can assist in visualization of multiple conformational states and hypothesize potential protein structures, these models have limited applicability to residue-specific analyses of the PHD finger as structural zinc ions are not included in the modeling process.

Following SEC-SAXS, we used limited proteolysis mass spectrometry (LiP-MS) to characterize structural changes at the amino acid level to BPTF-PHD-BRD upon **3** binding. LiP-MS is a solution-based proteomics technique that monitors ligand-induced conformational changes to proteins.^65,66,67^ LiP-MS relies on the use of nonspecific proteases (such as proteinase K) that cleave at solvent-exposed regions of folded proteins, rather than specific amino acids, resulting in the production of conformotypic peptides that change in abundance between ligand-bound and - unbound conditions.^66,68^ Proteolysis was conducted on BPTF-PHD-BRD in the absence of ligand or while bound to **3**. LiP-MS analysis uncovered seven conformotypic peptides that were differentially regulated in the presence of **3** (**Supplementary Figure S7A**). Structural mapping of the two conformotypic peptides that decreased in abundance revealed proteolytic cleavage sites on an alpha helix of the BRD that are adjacent to the acetyllysine binding channel in both a crystal structure and the model produced by MultiFoXS (**Supplementary Figure S7B and S7C**, respectively). The remaining conformotypic peptides increased in abundance and corresponded to cleavage sites on the PHD Finger, a disordered region of the BRD, and the alpha helical linker between both domains. Thus, we conclude that binding of **3** shields structured regions of the BRD from proteolytic cleavage while simultaneously promoting solvent exposure of multiple areas across the entire protein structure. Indeed, these unique proteolytic cleavage profiles may correspond to parts of the protein that make up external and internal faces of the globular peptide-protein complex observed in SAXS.

To better understand if structural flexibility can be represented in static crystal structures, we superposed the BPTF-PHD-BRD bound to **3** and a shorter H3(1-7)K4me_2_ peptide and compared them with published structures (**Supplementary Figure S3**). Superposing the PHD region of peptide **3** with the PHD region of the short peptide bound structures with an RMSD of 0.46 Å leads to a conformational shift of approximately 10 Å in the BRD region, bringing the BRD little closer to the PHD domain (**Supplementary Figure S3A**). Additional comparisons with deposited structures yielded different degrees of structural flexibility between 3 to 10Å, depending on the bound-state (**Supplementary Figure S3B-3E**). This highlights the context-dependent flexibility of domain positioning that may explain results observed in the solution-based measurements.

Taken together, the marked reduction in R_g_ observed upon engagement by **3** in SAXS and the population distribution modeling by MultiFoXS corroborate previously conducted MD simulations with BPTF-PHD-BRD. These findings support the idea that, in solution, recognition of multiply modified H3 peptides in solution is mediated by shifting the conformational equilibrium of the reader domains towards the more globular, compact structure that is different from the major ensembles observed in crystallography, although we cannot rule out higher order complex formation under conditions that more closely mirror crystallography conditions. However, additional experimental support for formation of a monomeric peptide-protein complex mediated by induced conformational change in solution was further verified through Diffusion Ordered NMR Spectroscopy (DOSY; **Supplementary Figure S9**), oligomeric chemical crosslinking (**Supplementary Figure S4**), and native polyacrylamide gel electrophoresis (**Supplementary Figure S5**). Furthermore, detailed analysis of deposited crystal structures revealed that conformational heterogeneity exists between different models, complementing the general conclusion from our solution-based studies that conformational changes are possible and, perhaps, necessary to accommodate ligand binding.

### Small Molecule-Peptide Conjugates Engage Both Domains of BPTF

Elucidating the selective, combinatorial readout of K4me_3_ and K14ac PTMs within the H3 peptide sequence provides a scaffold for the design of bivalent inhibitors which could engage both domains of the BPTF-PHD-BRD. Although existing crystal structures estimated a distance of 70 Å between the BPTF BRD and PHD binding sites,^15^ measurement of the amino side chains of H3K4 and H3K14 from an NMR solution structure of the H3(1-15) peptide (PDB ID 2FUI)^41^ demonstrated that these moieties occupy distances ranging from 13 to 33 Å, with a median distance of approximately 26 Å (**Supplementary Figure S8**). Based on these insights, we sought to discover the first peptide-small molecule conjugate by modifying the H3 peptide structure to include a BRD inhibitor. BZ1-THQ, reported previously by Zahid et al., demonstrates sub-micromolar affinity for the BPTF BRD and contains a solvent-exposed secondary amine for attachment of additional components.^28,69^ Additionally, shorter H3K4me_3_ peptides are known to exhibit comparable PHD binding affinities relative to their longer counterparts.^70^ Since PEG5 linkers have been shown to span distances of ∼26 Å,^71^ we fused a truncated H3(1-5)K4me_3_ peptide to BZ1-THQ via a PEG5 linker resulting in the dual conjugate **MJ-1**(**Figure 8A**) that should be compatible with the compact form observed in solution measurements. DSF of BPTF-PHD-BRD with **MJ-1** showed a ΔT_m_ of +9.28 °C (**Figure 8B and 8E**), similar to the maximum stabilization observed previously with H3 peptides. Next, individual binding site contributions of **MJ-1** were investigated using AlphaScreen^TM^. We observed potent BRD and PHD interactions (PHD IC_50_ = 70 nM, BRD IC_50_ = 10 nM; **Figure 8C-E**) with **MJ-1**, representing roughly 50 and 25-fold enhancements over the IC_50_ values observed in this assay for H3(1-20)K4me3 (**5**) and BZ1-THQ (250 nM),^28^ respectively, and were suggestive of a cooperative binding mode as observed with dually modified H3 peptides. ITC analysis afforded a K_d_ of 1.95 μM that was accompanied by an N value close to 1 and substantial entropic penalty (**Figure 8F**). Thus, we conclude that **MJ-1** likely participates in a cooperative and simultaneous interaction with the BPTF-PHD-BRD that is permitted by domain rearrangement, as observed with multiply modified H3 peptides. Importantly, despite using a higher-affinity BRD ligand, the entropy costs associated both from **MJ-1** binding and protein conformational changes do not improve binding potencies over H3 peptide ligands in ITC. DSF and AlphaScreen^TM^ was performed with mutant BRD and PHD proteins to verify the binding location of **MJ-1** and glean insight on domain dependence of bivalent inhibitor binding. Inactivation of PHD binding resulted in ∼3 °C loss of thermal stabilization and a 70-fold reduction in potency towards the BRD (**Figure 8B-E**), supporting the importance of maintaining PHD functionality also for the binding of **MJ-1**. Additionally, inactivation of the BRD resulted in a near complete loss of thermal stabilization and a 17-fold reduction in inhibition against the PHD (**Figures 8B-E**).

**Figure 8.**
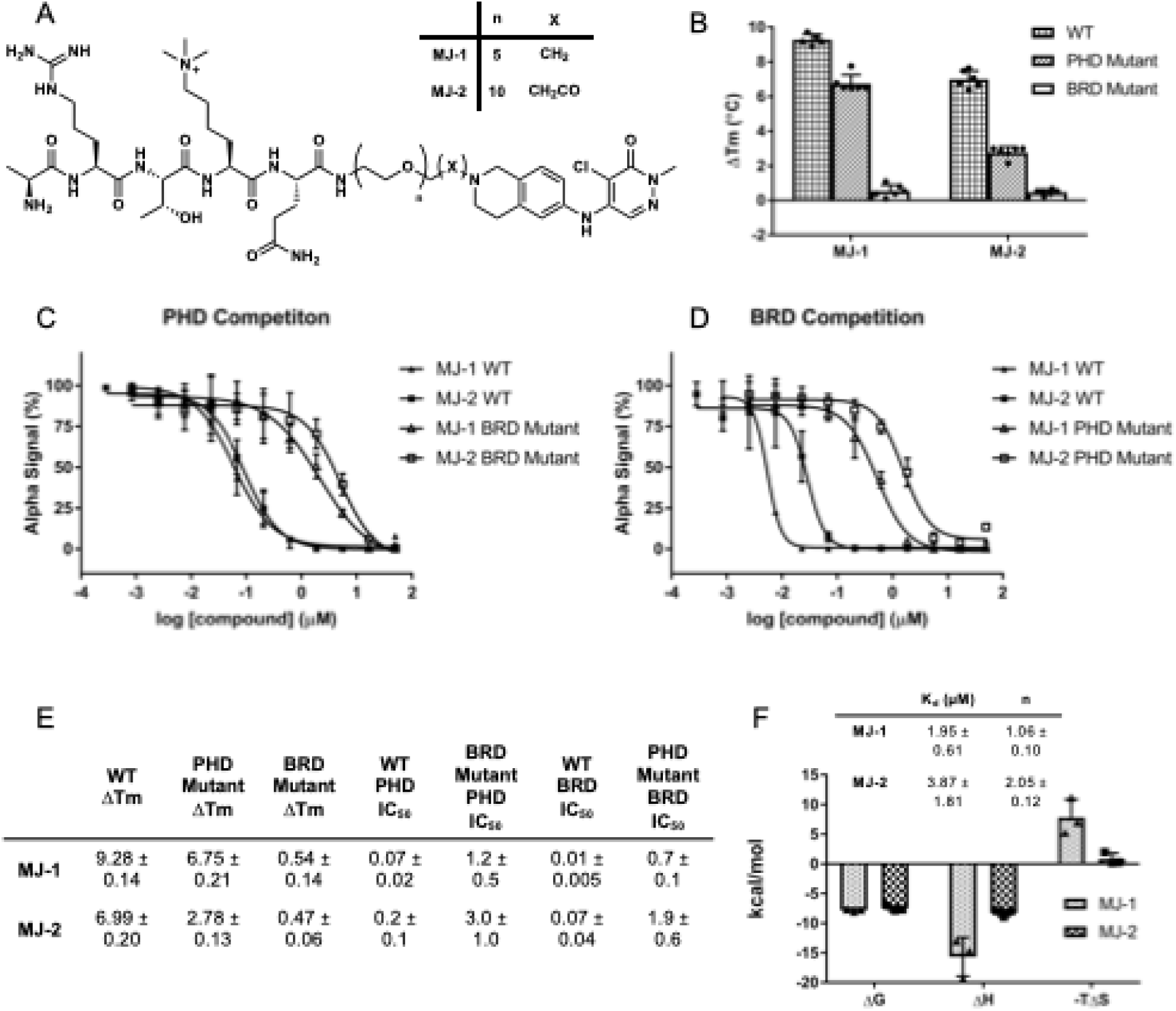
**A.** Structures of peptide-small molecule conjugates. B.) DSF and C,D.) Domain specific AlphaScreen^TM^ competition experiments with WT BPTF and mutants. E.) Summary table of DSF ΔT_m_ and AlphaScreen^TM^ IC_50_ values. F.) ITC thermodynamics, stoichiometry, and binding affinity data for bivalent inhibitors.

Next, the PEG10 derivative **MJ-2** was synthesized (**Figure 8A**). We envisioned this elongated molecule would place the Kme_3_ and BZ1-THQ pharmacophores in a suboptimal orientation that is incompatible with simultaneous engagement of both domains (as seen with **MJ-1**), resulting in general losses in binding affinity, reduced entropic penalties of binding, and inflated n-values. Thermal stabilization events and domain-specific competitive inhibition values were less potent than **MJ-1**, most exemplified by a 7-fold reduction in WT BRD IC_50_ (**Figure 8D and 8E**). A K_d_ of ∼4 µM was observed for **MJ-2**, coupled with a reduced enthalpic contribution, minimal entropic penalty, and a higher N-value (∼2) relative to **MJ-1** when a 1:1 fitting model was used (**Figure 8F**). We conclude that the inflated N-value and reduced entropic penalty suggest that **MJ-2** is incapable of inducing the conformational change associated with simultaneous PHD-BRD engagement due to its longer length, whereas the shorter PEG5 linker of **MJ-1** effectively mimics a multiply modified, cis-engaging H3K4me_3_K14ac peptide and permits cooperative, bivalent inhibition. We note that the loss of potency observed for **MJ-2** relative to **MJ-1** may reflect binding costs associated with unique motifs including a PEG amide linkage to BZ1-THQ, a longer PEG linker length, or a combination of both structural alterations.

### Conclusions

In this work, we expand upon existing studies that posit BPTF-PHD-BRD can engage H3 peptides containing multiple PTMs in *cis*. The acetyllysine binding preferences and structural mechanisms permitting tandem readability of H3K4me_3_ and H3Kac PTMs by BPTF were largely underexplored. We identified H3K14ac as the primary PTM responsible for BRD binding using multiple biophysical assays with higher affinity than the canonical H4K16ac modification, and show that this PTM exhibits cooperativity with H3K4me_3_. We demonstrate evidence for two plausible structural mechanisms driving H3K4me_3_K14ac molecular recognition at the peptide level based on crystallography and solution measurements: dimerization and conformational change. Our work suggests that the BPTF-BRD-PHD “structure” is best described by an interconverting ensemble of states. In solution, the conformational equilibrium is shifted towards a more compact state upon peptide binding that brings together both domains in *cis*, while crystallography shows that dimerization with the peptide binding across monomers in *trans* may also play a role in H3 ligand recognition.

Notably, these results expand the epigenetic histone code for BPTF at the peptide level to include H3K14ac. H3K14ac has been identified as a potential prognostic marker in oral squamous cell carcinoma and triple-negative breast cancer.^72,73^ Although H3K14ac is traditionally classified as a transcriptional activation mark,^74^ it is interpreted by many BRD-containing proteins and is associated with context-dependent biological processes. Indeed, the Brahma-associated protein (BAP) BRD engages this PTM in *Drosophila* and is associated with transcriptional activation.^75^ However, the Zinc Finger MYND-Type Containing 8 (ZMYND8) PHD-BRD cassette recognizes dual H3K4Me_1_K14ac modifications and functions as a repressor of metastasis-related genes in prostate cancer cells.^76^ Similarly, the bromodomain adjacent to zinc finger domain protein 2A (BAZ2A) BRD is a reader of H3K14ac PTMs at inactive enhancers and mediates transcriptional silencing of differentiation-related genes.^77^ Given the seemingly multifaceted role of H3K14ac in transcriptional regulation, future work investigating the biological processes governed by BPTF-mediated recognition of this PTM in cells is necessary.

Using the Histone H3 sequence as a template, we designed **MJ-1:** the first example of a small-molecule-peptide conjugate targeting BPTF. **MJ-1** exhibits single-digit micromolar affinity by ITC and nM IC_50_ values for both chromatin binding domains in AlphaScreen™, thereby overcoming druggability challenges associated with PHD fingers.^31^ Due to the discrepancy in ITC and AlphaScreen^TM^ potencies, we note the AlphaScreen^TM^ format may be less sensitive to entropic penalties given the multivalent display of ligands and protein. This may be due to avidity effects and reduced entropic contributions that arise from immobilization of binding partners, as observed in assays like surface plasmon resonance.^53,78^ Peptide-drug conjugates are an emerging class of therapeutics demonstrating potential as anticancer, antiviral, and antibiotic agents.^79,80^ Recent efforts have identified potent peptide-drug conjugates for the TRIM24 PHD-BRD cassette, further proving the tractability of developing bifunctional molecules derived from histone peptide ligands that target PHDs.^81^ To date, only 11 of the 291 PHD fingers found in humans^82^ have reported chemical matter (**Supplementary Table 4**). Since existing evidence suggests that PHD finger inhibition has therapeutic potential in multiple disease states,^32,83,84^ future endeavors will center on the development of cell-active peptide conjugates targeting BPTF. Collectively, our findings merge histone code expansion studies and rational drug design to uncover structural models and therapeutic starting points toward an understudied epigenetic reader. Such information is necessary for better elucidating the biological function of BPTF in normal and diseased states, as well as inspiring the design of next-generation chemical probes that target BPTF and other PHD finger containing proteins.

## Associated Content

The Supporting Information is available free of charge at (…).

Materials and Methods, MALDI-TOF spectra, HPLC purity analyses, synthetic schemes, ^1^H & ^13^C NMR, DSF melt curves, AlphaScreen™ inhibition curves, ITC binding isotherms, crystallography data, SAXS data, LiP-MS structure mapping, DOSY NMR, Native and SDS PAGE.

## Acknowledgements

This work was in part supported by the NIH R35GM140837 (W.CK.P.) and NIH NCI R01CA290805 (M.F.) and American Lebanese Syrian Associated Charities, ALSAC (M.F.). J.E.M. was supported by the NIH NCATS 1T32TR004385 through the University of Minnesota Clinical and Translational Science Institute. M.S. was additionally supported by a National Institutes of Health Chemical Biology Interface Training Grant T32GM132029. The content of this paper is solely the responsibility of the authors and does not necessarily represent the official views of the NIH. ITC experimentation was completed using instrumentation at the University of Minnesota Institute for Therapeutics Discovery and Development. SAXS data was collected at the Advanced Light Source (ALS) at the SIBYLS beamline, a national user facility operated by Lawrence Berkeley National Laboratory on behalf of the Department of Energy, Office of Basic Energy Sciences, through the Integrated Diffraction Analysis Technologies (IDAT) program, supported by DOE Office of Biological and Environmental Research. Additional support comes from the National Institute of Health project ALS-ENABLE (P30GM124169). We thank Dr. Gregory Hura, Dr. Michal Hammel, and other scientists and staff members at the ALS facility for assistance with SAXS experimentation. We thank Ketevan Chomakhidze for assistance with MALDI-TOF analyses. We thank Elliott Peterson for assistance with mutant BPTF plasmid design. We also thank Prof. Tim Lodge and Prof. Kanishk Jain for helpful discussions. Figures were created with Biorender. We thank X-ray center at St Jude Children’s Research Hospital for the crystallization facility. Also, we thank NSLS-II 17-ID beamline facility for X-ray data collection. This research used resources (17ID beamline) of the National Synchrotron Light Source II, a U.S. Department of Energy (DOE) Office of Science User Facility operated for the DOE Office of Science by Brookhaven National Laboratory under Contract No. DE-SC0012704. The Center for BioMolecular Structure (CBMS) is primarily supported by the National Institutes of Health, National Institute of General Medical Sciences (NIGMS) through a Center Core P30 Grant (P30GM133893), and by the DOE Office of Biological and Environmental Research (KP1605010). The 500 MHz NMR spectrometer used for this work was funded, in part, by NIH Award Number S10OD011952

## Author Information

### Corresponding Author

**William C.K. Pomerantz** – Department of Medicinal Chemistry, University of Minnesota, Minneapolis, Minnesota 55455, United States; Department of Chemistry, University of Minnesota, Minneapolis, Minnesota 55455, United States; orcid.org/0000-0002-0163-4078;

### Authors

**Joseph E. McPherson** – Department of Medicinal Chemistry, University of Minnesota, Minneapolis, MN 55455 United States; orcid.org/0009-0008-7072-9645

**Molly S. Sneddon** – Department of Chemistry, University of Minnesota, Minneapolis, Minnesota 55455, United States; orcid.org/0000-0002-7249-3401

**Richard K Begyinah** – Department of Chemical Biology and Therapeutics, St. Jude Children’s Research Hospital, Memphis, Tennessee, 38105, United States; orcid.org/0009-0005-8134-6867 **Kesavan Babu** – Department of Chemical Biology and Therapeutics, St. Jude Children’s Research Hospital, Memphis, Tennessee, 38105, United States; orcid.org/0000-0002-7895-7246

**Kerstin E. Peterson** – Department of Chemistry, University of Minnesota, Minneapolis, Minnesota 55455, United States; orcid.org/0000-0002-0291-4039

**Ryan F. Friess** – Department of Medicinal Chemistry, University of Minnesota, Minneapolis, MN 55455 United States; orcid.org/0009-0006-8218-6583

**Thomas G. Smith** – Department of Chemistry, University of Minnesota, Minneapolis, Minnesota 55455, United States; orcid.org/0000-0003-1366-8053

**Natalia Y. Tretyakova** – Department of Medicinal Chemistry, University of Minnesota, Minneapolis, Minnesota 55455, United States; Department of Chemistry, University of Minnesota, Minneapolis, Minnesota 55455, United States; orcid.org/0000-0002-0621-6860

**Marcus Fischer** – Department of Chemical Biology and Therapeutics, St. Jude Children’s Research Hospital, Memphis, Tennessee, 38105, United States; orcid.org/0000-0002-7179-2581

## Notes

The authors declare no competing financial interest.

## Notes

### Competing Interest Statement

The authors have declared no competing interest.

