## Supplementary Information for "Bivalent Inhibitor Development for a Dynamic Epigenetic Tandem Reader Protein Using a Combinatorial Histone Code"

#### Experimental Section

*Synthesis and purification of histone H3 peptides **1-12**:*
Histone H3 peptides were synthesized similar to previously described methods[1] on a Liberty Blue Automated Microwave Peptide Synthesizer (CEM) using Fmoc protected amino acids and NovaSyn TGR resin (Novabiochem). Reactions were conducted on a 0.025 mmol scale. Oxyma (0.250 M), DIC (0.125 M), and Fmoc-canonical amino acids (0.2 M) were prepared in DMF. Fmoc-Lys-Kac (0.2 M; Chem-Impex), Fmoc-Lys-$Me_3$ (0.1 M; Ambeed) and Fmoc-Ahx (0.1; Ambeed) were prepared in DMF. All coupling steps proceeded via conventional coupling methods, except Arg (Double Arg coupling at 75 °C) and Fmoc-Lys-$Me_3$ and Fmoc-Ahx (Double conventional coupling). Following synthesis, peptides were cleaved from resin with solution containing 95% TFA, 2.5% TIPS, and 2.5% $H_2O$ for 3 hours at room temperature. TFA was removed under nitrogen and peptides were precipitated with cold ether. Peptides were pelleted via centrifugation and ether was decanted. The crude pellet was dried under nitrogen and the resulting solid was dissolved in $H_2O$ + 0.1% TFA and purified using semi-preparative HPLC on a C18 column (25005-159070; Thermo Fisher Scientific) via a 0-20% ACN in $H_2O$ + 0.1% TFA gradient spanning 20 minutes. Product masses were verified by MALDI-TOF. Percent purity analyses were conducted using an analytical C18 column (201HS54; Avantor/Vydac) via a 0-30% ACN in $H_2O$ + 0.1% TFA gradient spanning 50 minutes following a 10-minute isocratic hold at 0% ACN at the start of the run. All peptides were at least 95% pure by peak area analysis spanning 5-50 minutes, with percentages contributed by the blank omitted.

*Synthesis of Bivalent Inhibitors*
*5-((2-(17-amino-3,6,9,12,15-pentaoxaheptadecyl)-1,2,3,4-tetrahydroisoquinolin-6-yl)amino)-4-chloro-2-methylpyridazin-3(2H)-one (**14**):*
BZ1-THQ was synthesized according to previous literature.[2] Tosylate **13** was prepared as previously described[3] from tert-butyl (17-hydroxy-3,6,9,12,15-pentaoxaheptadecyl)carbamate and the crude product was used in subsequent steps. BZ1-THQ (0.200 g, 1.0 eq, 0.17 mmol) was dissolved in *N,N*-diisopropylethylamine (0.30 mL, 10 eq, 1.7 mmol) and 1,4-dioxane (1 mL, 163 mM) and **13** was added. The mixture was heated to 120 °C in a sealed tube and stirred for 18 hours. Upon reaction completion, the 1,4-dioxane was removed by rotary evaporation. The crude mixture was extracted into dichloromethane (3 x 20 mL), then washed with saturated sodium bicarbonate (3 x 20 mL) and brine (1 x 20 mL). The organic layer was dried with magnesium sulfate, filtered, and concentrated by rotary evaporation. Finally, purification was performed using flash chromatography (Combiflash Rf system: 4g of silica, DCM/methanol, 0-50% methanol, 15 min) to yield **14** as a brown solid (37% yield). $^1$H NMR (500 MHz, chloroform-d): δ 7.62 (s, 1H), 7.10 (dd, J = 8 Hz, 1H), 6.99 (d, J = 13.6, 2H), 6.63 (br. s, 1H), 4.00 (br. s, 2H), 3.86 (br. s, 2H), 3.73 (s, 3H), 3.67-3.58 (m, 22H), 3.50 (t, 5.4 Hz, 2H), 3.43 (br. s, 1H), 3.27 (br. s, 2H), 3.10-3.05 (m, 10H), 1.41 (s, 9H). DCM impurity at 5.3 ppm, DIEA impurity at 1.5 ppm. $^{13}$C NMR (126 MHz, chloroform-d) δ 157.84, 142.07, 128.66, 128.19, 126.86, 125.88, 123.87, 122.03, 70.56, 70.53, 70.48, 70.43, 70.40, 70.19, 67.06, 56.43, 53.72, 50.36, 47.11, 42.02, 40.33, 40.25, 28.42, 21.30, 18.91, 18.60, 17.36, 12.05; HRMS (ESI-TOF) calculated for $C31H49ClN5O8^+$ $[M+H]^+$ = 654.3264, observed 654.3254.

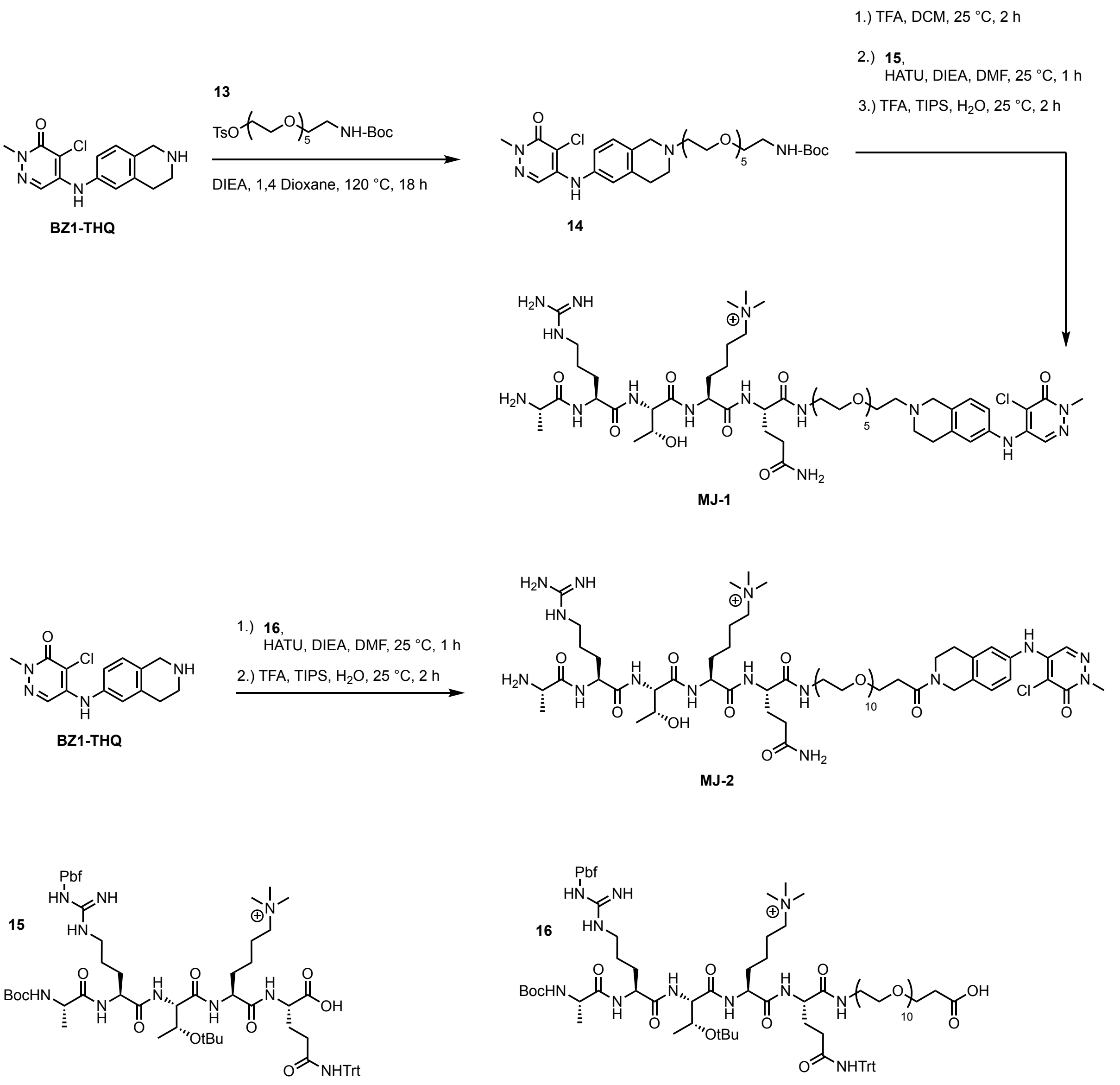

Scheme 1. Synthesis of peptide-small molecule conjugates **MJ-1** and **MJ-2**.

*Synthesis of H3(1-5) Peptide Intermediates* ***15 and 16****:*

The histone H3 peptides were synthesized using 2-chlorotrityl resin (Chem-Impex) (0.1 mmol scale). Oxyma (1 M), DIC (1 M), and Fmoc amino acids (0.2 M) were prepared in DMF. 2-chlorotrityl resin (0.100 g, 1.0 eq, 0.32 mmol) was preloaded with either Fmoc-Gln(Trt)-OH (1.95 g,10 eq, CAS 132327-80-1, Chem-Impex) or Fmoc-NH-PEG10-CH2CH2COOH (0.800 g, 10 eq, 3.2 mmol, CAS 2101563-45-3, PurePEG) by stirring at room temperature for 1 hour in anhydrous dichloromethane (120 mM) with *N,N*-diisopropylethylamine (0.557 mL, 10 eq, 3.2 mmol). After preloading, the resin was washed in vacuo (5 x 10 mL MeOH) and the subsequent amino acids were coupled using a Liberty Blue Automated Microwave Peptide Synthesizer. Boc-Ala-OH (Sigma Aldrich) was used in place of Fmoc-Ala-OH at the N-terminus. The resulting peptide was washed in vacuo (3 x 10 mL DMF and DCM). Resin cleavage was then performed by stirring the peptide in 1% TFA and 2% TIPS in DCM for 2 hours. The crude peptide was precipitated with cold diethyl ether and centrifuged to form a pellet from which the ether was decanted. Residual diethyl ether was evaporated over an N2 stream to afford white solids. The product masses were confirmed by MALDI-TOF (**15** Exp. = 1295.711, Obs. = 1295.699; **16** Exp. = 1807.010, Obs. = 1806.947), and the crude peptides were then used in subsequent syntheses.

*(20S,23S)-20-(3-amino-3-oxopropyl)-23-((2S,3R)-2-((S)-2-((S)-2-aminopropanamido)-5-guanidinopentanamido)-3-hydroxybutanamido)-1-(6-((5-chloro-1-methyl-6-oxo-1,6-dihydropyridazin-4-yl)amino)-3,4-dihydroisoquinolin-2(1H)-yl)-N,N,N-trimethyl-19,22-dioxo-3,6,9,12,15-pentaoxa-18,21-diazaheptacosan-27-aminium (**MJ-1**):*
The Boc protecting group was removed from intermediate **14** using 5 eq of TFA in dichloromethane at room temperature for two hours. Solvents were removed over an N2 stream to afford crude product that was used in subsequent steps. The H3(1-5) peptide **15** (0.010 g, 5 eq, 0.011 mmol) was dissolved in dry DMF (100 mM) and preincubated with HATU (0.005 g, 5.2 eq, 0.012 mmol) and *N,N*-diisopropylethylamine (0.008 mL, 20 eq, 0.046 mmol) by stirring for 10 minutes at room temperature. Then, compound **14** (0.006 g, 1.0 eq, 0.002 mmol,), dissolved in DMF (100 mM) and *N,N*-diisopropylethylamine (0.08 mL, 20 eq, 0.046 mmol), was added, and the mixture was stirred at room temperature for 1 hour. The crude product underwent global deprotection with 95% TFA, 2.5% TIPS, and 2.5% $H_2O$ for 2 hours at room temperature. Solvents were removed over an N2 stream, and the products were precipitated with cold diethyl ether and centrifuged to form a pellet from which the ether was decanted. Residual diethyl ether was evaporated by an N2 stream. The resulting solid was dissolved in $H_2O$ + 0.1% TFA and purified using semi-preparative HPLC on a C18 column (25005-159070; Thermo Fisher Scientific) via a 0-20% ACN in $H_2O$ + 0.1% TFA gradient spanning 20 minutes. Product masses were verified by MALDI-TOF. Percent purity analyses were conducted using an analytical C18 column (201HS54; Avantor/Vydac) via a 0-50% ACN in $H_2O$ + 0.1% TFA gradient spanning 50 minutes, calculated by integration of peak are spanning 5-50 minutes.

*(36S,39S)-36-(3-amino-3-oxopropyl)-39-((2S,3R)-2-((S)-2-((S)-2-aminopropanamido)-5-guanidinopentanamido)-3-hydroxybutanamido)-1-(6-((5-chloro-1-methyl-6-oxo-1,6-dihydropyridazin-4-yl)amino)-3,4-dihydroisoquinolin-2(1H)-yl)-N,N,N-trimethyl-1,35,38-trioxo-4,7,10,13,16,19,22,25,28,31-decaoxa-34,37-diazatritetracontan-43-aminium (**MJ-2**):*
The H3(1-5)PEG10-CH2CH2-COOH peptide **16** (0.015 g, 5 eq, 0.008 mmol) was dissolved in dry DMF (100 mM) and preincubated with HATU (0.003 g, 5.2 eq, 0.002 mmol) and *N,N*-diisopropylethylamine (0.005 mL, 15 eq, 0.025 mmol) by stirring for 10 minutes at room temperature. Then, BZ1-THQ was dissolved in DMF (0.002 g, 1 eq, 0.002 mmol) and *N,N*-diisopropylethylamine (0.005 mL, 15 eq, 0.025 mmol), was added, and the mixture was stirred at room temperature for 1 hour. The crude product underwent global deprotection with 95% TFA, 2.5% TIPS, and 2.5% $H_2O$ for 2 hours at room temperature. Solvents were evaporated over an N2 stream, and peptides were precipitated with cold diethyl ether and centrifuged to form a pellet from which the ether was decanted. Residual diethyl ether was evaporated over an N2 stream. The resulting solid was dissolved in $H_2O$ + 0.1% TFA and purified using semi-preparative HPLC on a C18 column (25005-159070; Thermo Fisher Scientific) via a 0-20% ACN in $H_2O$ + 0.1% TFA gradient spanning 20 minutes. Product masses were verified by MALDI-TOF. Percent purity analyses were conducted using an analytical C18 column (201HS54; Avantor/Vydac) via a

0-50% ACN in $H_2O$ + 0.1% TFA gradient spanning 50 minutes, calculated by integration of peak area spanning 5-50 minutes.

*Expression and purification of BPTF-PHD-BRD constructs*
A gene sequences encoding residues 2865 through 3033 of the c-terminal PHD-BRD region of full length BPTF (UniProt ID Q12830) with an n-terminal His-9 tag and TEV cleavage site (MGSSHHHHHHHHHSSGENLYFQGHMDTKLYCICKTPYDESKFYIGCDRCQNWYHGRC VGILQSEAELIDEYVCPQCQSTEDAMTVLTPLTEKDYEGLKRVLRSLQAHKMAWPFLE PVDPNDAPDYYGVIKEPMDLATMEERVQRRYYEKLTEFVADMTKIFDNCRYYNPSDSPF YQCAEVLESFFVQKLKGFKA) was inserted into a pET-28a(+) vector (Genscript) harboring kanamycin resistance. The resulting plasmid DNA (1 ng) was transformed into BL21 Star (DE3) competent cells (Thermo Fisher Scientific) via heat shock for 45 seconds at 42 °C followed by recovery in SOC outgrowth medium (New England Biolabs) for 60 minutes at 37 °C with shaking at 250 rpm. Cells were plated on LB agar plates containing kanamycin (50 µg/ml) and incubated at 37 °C overnight. Successful transformants were picked for glycerol stock preparation and 5 mL primary culture inoculation of LB containing kanamycin (50 µg/mL). Primary cultures were grown overnight at 37 °C with shaking and secondary 1 L cultures were grown under the same conditions from these primaries to $OD_{600}$ 0.6 to 0.8. 1 L cultures were cooled to 18 °C and 100 µM $ZnCl_2$ was supplemented to the cultures. 400 µM IPTG was added to induce protein expression overnight. The following day, cells were pelleted via centrifugation (10 minutes at 4000 xg) and stored at -80 °C. All steps regarding protein purification were performed on ice or in a cold room. Frozen cell pellets were resuspended in lysis buffer (50 mM $Na_2HPO_4$, 500 mM NaCl, 5 µM $ZnCl_2$, pH 7.4) and lysed via sonication (30 seconds on, 60 seconds off, 40% amplitude, 12 minutes total sonication). Lysates were cleared via centrifugation (45 minutes at 12,000 xg) and the cleared lysate was applied to a 5 mL HisTrap FF crude column (Cytiva) equilibrated with wash buffer (50 mM $Na_2HPO_4$, 500 mM NaCl, 5 µM $ZnCl_2$, 20 mM imidazole, pH 7.4) and the column was washed with >20 column volumes until all unbound proteins were depleted. Remaining His-tagged BPTF-PHD-BRD was eluted (50 mM $Na_2HPO_4$, 500 mM NaCl, 5 µM $ZnCl_2$, 400 mM imidazole, pH 7.4) and exchanged into the appropriate storage buffer using a 50 mL HiPrep desalting column (Cytiva). Buffer composition was dependent on the subsequent assay being used and is outlined below for each assay method.

*Site-directed mutagenesis*
Site-directed mutagenesis was conducted with the QuikChange II Mutagenesis Kit (Agilent) using the manufacturer's protocols. One reaction was prepared for each mutant. 50 µL reactions were prepared containing 125 ng of each primer (4 µL total), 50 ng wild type BPTF-PHD-BRD plasmid (1µl), dNTP mix (1µl), 10x reaction buffer (5 µl), and $H_2O$ (39 µl). 1 µl of *PfuUltra* HF DNA polymerase was added and cycling was performed using an initial, 95 °C 30 second denaturation step followed by 16 subsequent cycling steps containing the following: 30 seconds at 95 °C, 1 minute at 55 °C, 6 minutes at 68 °C. After thermal cycling, reactions were cooled on ice and 1 µl of *Dpn I* restriction enzyme was added and incubated for 60 minutes at 37 °C. The resulting plasmid was transformed into XL1-Blue supercompetent cells (Agilent) using the same protocol as described for transformation of wild type BPTF-PHD-BRD. Successful incorporation of the mutant amino acids was confirmed via DNA sequencing. The mutant BPTF-PHD-BRD

plasmids were transformed into Rosetta(DE3) competent cells (Novagen) and protein expression and purification was conducted as described for wild type BPTF-PHD-BRD.

*Differential scanning fluorimetry*

Melting temperature experiments were performed on a QuantStudio™ 5 PCR system (Thermo Fisher Scientific) using MicroAmp 384-well PCR reaction plates (Thermo Fisher Scientific) and MicroAmp plate seals. BPTF-PHD-BRD was exchanged into DSF buffer (50 mM HEPES, 100 mM NaCl, pH 7.4). 20 µL reactions containing DSF buffer, BPTF-PHD-BRD (20 µM), ligand (100 µM), and 1x SYPRO Orange dye (diluted from 1000x stock; Thermo Fisher Scientific) were plated and sealed. Fluorescence was monitored as a function of increasing temperature gradient from 25 to 95 °C at a ramp rate of 0.5 °C per minute. Experiments with inhibitors dissolved in DMSO were matched to 2% DMSO (v/v). Ligands were monitored for autofluorescence and dye interactions using ligand only and no protein controls. Protein melt curves were analyzed in the Design and Analysis 2 Software (Thermo Fisher Scientific) and $T_m$ values were calculated using the derivative function. Reported $\Delta T_m$ values for ligands represent the change in average $T_m$ between BPTF-PHD-BRD unliganded and BPTF-PHD-BRD with ligands. For visualization, melt curves were plotted in GraphPad Prism 7 using the average fluorescence reading as a function of temperature (°C) and the resulting $\Delta T_m$ values were annotated directly.

*Alphascreen™ competition assays*

Competition experiments were performed using the Alphascreen™ Nickel Chelate Detection Kit (Revvity) based on existing protocols for BRD[2] and PHD[4] competitive inhibition assays. BPTF-PHD-BRD was exchanged into the same buffer used for DSF. All reactions and reagents were prepared in assay buffer (50 mM HEPES, 100 mM NaCl, 0.05% CHAPS, 0.1% BSA, pH 7.4) and DMSO concentration was matched to 1% for experiments with inhibitors. 20 µL reactions contained 125 nM His-BPTF-PHD-BRD, 125 nM acetylated histone H4 (1-21) peptide probe for BRD competition experiments (SGRGKacGGKacGLGKacGGAKacRHRKVGGK-biotin, Anaspec), 20 µg/ml streptavidin donor beads, and 20 µg/ml nickel chelate acceptor beads. These conditions were matched for PHD competition experiments, except 8 nM trimethylated histone H3 (1-21) peptide probe (ARTKme$_3$QTARKSTGGKAPRKQLAGGYK-biotin, Active Motif) was used instead. 5 µl of serially diluted competing ligand with His-BPTF-PHD-BRD was added to a 384-well ProxiPlate (Revvity), followed by 5 µl of biotinylated peptide probe, then 10 µl of combined bead solution. Plates were sealed, incubated for 45 minutes at room temperature, and read using Alphascreen™ luminescence filter settings (520 nm – 620 nm) on a Tecan Spark plate reader. All steps involving Alphascreen™ bead preparation, dispensing, and incubation were completed under subdued lighting. Percent alpha signal values were determined by normalization to uninhibited, no ligand controls (100%) and the background signal generated at full inhibition (0%). $IC_{50}$ values were calculated in GraphPad Prism 7 using the sigmoidal four-parameter logistic curve fit.

*Isothermal titration calorimetry*
ITC experiments were conducted similar to established conditions.[5] BPTF-PHD-BRD was exchanged into buffer containing 50 mM $Na_2HPO_4$, 100 mM NaCl, 2 mM TCEP, pH 7.4. Protein and ligand concentrations were verified using A280 and the extinction coefficient of BPTF-PHD-BRD (32,360 $M^{-1}cm^{-1}$ in aqueous solution), tyrosine (1,490 $M^{-1}cm^{-1}$ in aqueous solution; peptides) or BZ1-THQ (2,333 $M^{-1}cm^{-1}$ in DMSO; inhibitors). Experiments were performed on a MicroCal PEAQ-ITC instrument (Malvern Panalytical). Ligands were injected into solutions of BPTF-PHD-BRD at approximately 10-fold excess of ligand. Each run consisted of 18 injections (volume = 2 µL, duration = 4 s, spacing = 150 s) following an initial 0.4 µL equilibration injection that was excluded from data analysis. Runs were conducted using the standard instrument parameters provided by Malvern (temperature = 25 °C, reference power = 10 µcal/s, feedback mode = high, stir speed = 750 rpm). Data was analyzed in the MicroCal PEAQ-ITC Analysis Software using the one set of sites fitting model. Peaks were integrated and the baseline was subtracted to give binding isotherms with associated affinities, free energy changes, enthalpy changes, entropy changes, and stoichiometries. Reported values for each ligand are the average and standard error of experimental triplicates.

*DOSY NMR*
$^1$H DOSY NMR spectra were acquired at 25 °C on a Bruker Avance III HD 500 MHz spectrometer equipped with a 5 mm TCI Prodigy cryoprobe using a stimulated echo sequence with bipolar gradient pulses and excitation sculpting for water suppression (stebpesgp1s). Samples were prepared in 50 mM phosphate, 100 mM NaCl, pH 7.4 with 10% D2O and analyzed in a D2O-matched Shigemi tube. Data were collected over 16 linear gradient steps ranging from 2% to 98% maximum gradient strength (NS = 256 per step). Experimental parameters included a diffusion time (big delta, d20) of 200 ms and a total diffusion gradient pulse length (little delta, 2 x p30) of 4.62 ms (p30 = 2310 us). Samples were individually tested as 1) peptide only at either 35 or 50 µM, protein only at either 35 or 50 µM, and 3) peptide plus protein at either 35 or 50 µM. Diffusion coefficients were calculated as described.[6]

*Crystallography*
The Human BPTF PHD-BRD construct fused with GST tag at the N-termini in pGEX-6P-1 vector obtained from GenScript. This protein was recombinantly produced in *E. coli* BL21 (DE3) cells with 2XYT medium and 100 uM $ZnCl_2$ is added to the cells. The cells were grown until they reached $OD_{600}$ of 0.6 -0.8. 0.8 mM IPTG was added and incubated for 16 hours at 18°C. The purification protocol was adopted from the previous report. The collected cell pellets were mixed with lysis buffer, 50 mM HEPES pH 7.5, 500 mM NaCl, 5 mM DTT containing SIGMAFAST inhibitor cocktail (Sigma Aldrich, #S8830). Cells were subjected to sonication and soluble protein collected by centrifugation at 35,000 rpm for 1 hr. The GST-tagged BPTF PHD-BRD protein was purified by GSTrap column and 20 mM glutathione was used to elute the protein. Protein fractions were pooled and dialyzed with PreScission protease overnight and re-applied to the GSTrap HP column to purify cleaved PHD-BRD protein. Final purification was performed by Superdex-75 gel filtration column in 20 mM HEPES pH 7.5, 100 mM KCl, and 5 mM DTT. The protein fractionated as homogenous population and the protein was concentrated to 10.40 mg/ml. The frozen protein was stored at -80°C freezer.

The concentrated BPTF PHD-BRD was incubated with final concentration of 670 μM of peptide 3 at room temperature for 2 hrs. In same way, the binding reaction of a short H3(1-7)K4me$_2$ peptide (520 μM) with the BPTF PHD-BRD is initiated. Using the hanging drop method in a 96 well MRC plate with well solutions of 100 mM HEPES pH 7.5, 10% 2-propanol, and 14- 16% PEG 4000 at 4°C, and the crystals appeared in 3 to 5 days. Crystals were cryoprotected by 20% glycerol, and the liquid nitrogen-frozen crystals were shipped to synchrotron. X-ray data was collected at NSLS-II-17ID-1. Initial structure was obtained using molecular replacement with 2F6N[7] as a PDB template. The structure was further refined and built using Phenix[8] and Coot[9] respectively. The structure figures were generated using Pymol (Schrödinger, New York).

*Dihydrazide sulfoxide mediated crosslinking and SDS PAGE*
BPTF-PHD-BRD crosslinking experiments were performed using previously established conditions.[10] Series of reactions (30 μL final volume) in phosphate buffer (50 mM $Na_2HPO_4$, 100 mM NaCl, pH 7.4) were prepared containing 20 μM, 80 μM, and 200 μM of His-cleaved BPTF-PHD-BRD with and without 3-fold molar excess of peptide 3 and incubated for 30 minutes at room temperature. To all reactions DHSO (CAS 1971857-82-5) was added to a final concentration of 400 μM with DMTMM (CAS 3945-69-5) to a final concentration of 800 μM, and reactions were incubated for 120 minutes at room temperature. 5.5 μg of resulting BPTF-PHD-BRD was removed from each reaction and boiled at 95 °C for 10 minutes in Nu-PAGE SDS sample buffer (Novex). Denatured samples were analyzed on a Nu-PAGE 4-12% Bis Tris gel (Invitrogen) and visualized via Coomassie staining using the colorimetric channel on a LI-COR Odyssey imager.

*Native PAGE*
His-cleaved BPTF-PHD-BRD, 58 μM, in phosphate buffer (50mM $Na_2HPO_4$, 100 mM NaCl, pH 7.4), was briefly vortexed, and an aliquot was removed. The remaining BPTF dual domain solution was concentrated to 215 μM using an Amicon Ultra-0.5 mL 10 kDa centrifugal filter (Millipore Sigma,UFC501096). PBS buffer, protein, and peptide were sequentially added to the samples. Samples were pipette mixed, manually vortexed, and incubated at room temperature for 30 min. To the samples, 50% glycerol in 1:6 ratio was added and manually vortexed. Concentrated samples were further diluted with 1:6 50% glycerol to a final protein concentration of 20 μM and 5.5 μg of protein was loaded in each well. Samples were manually vortexed and immediately loaded into prewashed wells of a Novex 4-12%, Tris-Glycine Plus WedgeWell Gel (XP04120). Running buffer (1x Novex Tris-Glycine Native Running Buffer pH 8.3, Invitrogen LC2672) was added and the gel was run at 225 V for 55 min at room temperature. During the last 15 min the gel tank was placed on ice to cool the buffer back to RT. The gel was stained with Coommassie Brilliant Blue R-250 Bio-Rad (Cat # 1610400) and destained before imaging at 700 nm on a LI-COR Odyssey imager.

*Size exclusion chromatography coupled small angle X-ray scattering (SEC-SAXS)*
SAXS experiments were performed at the SIBYLS beamline 12.3.1 at the Advanced Light Source using the Mail-in program. This beamline has additional inline instrumentation and detectors coupled to a size exclusion column.[11,12] BPTF-PHD-BRD was exchanged into SAXS

buffer (50 mM Tris, 100 mM NaCl, 2 mM TCEP, 1% glycerol, pH = 7.5), and two samples were prepared: one with BPTF-PHD-BRD alone and another containing BPTF-PHD-BRD with peptide 3. For both samples, 70 µl of 7 mg/mL BPTF-PHD-BRD was suspended in SAXS buffer, which was also used as the SEC running buffer. The X-ray wavelength was set to 1.127 Å and the sample-to-detector distance to 2,100 mm, giving scattering vectors (q) ranging from 0.01 $Å^{-1}$ to 0.4 $Å^{-1}$. The SAXS flow cell was coupled to an Agilent 1260 Infinity HPLC system using a Shodex KW-802.5 SEC column equilibrated with the SEC running buffer, as indicated above, with a flow rate of 0.65 ml/min. The eluent was subject to the following: 1) ultra-violet light (UV) at 280 nm, 2) multi-angle light scattering (MALS), 3) quasi-elastic light scattering (QELS), 4) SAXS, and 5) a refractometer. For SAXS measurements, 2 sec X-ray exposures were collected continuously during a 25 min elution. All frames for analyses had one SAXS frame corresponding to the running buffer before the detection of a peak subtracted from each. The radius of gyration (Rg) was calculated for each of the subtracted frames using the Guinier approximation: $I(q) = I(0)\exp(-q^2Rg^2/3)$ with the limits $qRg < 1.3$. The elution peak was compared to the integral of ratios to background and Rg relative to the recorded frame using the program RAW.[13] Uniform Rg values across an elution peak represent a homogeneous sample. Final merged SAXS profiles, derived by integrating multiple frames at the elution peak, were used for further analyses. The Guinier plot to provide information on the aggregation state, the volume of correlation (Vc) to estimate the molecular weight,[14] and the pair distribution function [P(r)] to calculate the maximal inter-particle dimension.[15]

*MutliFoXS modeling of SAXS data*
Multi-state modeling of the different conformational poses of BPTF-PHD-BRD in both conditions (no peptide versus bound to peptide 3) was performed using the MultiFoXS server. 2F6J was used as the input PDB with chains B and C, as well as the H3(1-15)K4$me_3$ co-crystallized ligand, removed from the input file to recapitulate a monomeric representation of chain A. Residues V65 through L69 were designated as flexible. SAXS scattering data containing q, I(q), and intensity error was used to model BPTF-PHD-BRD populations between both SAXS conditions, generated by sampling 10,000 conformations. Output .pdb files were visualized in UCSF Chimera (v 1.17.3).

*Limited Proteolysis Mass Spectrometry (LiP-MS)*
LiP-MS experiments were performed based on previously described methods.[16,17] Reactions were prepared containing 78 µM BPTF-PHD-BRD in 50 mM $Na_2HPO_4$, 100 mM NaCl, 2 mM TCEP, pH 7.4 alone or in the presence of equimolar concentrations of peptide **3**, with each condition tested in triplicate. Reactions were incubated for 30 minutes at room temperature. Then, reactions were incubated with proteinase K (Sigma-Aldrich; 1:100 ratio proteinase K:BPTF-PHD-BRD) for 1 minute at room temperature and immediately boiled at 95 °C to inactivate proteinase K. Sodium deoxycholate (DOC; Sigma-Aldrich) was added to reactions at a final concentration of 2% (vol/vol). Then, DTT (20 mM final) was added to reactions and incubated at 60°C for 30 minutes, followed by Iodoacetamide (40 mM final) and further incubated for 30 mins at room temperature away from light. Then, Lys-C/Trypsin protease cocktail (Thermo Fischer Scientific) was added (1:20 ratio protease cocktail:BPTF-PHD-BRD) and reactions were incubated overnight at 37 °C. Samples were brought to room temperature and DOC was precipitated by addition of 2% formic acid (FA; vol/vol), followed by centrifugation at 16,000 xg for 10 minutes. The supernatant was collected and samples were desalted using Sep-

Pak C18 cartridges (Waters). Samples were concentrated via speed vacuum centrifugation to yield dried peptide mixtures that were dissolved into liquid chromatography mass spectrometry grade $H_2O$ supplemented with 0.1% FA for subsequent LC-MS analysis.

*LC-MS conditions for LiP-MS*

LC-MS was performed on a QExactive Orbitrap Hybrid Mass Spectrometer interfaced with an UltiMate 3000 UHPLC system run in nanoflow mode at 0.3 µL/min. A nanoLC column was packed with Luna C18 stationary phase and peptides were separated using 0.1% FA in LC-MS $H_2O$ (buffer A) and 0.1% FA in acetonitrile (buffer B). Peptides were separated by a 5-50% gradient of buffer B spanning 45 minutes, followed by a 50-95% gradient of buffer B spanning 5 minutes and a 5 minute wash at 95% buffer B, and completed by a 95-5% equilibration for 5 minutes. Peptides were analyzed in positive ion mode using a Top12 Full MS/dd-$MS^2$ experiment. Full MS resolution was set to 70,000 $MS^1$ resolution with an AGC target of 1 x $10^6$, maximum IT of 30 ms, and a scan range of 300 to 2000 *m/z*. Tandem mass spectra were collected at 17,500 $MS^2$ resolution, AGC target of 5 x $10^4$, maximum IT of 50 ms, and isolation window of 2.0 *m/z*, and normalized collision energy of 30%. Data were collected in centroid mode. The resulting .RAW files were used for subsequent data analysis.

*LiP-MS data processing*

FragPipe (v 22.0), with MSFragger[18] and IonQuant[19] imbedded workflows, was used to analyze LC-MS data. The label free quantification match-between-runs (LFQ-MBR) workflow supporting data dependent acquisition was selected. Peptides were searched against a database generated from a FASTA file containing the amino acid sequence for BPTF-PHD-BRD used in this study, with decoys added. Generally, default settings were used including those within IonQuant to generate LFQ intensities.[20] Within MSFragger, the peptide digest pattern was set to nonspecific with up to two missed cleavages allowed. Peptide length was limited to capture sequences of 7-25 amino acids. Variable modifications of methionine oxidation and cysteine carbamidomethylation were included. Peptides, and their corresponding LFQ intensities, were further analyzed in Perseus (v 2.0.9.0) to analyze changes in abundance between conditions. Peptides were filtered to only include those with replicate LFQ intensities in a single condition, and remaining intensities were Log2 transformed. A two-tailed, two sample *t*-test was performed to identify peptide sequences that underwent statistically significant differential regulation between the unliganded controls and samples bound to ligand **3**. Statistical significance was determined using a Benjamini-Hochberg corrected FDR of 0.05 and a minimal coefficient of variation (S0) of 0.1 and proteinase K cleavage sites were mapped onto structures using UCSF Chimera (v 1.17.3).

Ethical use of animals or human participants in research: No animal or human participants used for this work.

Safety: No unexpected, new, and/or significant hazards or risks were associated with this work.

**Supplementary Table 1.** Masses (in Da) obtained using MALDI-TOF for H3 peptides and bivalent inhibitors, with doubly charged ions labeled if observed. Kme$_3$ = trimethyllysine; Kac = acetyllysine; Ahx = 6-aminohexanoic acid.

| | Sequence | Exp. $[M]^+$ or $[M+H]^+$ | Obs. $[M]^+$ or $[M+H]^+$ | Exp. $[M+H]^{2+}/2$ or $[M+2H]^{2+}/2$ | Obs. $[M+H]^{2+}/2$ or $[M+2H]^{2+}/2$ |
|---|---|---|---|---|---|
| 1 | ARTKme$_3$QTARKSTGGK-Ahx-Y | 1878.103 | 1878.446 | - | - |
| 2 | ARTKme$_3$QTARKacSTGGKac-Ahx-Y | 1961.129 | 1962.143 | - | - |
| 3 | ARTKme$_3$QTARKacSTGGKacAPRKacQL-Ahx-Y | 2626.526 | 2626.508 | 1313.767 | 1314.275 |
| 4 | ARTKQTARKacSTGGKacAPRKacQL-Ahx-Y | 2584.479 | 2585.000 | - | - |
| 5 | ARTKme$_3$QTARKSTGGKAPRKQL-Ahx-Y | 2501.996 | 2500.986 | 1250.751 | 1251.013 |
| 6 | ARTKme$_3$QTARKacSTGGKacAPRKQL-Ahx-Y | 2584.516 | 2584.958 | - | - |
| 7 | ARTKme$_3$QTARKSTGGKacAPRKacQL-Ahx-Y | 2584.516 | 2584.484 | 1292.762 | 1293.242 |
| 8 | ARTKme$_3$QTARKacSTGGKAPRKQL-Ahx-Y | 2542.505 | 2542.536 | 1271.756 | 1272.308 |
| 9 | ARTKme$_3$QTARKSTGGKacAPRKQL-Ahx-Y | 2542.505 | 2541.867 | 1271.756 | 1271.407 |
| 10 | ARTKme$_3$QTARKSTGGKAPRKacQL-Ahx-Y | 2542.505 | 2543.132 | 1271.756 | 1272.089 |
| 11 | ARTKQTARKSTGGKacAPRKQL-Ahx-Y | 2500.458 | 2500.498 | 1250.733 | 1250.782 |
| 12 | ARTKQTARKSTGGKAPRKQL-Ahx-Y | 2458.448 | 2458.634 | 1229.728 | 1230.345 |
| | MJ-1 | 1180.660 | 1180.646 | | |
| | MJ-2 | 1428.786 | 1428.748 | | |

**Analytical HPLC traces for H3 peptides and Bivalent Inhibitors:**

Blank

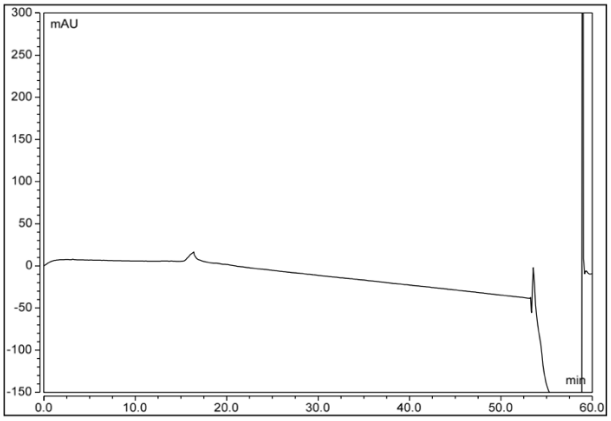

1.) ARTKme$_3$QTARKSTGGK-Ahx-Y

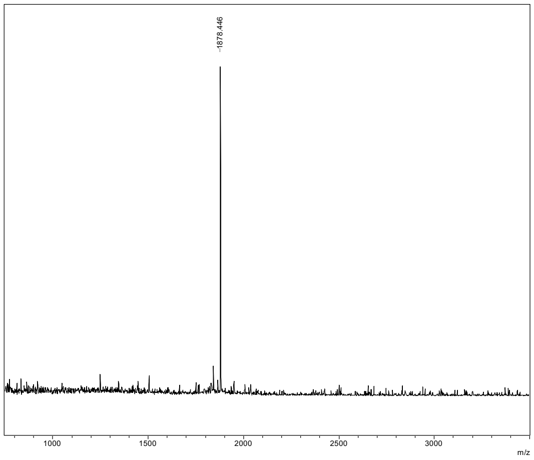

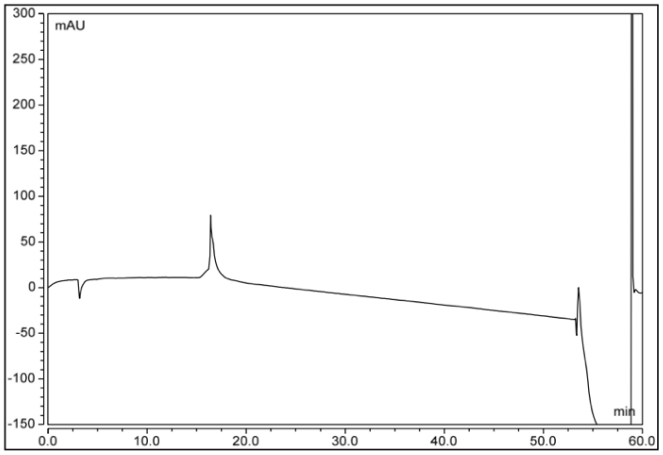

2.) ARTKme$_3$QTARKacSTGGKac-Ahx-Y

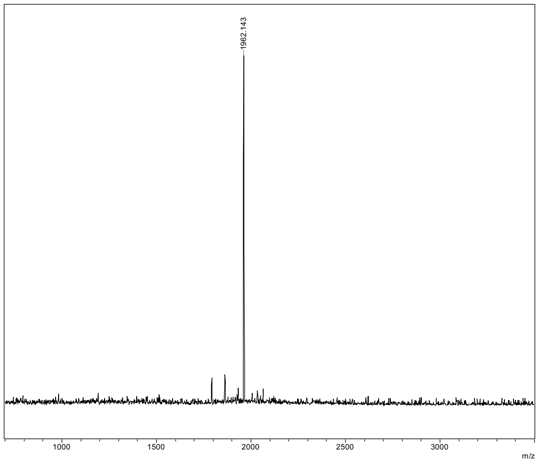

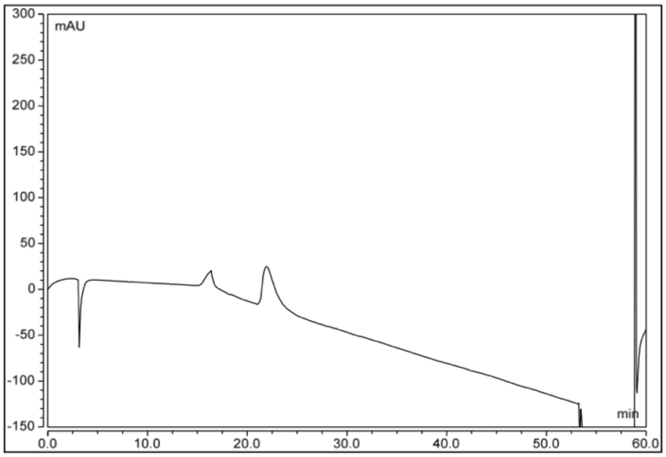

3.) ARTKme$_3$QTARKacSTGGKacAPRKacQL-Ahx-Y

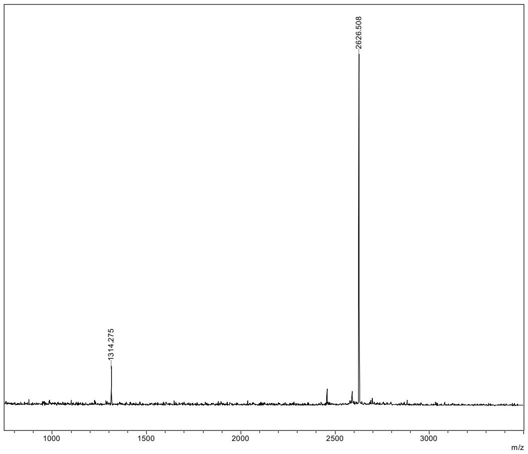

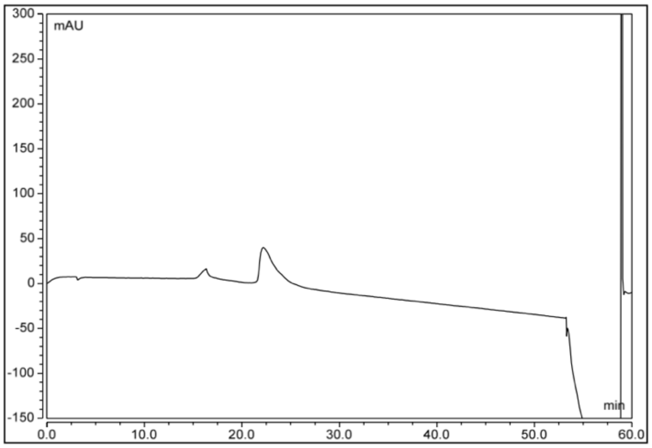

4.) ARTKQTARKacSTGGKacAPRKacQL-Ahx-Y

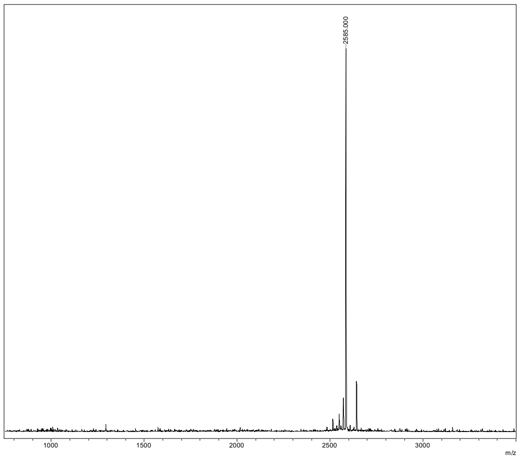

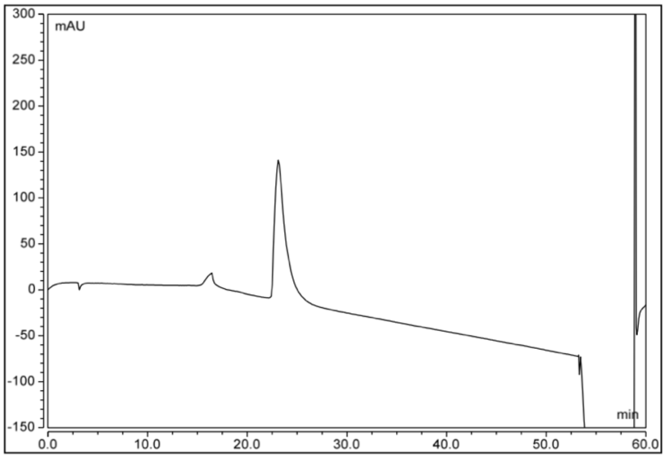

5.) ARTKme$_3$QTARKSTGGKAPRKQL-Ahx-Y

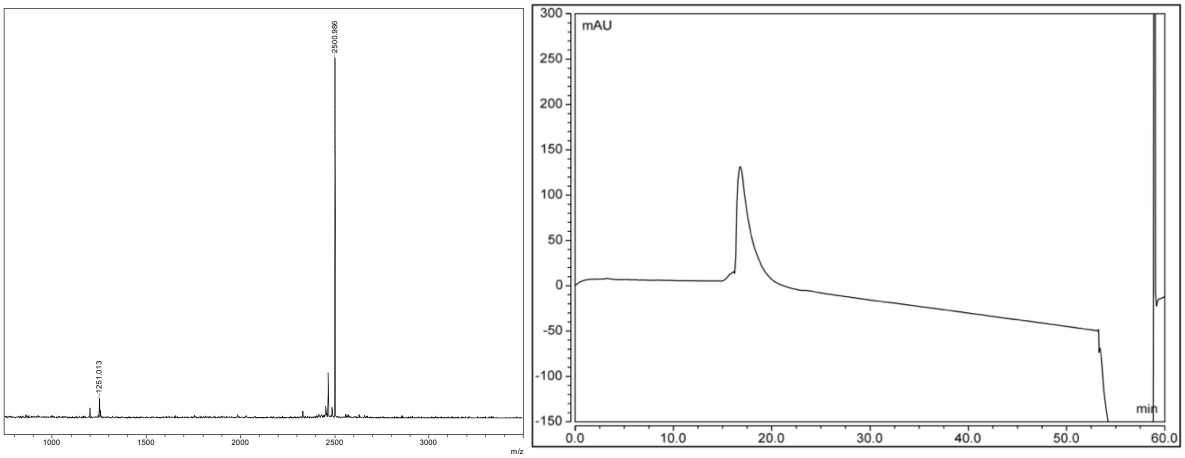

6.) ARTKme$_3$QTARKacSTGGKacAPRKQL-Ahx-Y

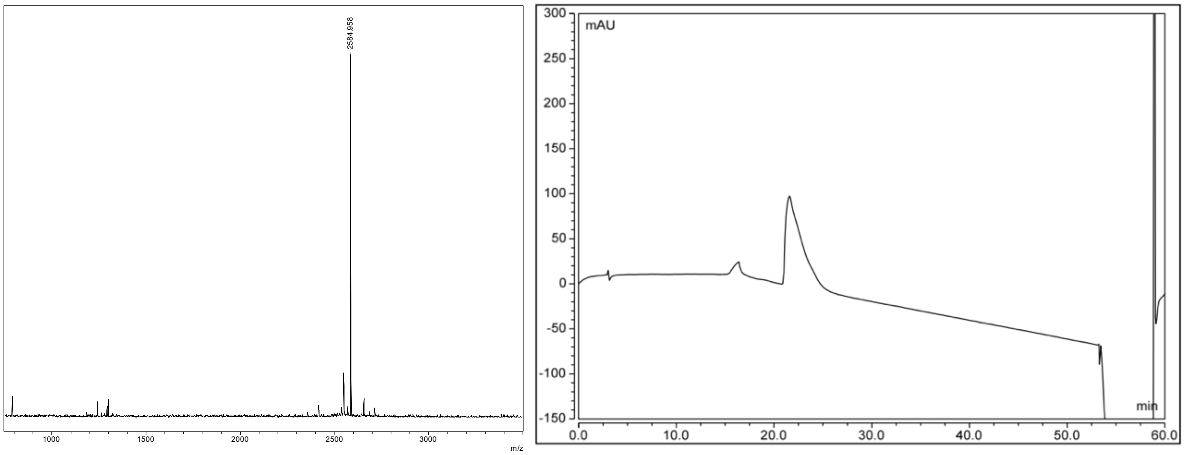

7.) ARTKme$_3$QTARKSTGGKacAPRKacQL-Ahx-Y

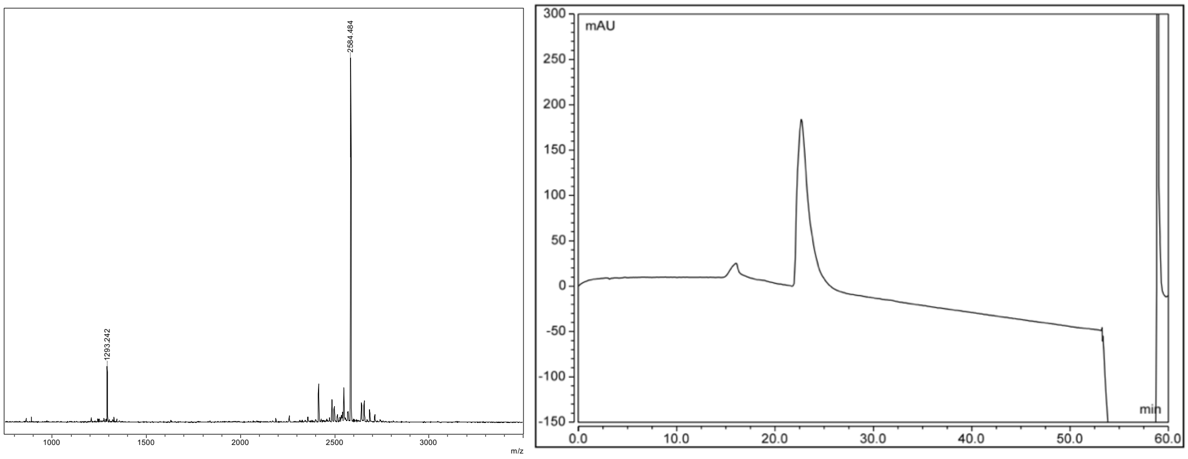

8.) ARTKme$_3$QTARKacSTGGKAPRKQL-Ahx-Y

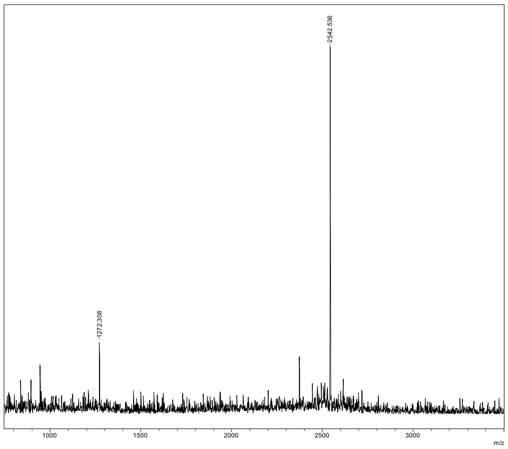

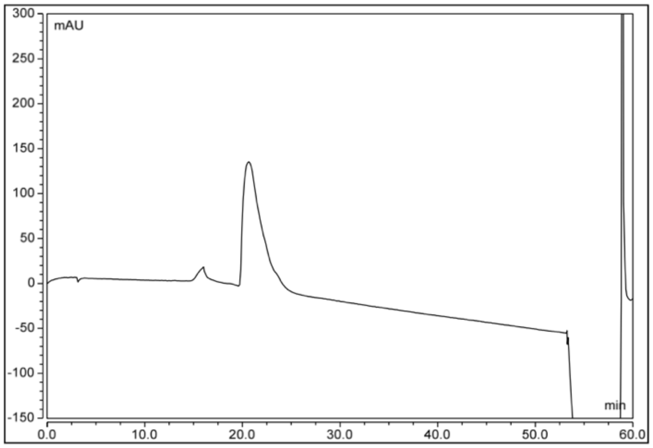

9.) ARTKme$_3$QTARKSTGGKacAPRKQL-Ahx-Y

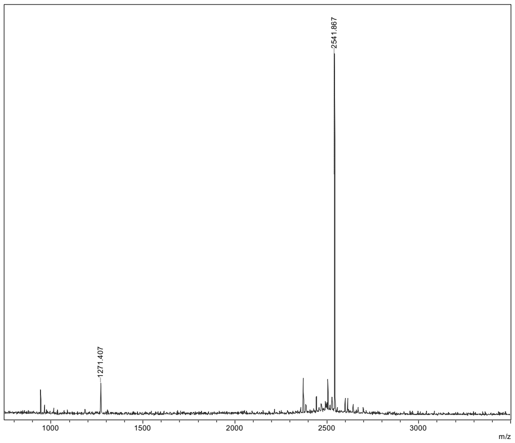

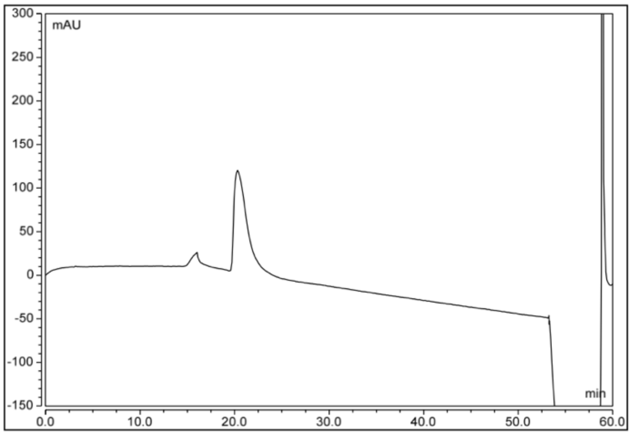

10.) ARTKme$_3$QTARKSTGGKAPRKacQL-Ahx-Y

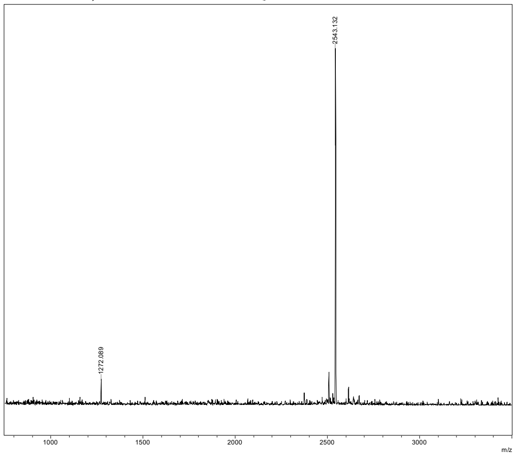

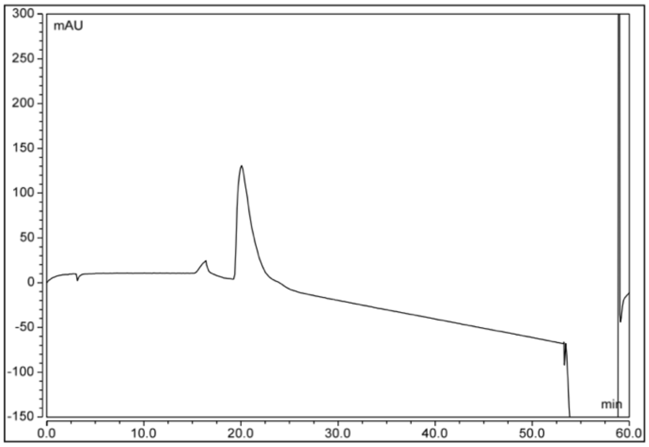

11.) ARTKQTARKSTGGKacAPRKQL-Ahx-Y

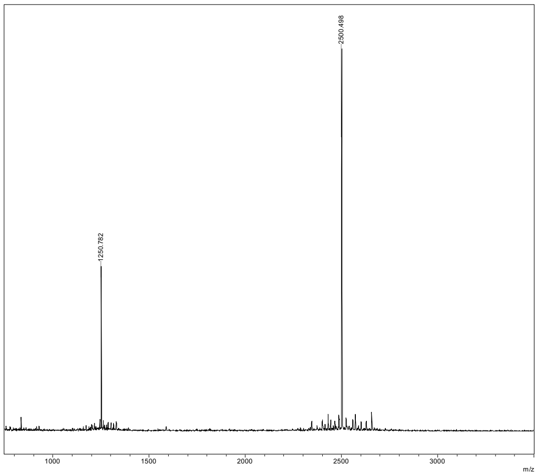

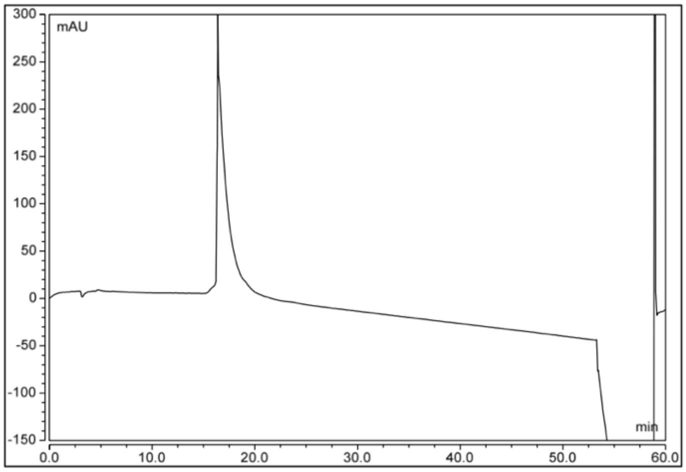

12.) ARTKQTARKSTGGKAPRKQL-Ahx-Y

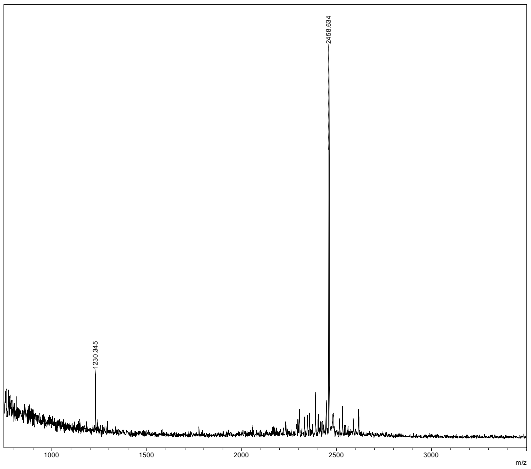

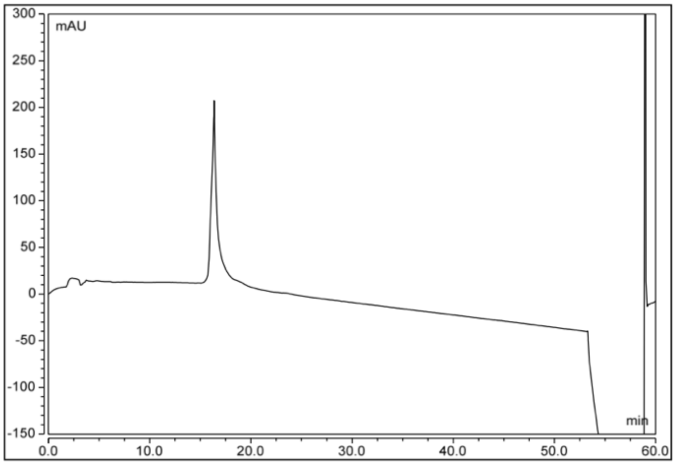

MJ-1

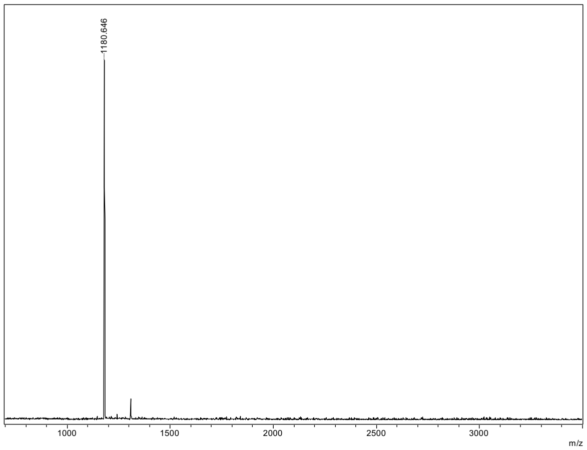

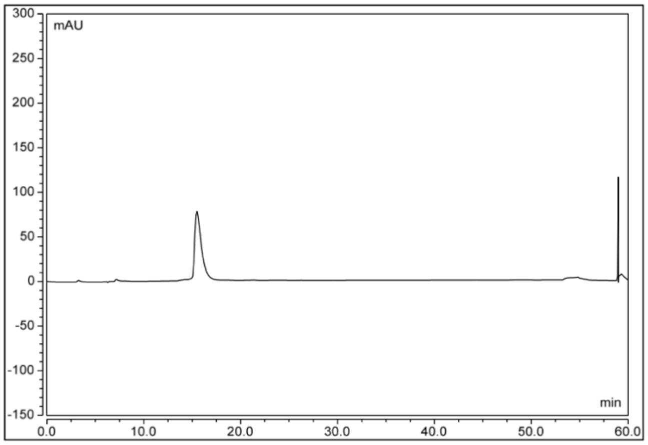

MJ-2

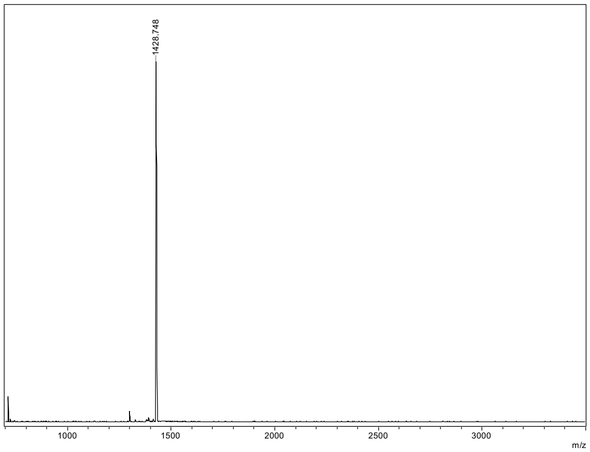

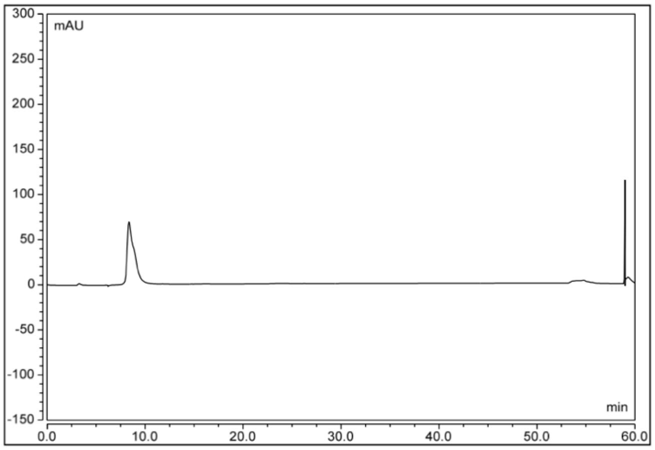

### [1]H and [13]C Spectra

#### 14, [1]H NMR (500 MHz, chloroform-d)

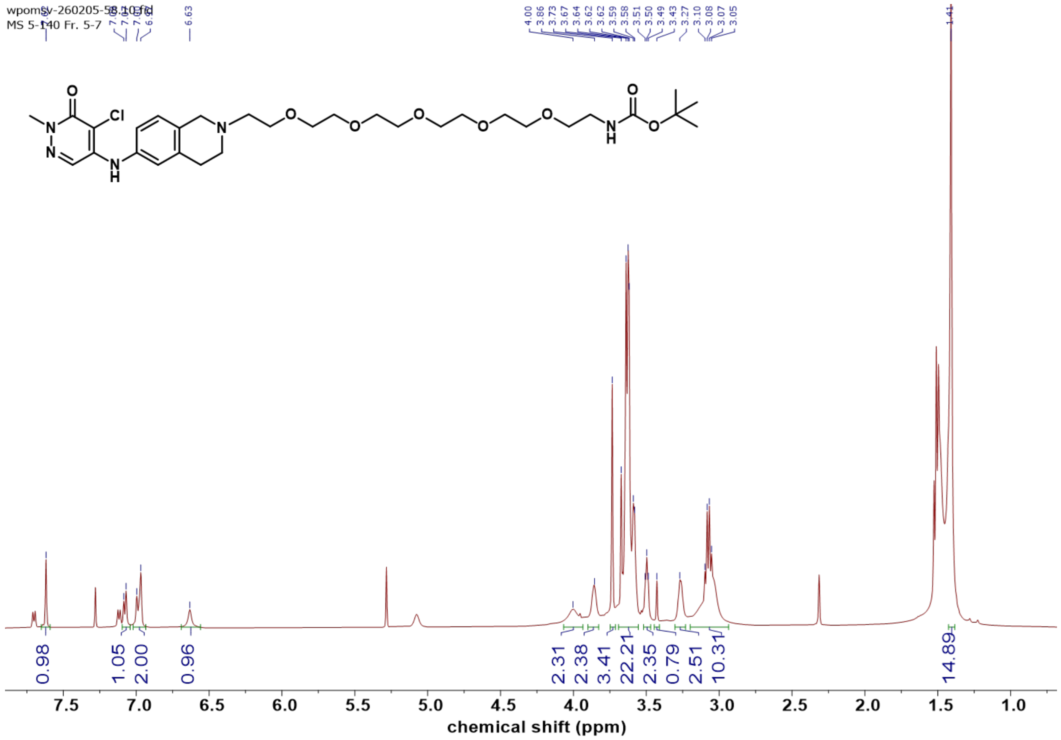

#### 14, [13]C NMR (126 MHz, chloroform-d)

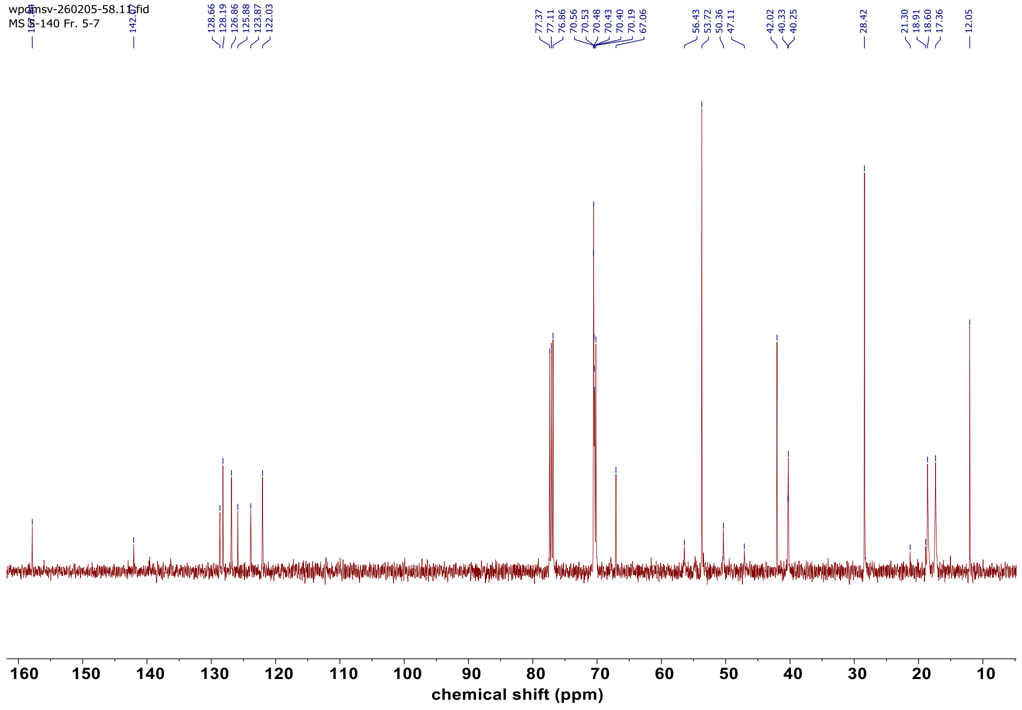

#### DSF Melt Curves for WT and Mutant BPTF-PHD-BRD

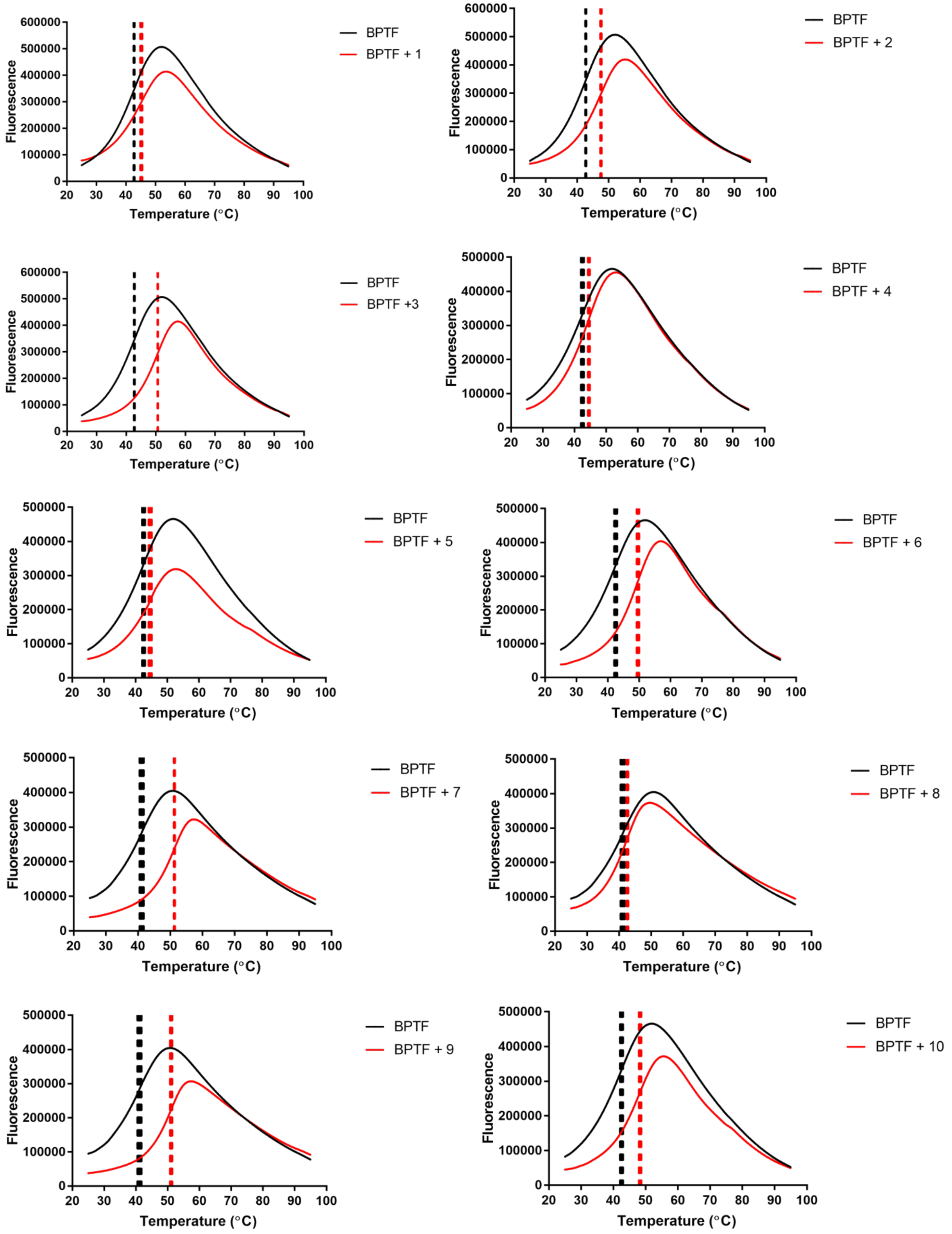

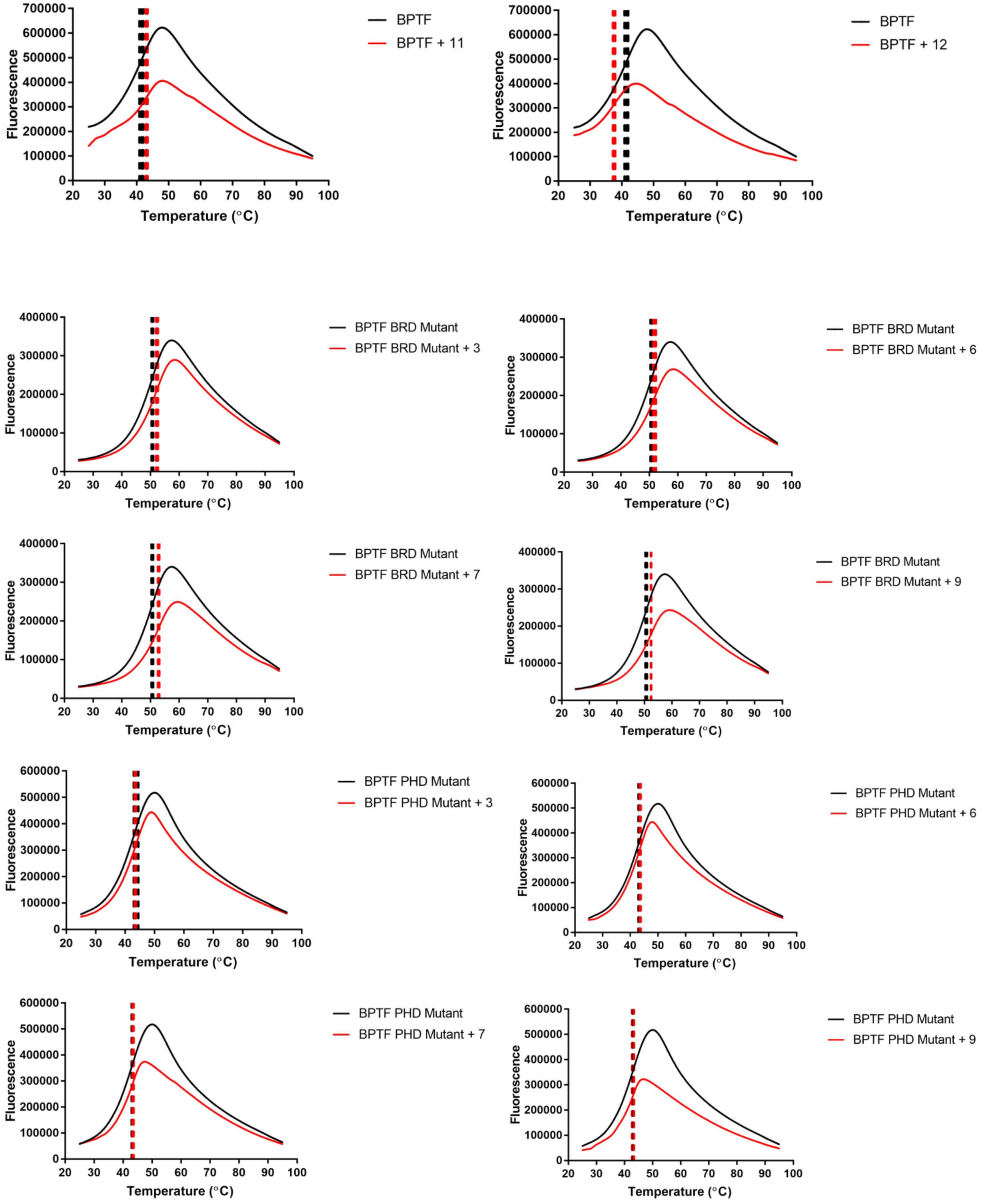

Fluorescence
Temperature (°C)
BPTF
BPTF + 11
BPTF + 12
BPTF BRD Mutant
BPTF BRD Mutant + 3
BPTF BRD Mutant + 6
BPTF BRD Mutant + 7
BPTF BRD Mutant + 9
BPTF PHD Mutant
BPTF PHD Mutant + 3
BPTF PHD Mutant + 6
BPTF PHD Mutant + 7
BPTF PHD Mutant + 9

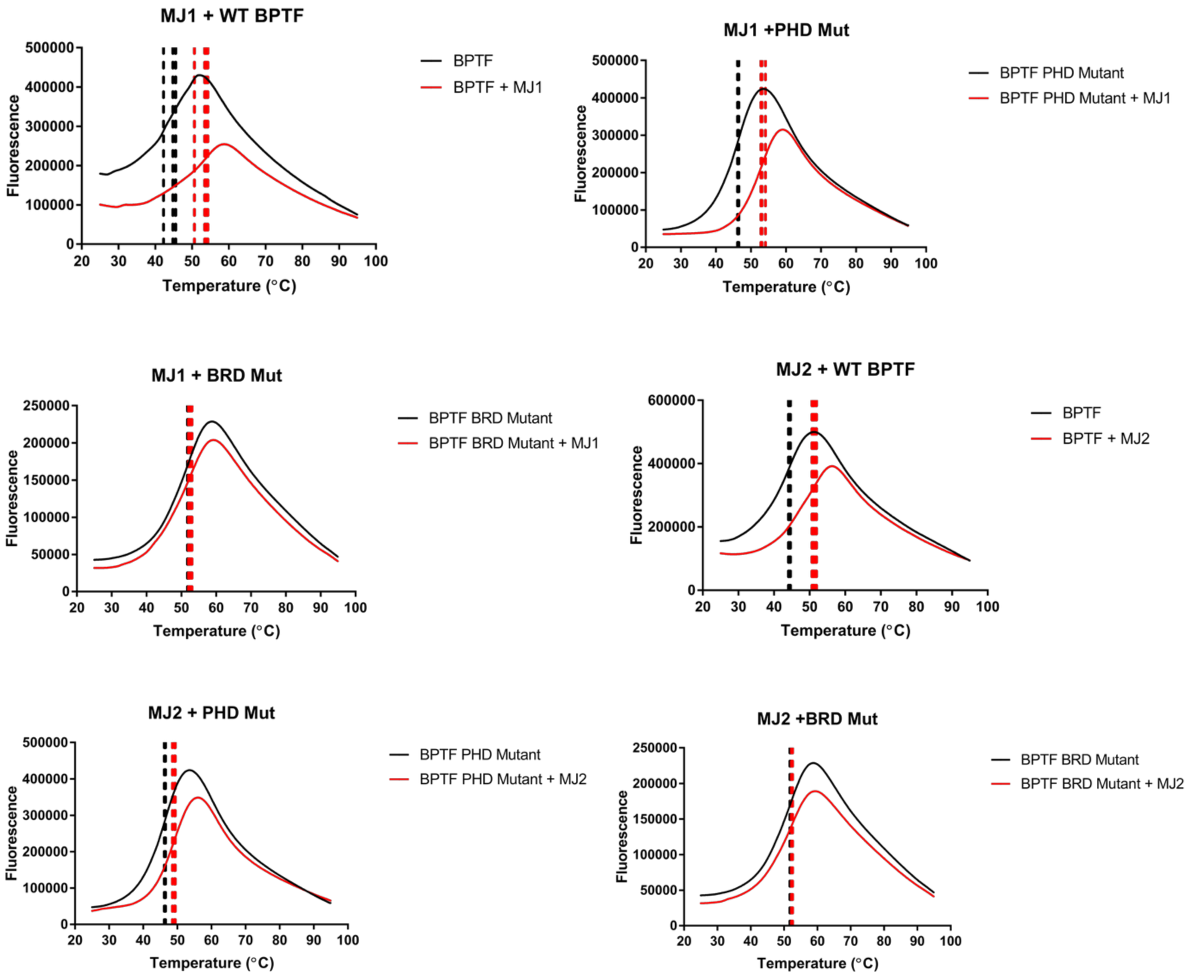
MJ1 + WT BPTF
BPTF
BPTF + MJ1
MJ1 +PHD Mut
BPTF PHD Mutant
BPTF PHD Mutant + MJ1
MJ1 + BRD Mut
BPTF BRD Mutant
BPTF BRD Mutant + MJ1
MJ2 + WT BPTF
BPTF
BPTF + MJ2
MJ2 + PHD Mut
BPTF PHD Mutant
BPTF PHD Mutant + MJ2
MJ2 +BRD Mut
BPTF BRD Mutant
BPTF BRD Mutant + MJ2
Fluorescence
Temperature (°C)

#### Alphascreen™ PHD Competition Replicates with WT BPTF-PHD-BRD

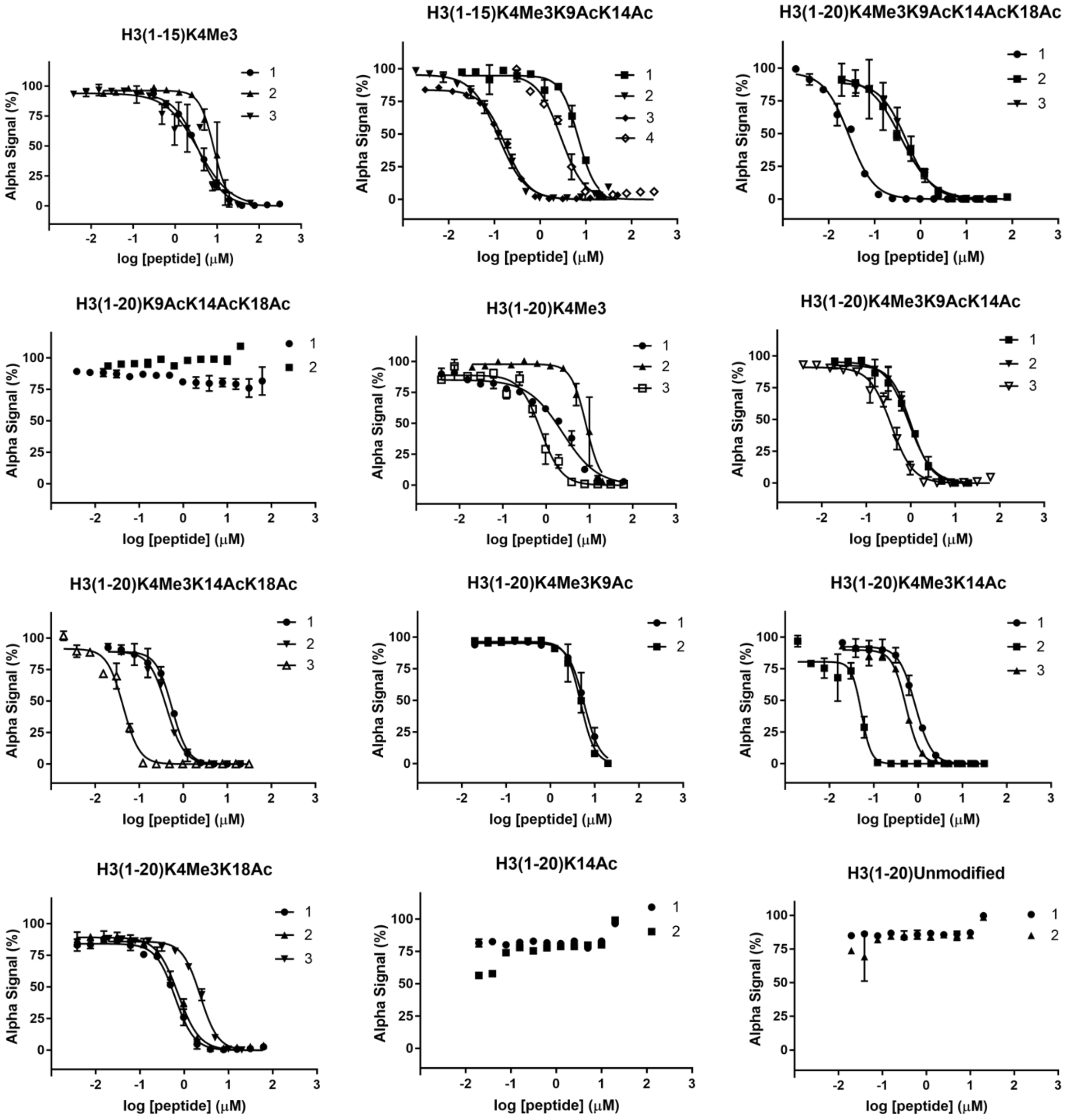

#### Alphascreen™ BRD Competition Replicates with WT BPTF-PHD-BRD

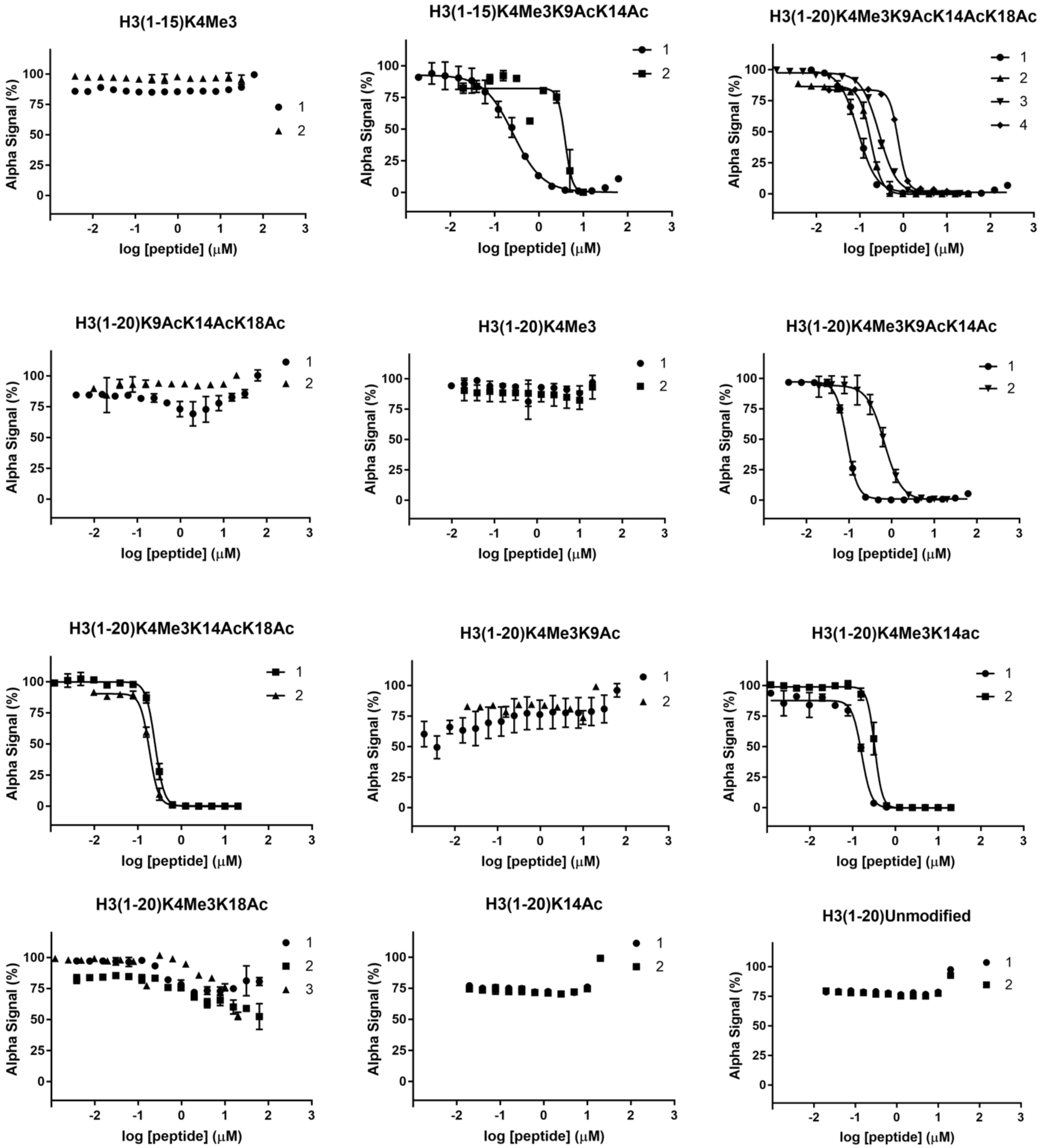

#### Alphascreen™ PHD Competition Replicates with BPTF-PHD-BRD BRD Mutants

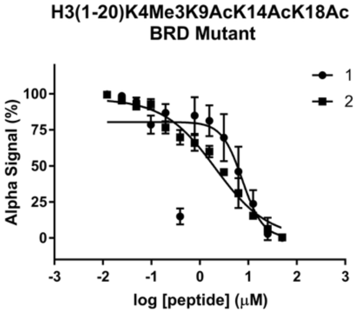

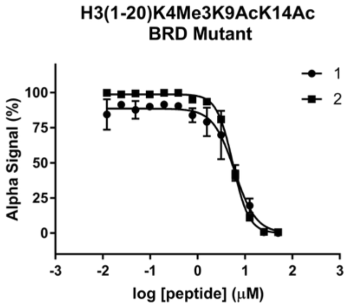

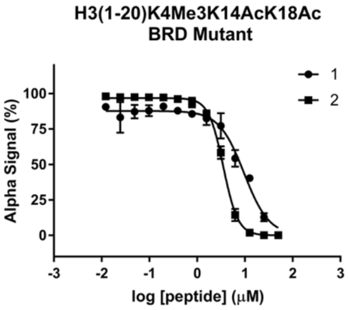

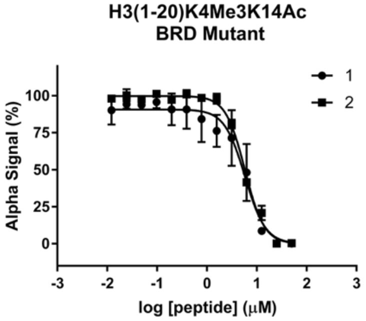

#### Alphascreen™ BRD and PHD Competition Replicates with MJ1 and MJ2

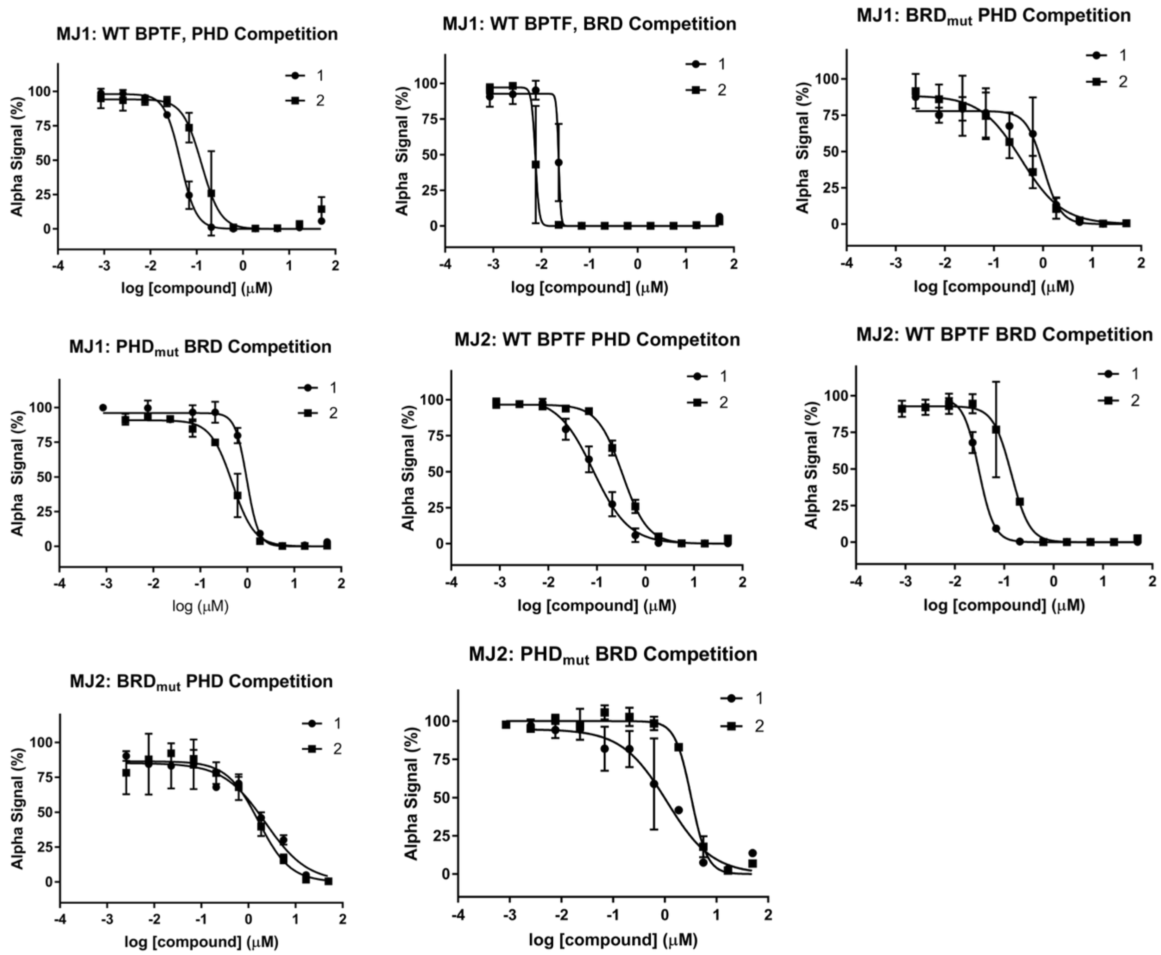

#### Isothermal Titration Calorimetry Binding Isotherms

1: H3(1-15)K4me$_3$

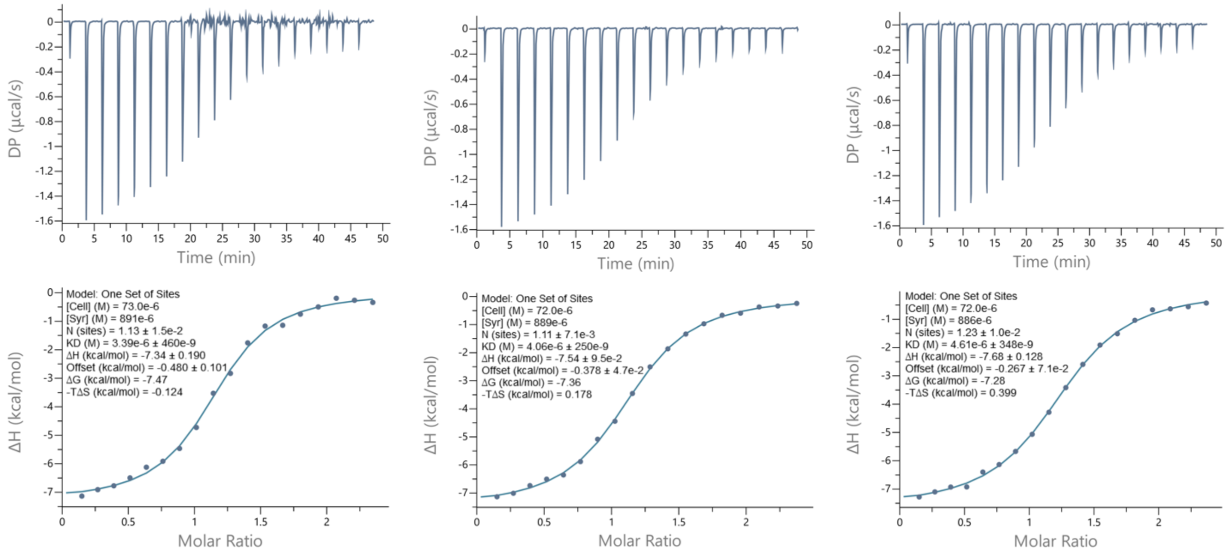

2: H3(1-15)K4me$_3$K9acK14ac

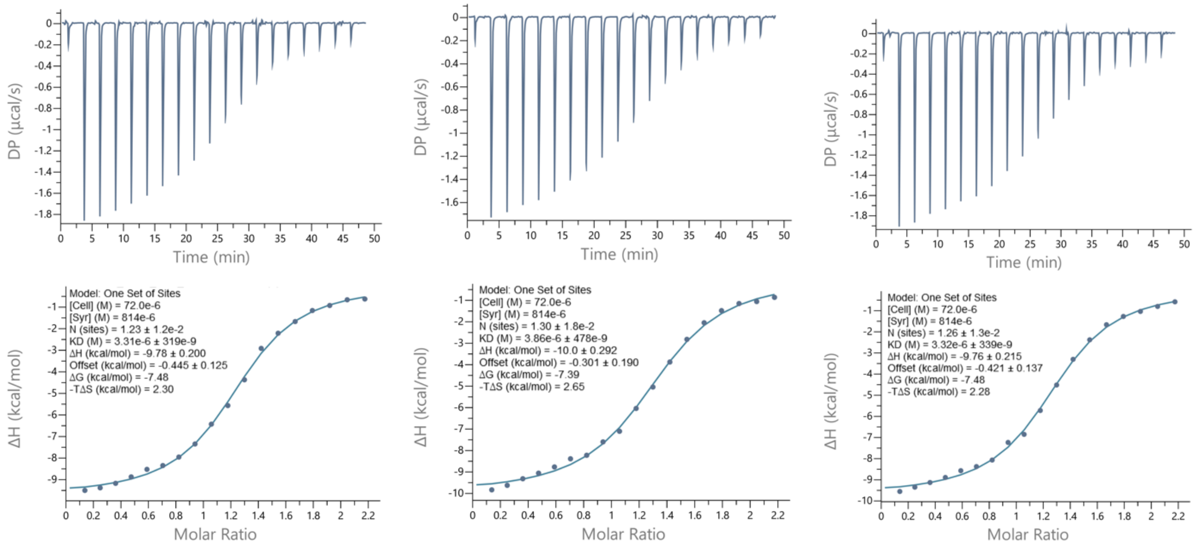

3: H3(1-20)K4me$_3$K9acK14acK18ac

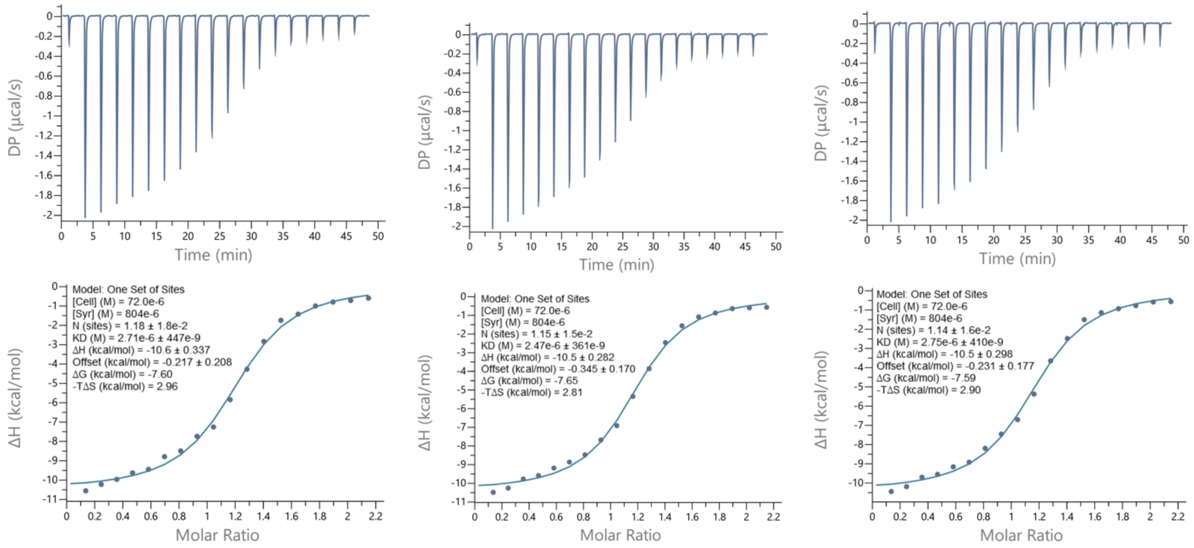

4: H3(1-20)K9acK14acK18ac

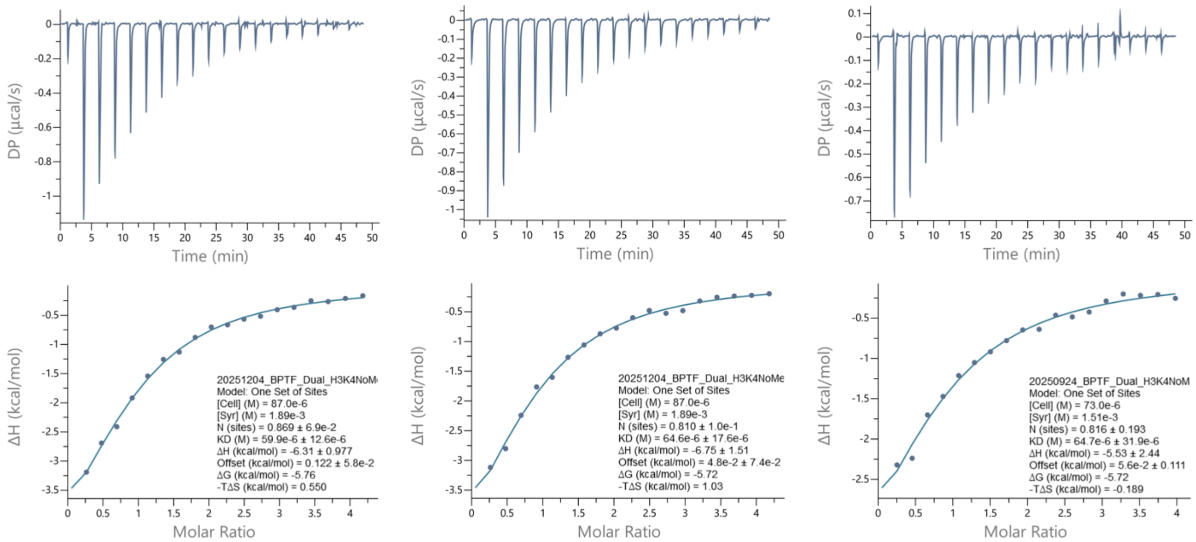

5: H3(1-20)K4me$_3$

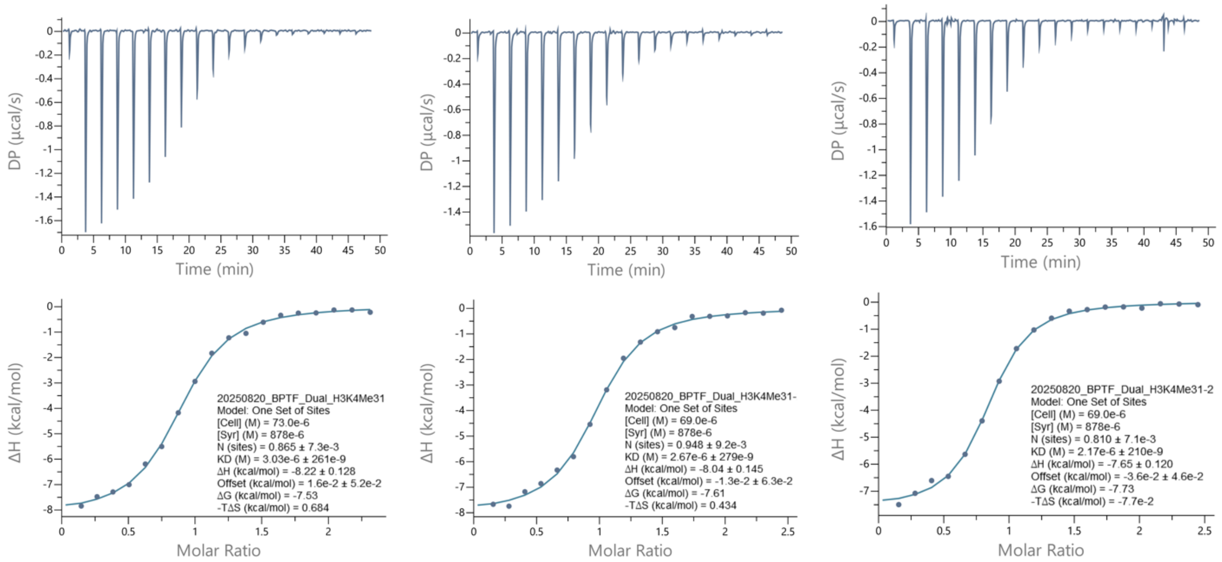

6: H3(1-20)K4me$_3$K9acK14ac

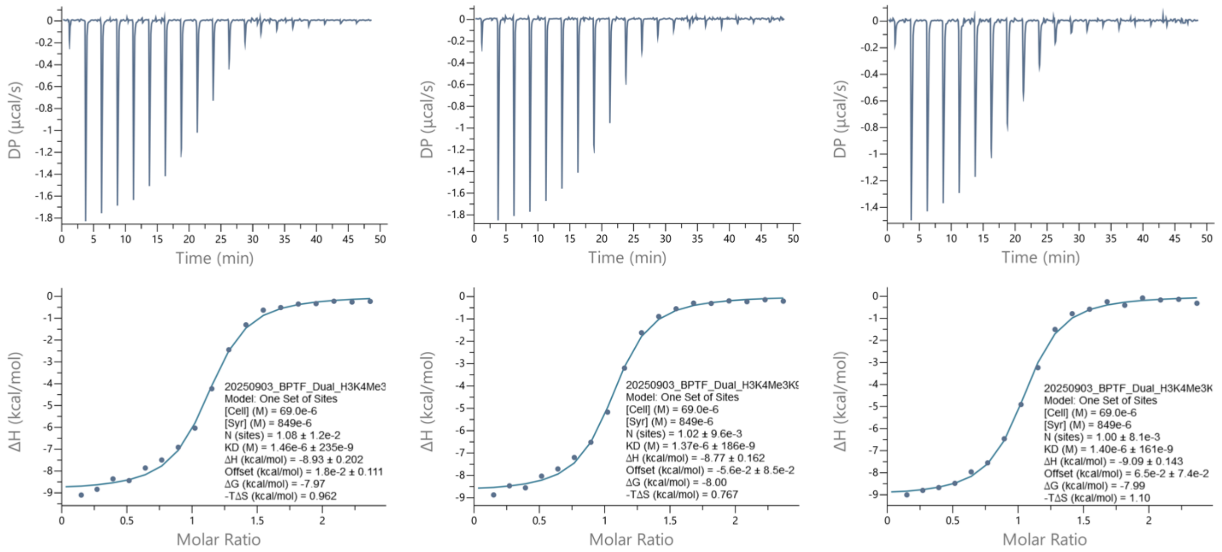

7: H3(1-20)K4me$_3$K14acK18ac

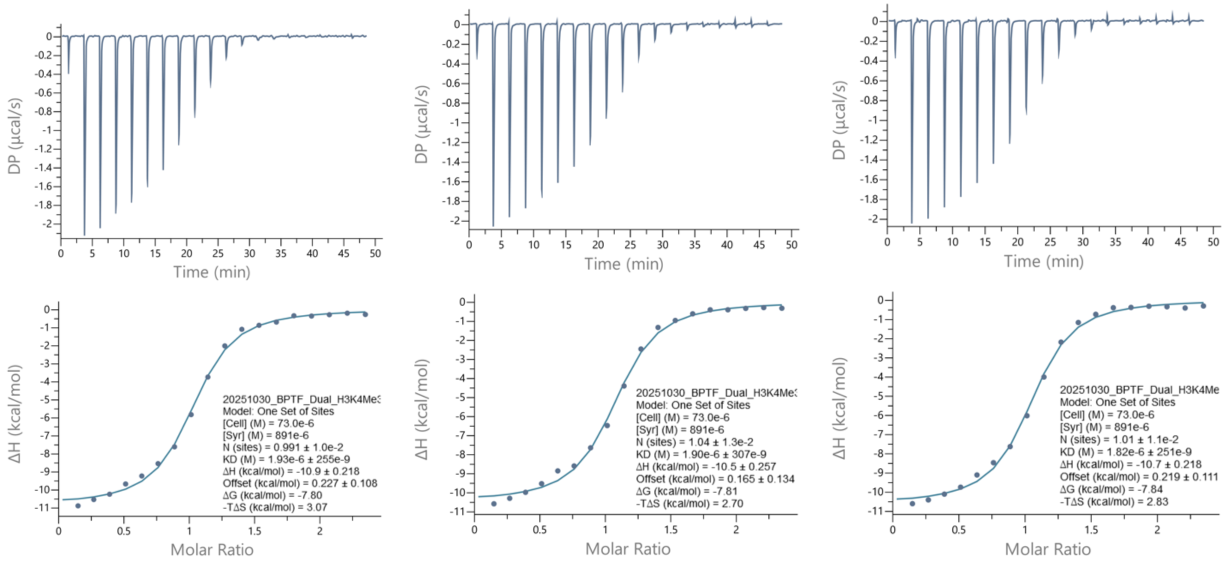

8: H3(1-20)K4me$_3$K9ac

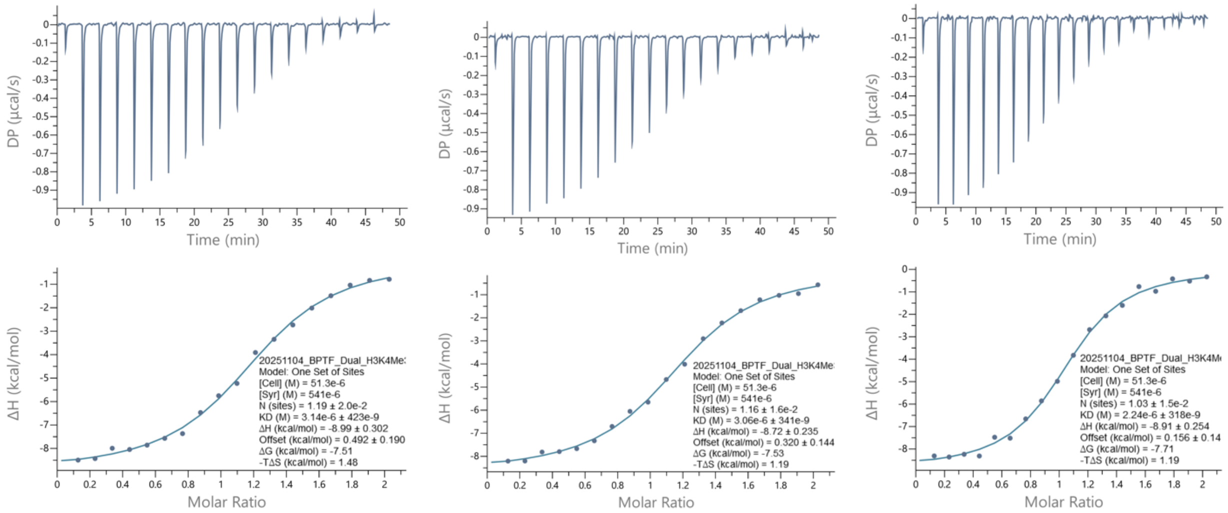

9: H3(1-20)K4me$_3$K14ac

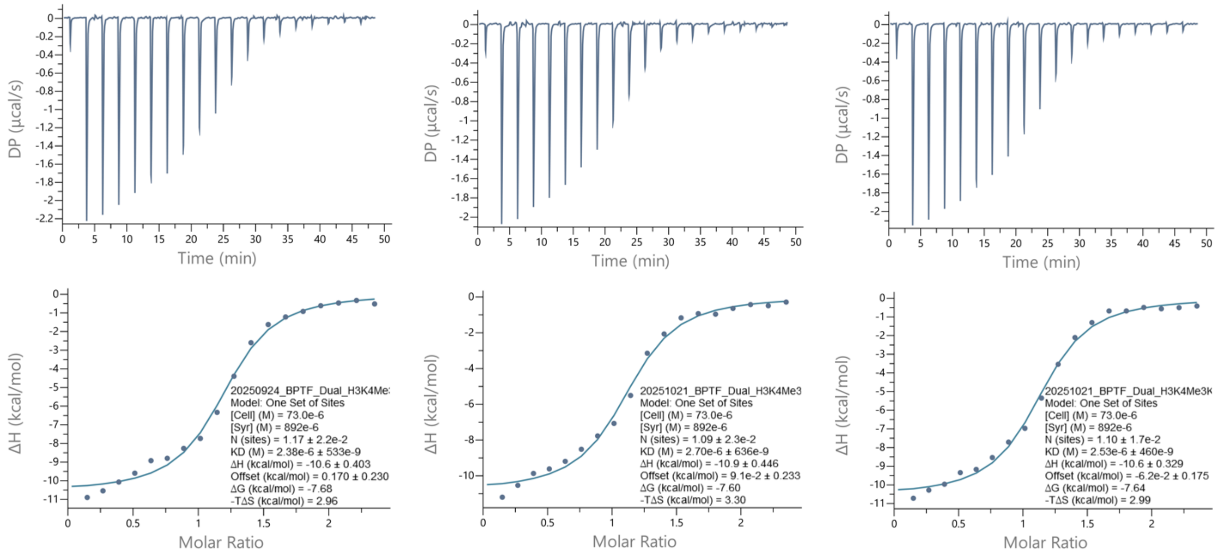

10: H3(1-20)K4me$_3$K18ac

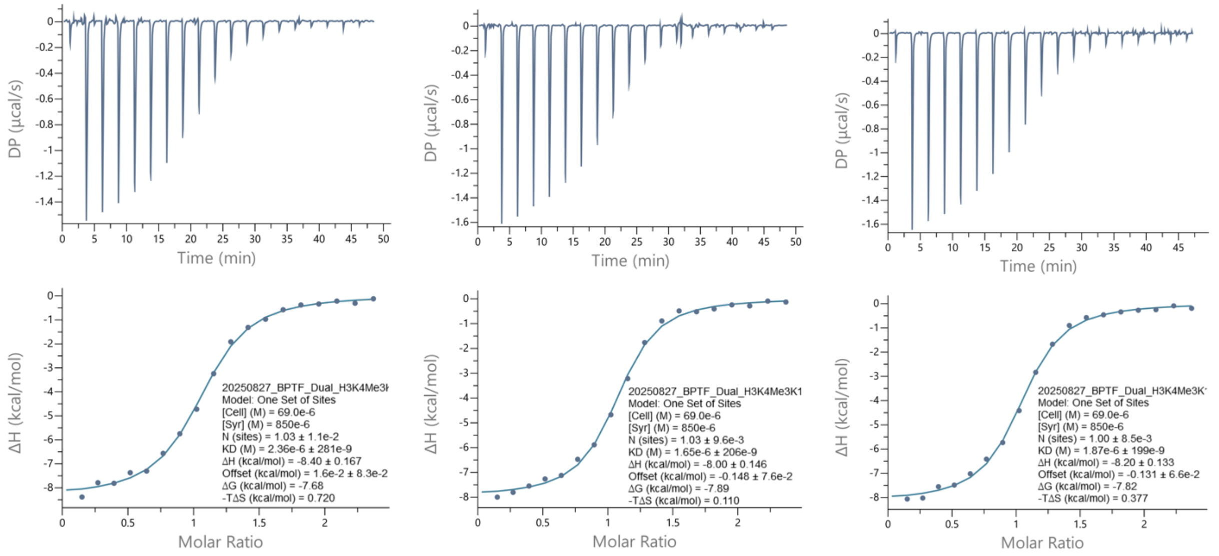

11: H3(1-20)K14ac

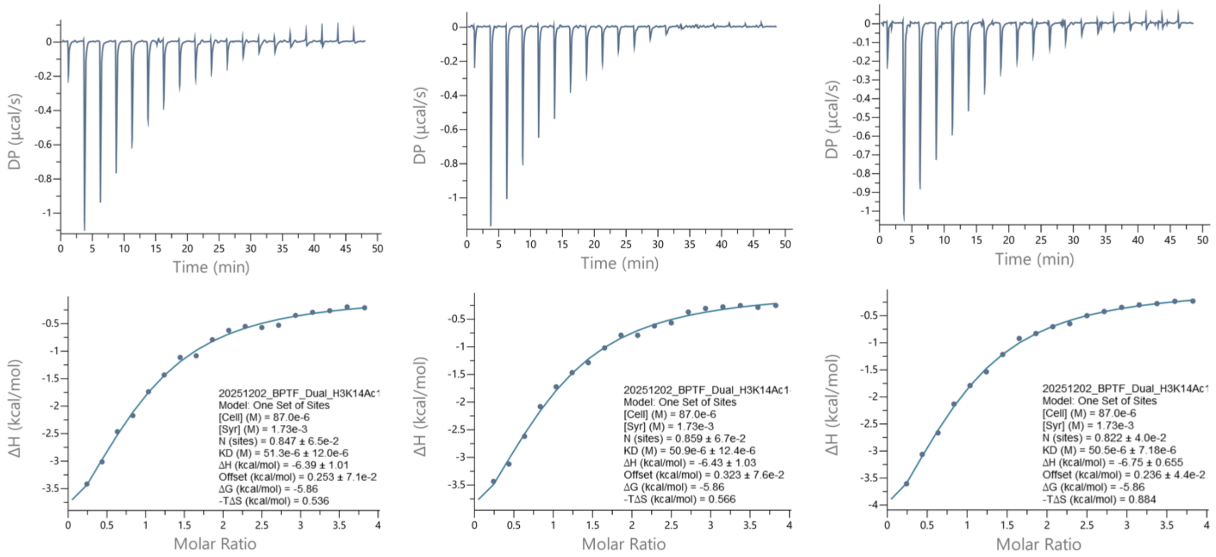

12: H3(1-20)Unmodified

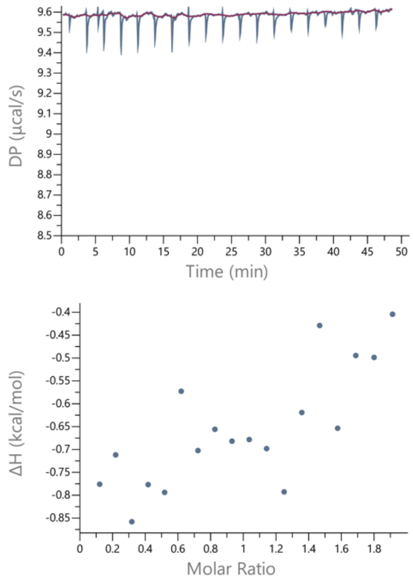

MJ-1:

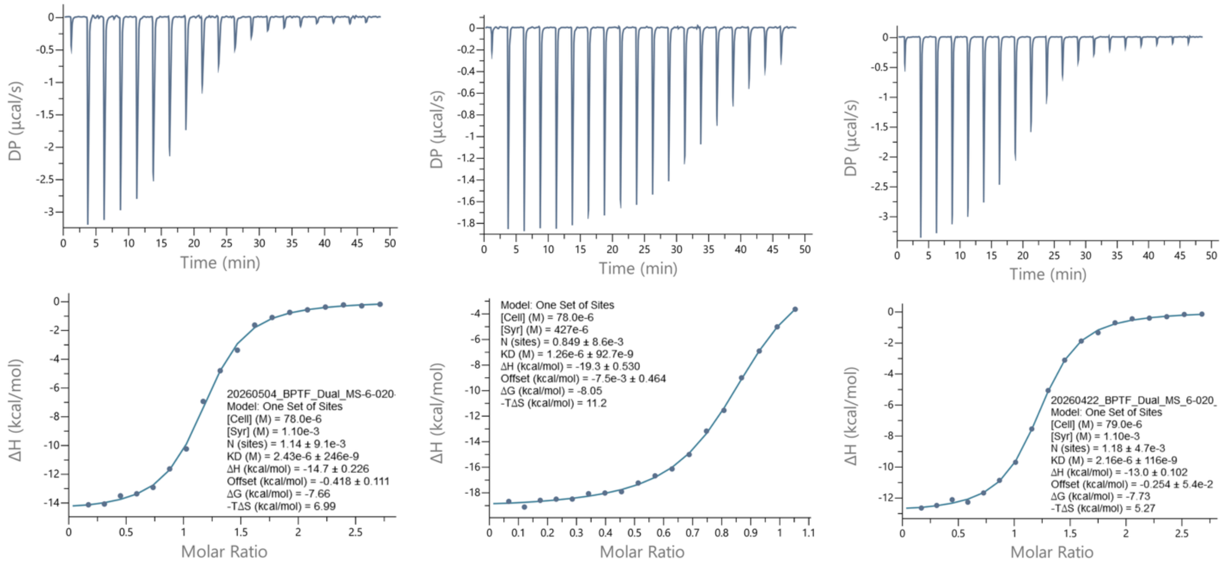

DP (µcal/s)
Time (min)
ΔH (kcal/mol)
Molar Ratio
20260504_BPTF_Dual_MS-6-020-
Model: One Set of Sites
[Cell] (M) = 78.0e-6
[Syr] (M) = 1.10e-3
N (sites) = 1.14 ± 9.1e-3
KD (M) = 2.43e-6 ± 246e-9
ΔH (kcal/mol) = -14.7 ± 0.226
Offset (kcal/mol) = -0.418 ± 0.111
ΔG (kcal/mol) = -7.66
-TΔS (kcal/mol) = 6.99
Model: One Set of Sites
[Cell] (M) = 78.0e-6
[Syr] (M) = 427e-6
N (sites) = 0.849 ± 8.6e-3
KD (M) = 1.26e-6 ± 92.7e-9
ΔH (kcal/mol) = -19.3 ± 0.530
Offset (kcal/mol) = -7.5e-3 ± 0.464
ΔG (kcal/mol) = -8.05
-TΔS (kcal/mol) = 11.2
20260422_BPTF_Dual_MS_6-020_
Model: One Set of Sites
[Cell] (M) = 79.0e-6
[Syr] (M) = 1.10e-3
N (sites) = 1.18 ± 4.7e-3
KD (M) = 2.16e-6 ± 116e-9
ΔH (kcal/mol) = -13.0 ± 0.102
Offset (kcal/mol) = -0.254 ± 5.4e-2
ΔG (kcal/mol) = -7.73
-TΔS (kcal/mol) = 5.27

MJ-2:

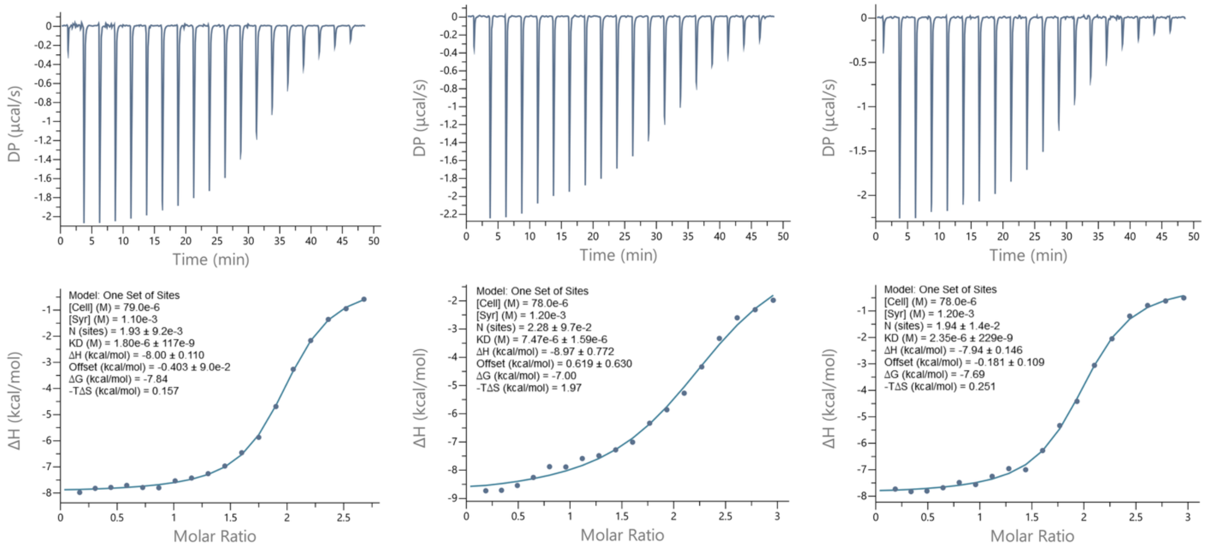

DP (µcal/s)
Time (min)
ΔH (kcal/mol)
Molar Ratio
Model: One Set of Sites
[Cell] (M) = 79.0e-6
[Syr] (M) = 1.10e-3
N (sites) = 1.93 ± 9.2e-3
KD (M) = 1.80e-6 ± 117e-9
ΔH (kcal/mol) = -8.00 ± 0.110
Offset (kcal/mol) = -0.403 ± 9.0e-2
ΔG (kcal/mol) = -7.84
-TΔS (kcal/mol) = 0.157
Model: One Set of Sites
[Cell] (M) = 78.0e-6
[Syr] (M) = 1.20e-3
N (sites) = 2.28 ± 9.7e-2
KD (M) = 7.47e-6 ± 1.59e-6
ΔH (kcal/mol) = -8.97 ± 0.772
Offset (kcal/mol) = 0.619 ± 0.630
ΔG (kcal/mol) = -7.00
-TΔS (kcal/mol) = 1.97
Model: One Set of Sites
[Cell] (M) = 78.0e-6
[Syr] (M) = 1.20e-3
N (sites) = 1.94 ± 1.4e-2
KD (M) = 2.35e-6 ± 229e-9
ΔH (kcal/mol) = -7.94 ± 0.146
Offset (kcal/mol) = -0.181 ± 0.109
ΔG (kcal/mol) = -7.69
-TΔS (kcal/mol) = 0.251

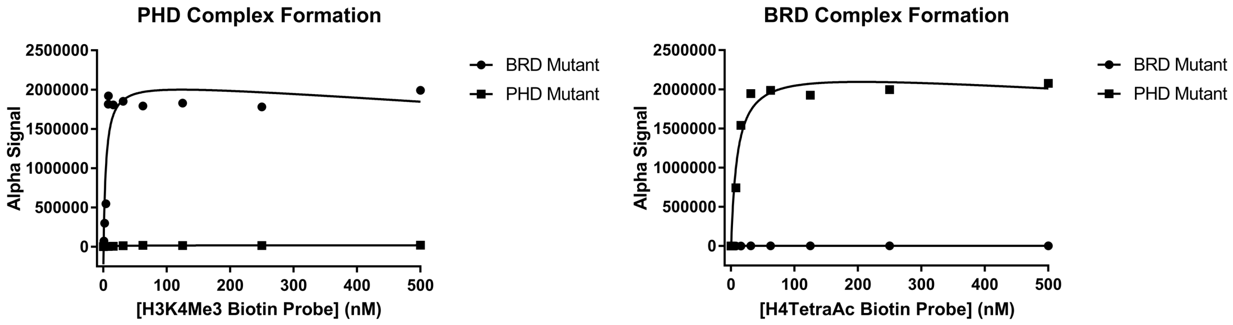

**Supplementary Figure S1.** AlphaScreen™ ternary complex formation with single domain mutants.

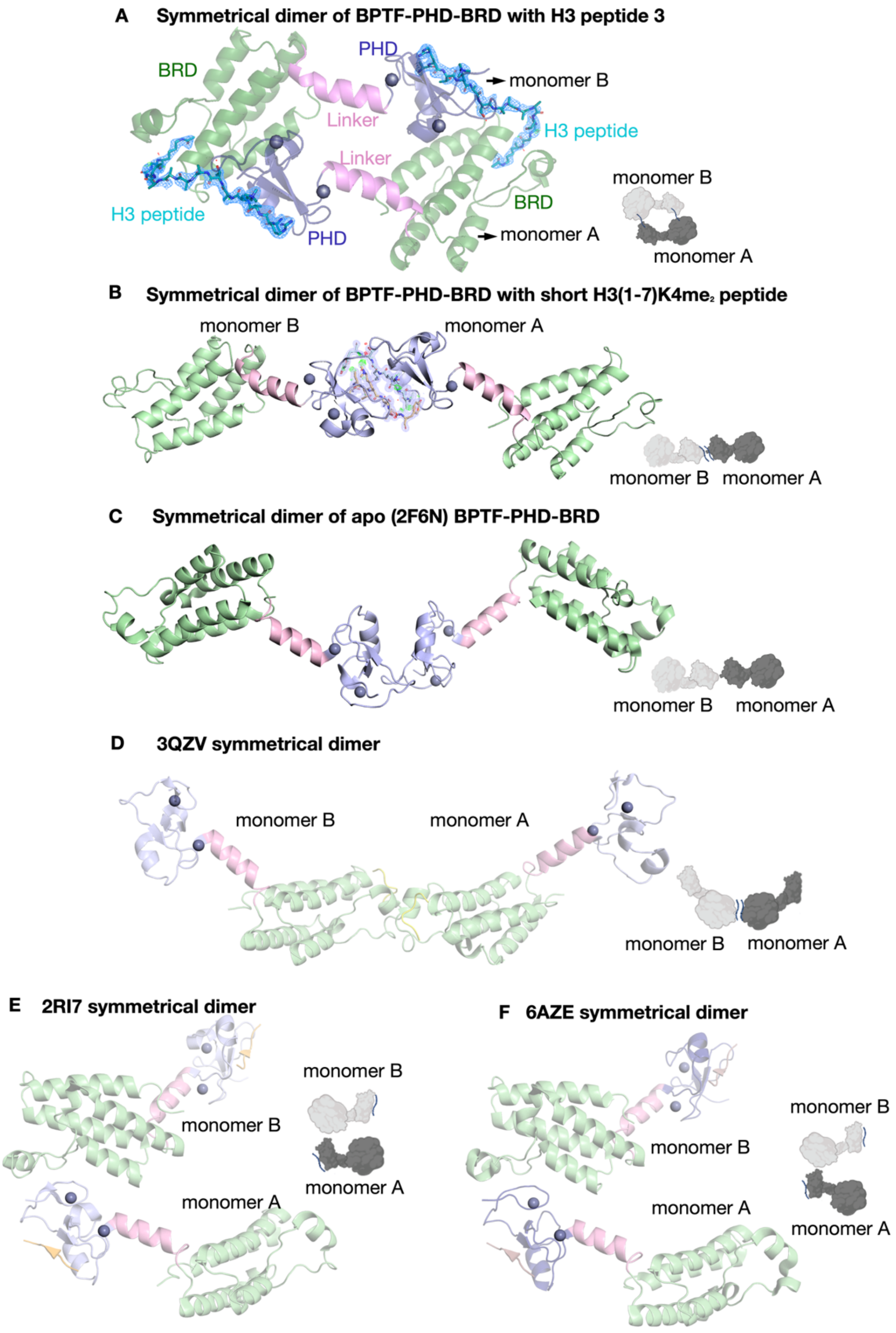
A Symmetrical dimer of BPTF-PHD-BRD with H3 peptide 3
BRD
PHD
monomer B
Linker
H3 peptide
Linker
monomer B
BRD
H3 peptide
PHD
monomer A
monomer A
B Symmetrical dimer of BPTF-PHD-BRD with short H3(1-7)K4me2 peptide
monomer B
monomer A
monomer B monomer A
C Symmetrical dimer of apo (2F6N) BPTF-PHD-BRD
monomer B monomer A
D 3QZV symmetrical dimer
monomer B
monomer A
monomer B monomer A
E 2RI7 symmetrical dimer
monomer B
monomer B
monomer A
monomer A
F 6AZE symmetrical dimer
monomer B
monomer B
monomer A
monomer A

**Supplementary Figure S2:** Dimer arrangement differs across BPTF-PHD-BRD structures. (**A**) Symmetrical dimer arrangement in the crystal structure of BPTF-PHD-BRD-peptide 3 complex. **B**) Symmetrical dimer of short K4me$_2$ peptide bound BPTF-PHD-BRD. 2Fo-Fc electron density is shown in blue at 1σ and Fo-Fc electron density is shown in green and red at 3σ and -3σ, respectively. (**C-F**) Symmetrical dimer arrangement of published BPTF-PHD-BRD structures, apo form[7] (**C**), bound with acetylated peptide[21] (**D**), bound with dimethylated peptide[22] (**E**), bound with trimethylated peptide[23] (**F**). Inset shown in all the panels  exemplify the various arrangement of symmetrical dimers.

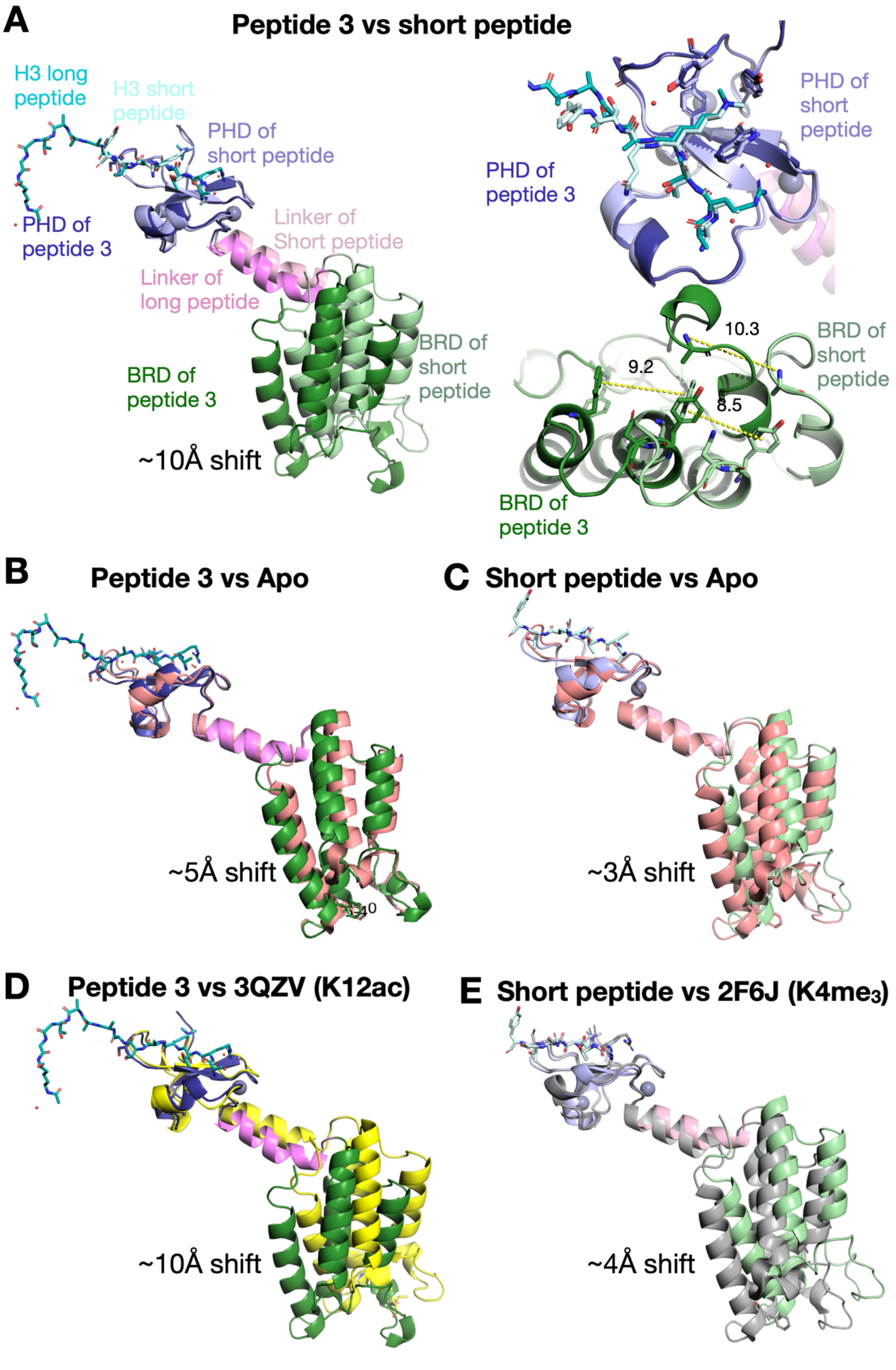
A
Peptide 3 vs short peptide
H3 long peptide
H3 short peptide
PHD of short peptide
PHD of peptide 3
Linker of Short peptide
Linker of long peptide
BRD of short peptide
BRD of peptide 3
~10Å shift
PHD of short peptide
PHD of peptide 3
10.3
9.2
8.5
BRD of short peptide
BRD of peptide 3
B
Peptide 3 vs Apo
~5Å shift
C
Short peptide vs Apo
~3Å shift
D
Peptide 3 vs 3QZV (K12ac)
~10Å shift
E
Short peptide vs 2F6J (K4me3)
~4Å shift

**Supplementary Figure S3:** Analysis of the relative flexibility of BPTF-PHD-BRD structures. Superposition of peptide **3** and H3(1-7)K4me$_2$ bound BPTF-PHD-BRD structures (**A- left**). Closeup views of PHD and BRD domains (A- right). Yellow dashed lines are distance between the respective amino acids. Structural alignment of peptide **3** bound and short peptide bound BPTF-PHD-BRD (**B**), H3(1-7)K4me$_2$ peptide and apo (2F6N) structure[7] (**C**), peptide **3** and 3QZV[21] (**D**), short peptide and 2F6J[7] (**E**). The structures were aligned at the PHD domain as a reference for all the pairwise comparisons.

**Supplementary Table 2**: Crystallographic data statistics: Values in parentheses are for high resolution bins.

| Parameter | BPTF-PHD-BRD-peptide 3 | BPTF-PHD-BRD-H3(1-7)K4me$_2$ |
|---|---|---|
| PDB ID | 38ZY | 38ZZ |
| Wavelength | 0.9199 | 0.9199 |
| Resolution range (Å) | 34.08 - 2.17 (2.49 - 2.17) | 38.08 - 1.6 (1.66 - 1.6) |
| Space group | C 2 2 21 | C 2 2 21 |
| Unit cell | a= 50.932, b= 56.047, c= 136.314, α= 90, β= 90, γ= 90 | a= 44.83, b= 61.068 c= 152.304, α= 90, β= 90, γ= 90 |
| Unique reflections | 9026 (1848) | 28044 (2755) |
| Multiplicity | 8.4 (8.0) | 8.2 (6.6) |
| Completeness (%) | 84.63 (53.03) | 99.90 (99.53) |
| Mean I/sigma(I) | 12.7 (1.1) | 18.0 (1.0) |
| Wilson B-factor | 42.1 | 25.07 |
| R-merge | 0.127 (1.469) | 0.053 (0.705) |
| R-meas | 0.144 (1.683) | 0.061 (0.834) |
| R-pim | 0.067 (0.813) | 0.029 (0.437) |
| CC1/2 | 0.998 (0.555) | 0.999 (0.850) |
| Reflections used in refinement | 9026 (1848) | 28044 (2755) |
| Reflections used for R-free | 437 (89) | 1362 (158) |
| R-work | 0.2246 | 0.2085 |
| R-free | 0.2497 | 0.2198 |
| Number of non-hydrogen atoms | 1500 | 1604 |
| macromolecules | 1445 | 1454 |
| ligands | 2 | 8 |
| solvent | 53 | 142 |
| Protein residues | 182 | 176 |
| RMS(bonds) | 0.012 | 0.006 |
| RMS(angles) | 1.67 | 0.83 |
| Ramachandran favored (%) | 97.08 | 98.22 |
| Ramachandran allowed (%) | 2.92 | 1.78 |
| Ramachandran outliers (%) | 0.00 | 0.00 |
| Rotamer outliers (%) | 0.68 | 0.00 |
| Clash score | 3.22 | 2.10 |
| Average B-factor | 50.84 | 30.41 |
| macromolecules | 50.80 | 29.89 |

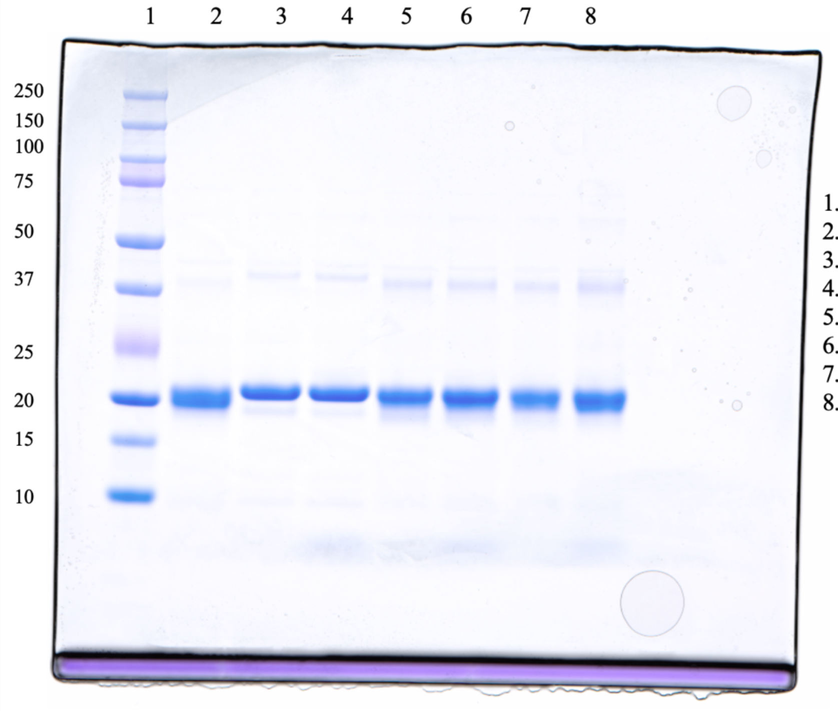

**Supplementary Figure S4.** SDS-PAGE of DHSO/DMTMM mediated crosslinking of BPTF with and without peptide **3** at protein and ligand concentrations that match or exceed those used in biophysical assays. DHSO is an acidic residue reactive cross linking agent used to covalently trap protein-protein interactions that form higher-ordered oligomeric protein species.[10,24] Lanes 3-8 contain 400 µM DHSO and 800 µM DMTMM. Monomer MW = 20.1 kDa; dimer MW = 40.2 kDa. Most of the sample remains in the uncrosslinked, monomeric state with a minor amount of pre-existing dimer that was not induced by the presence of **3**.

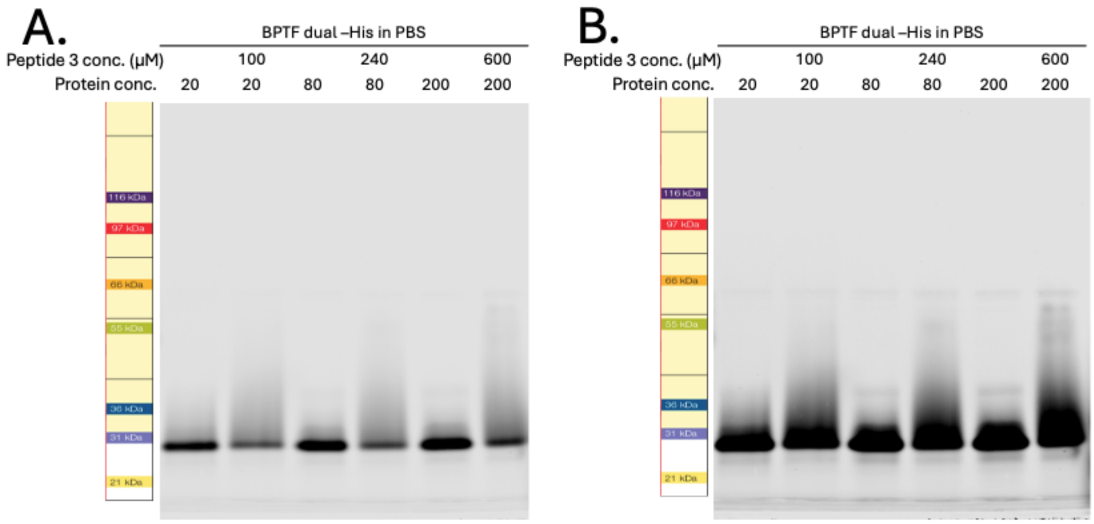

**Supplementary Figure S5**. Native-PAGE of BPTF dual domain with and without peptide **3** added at concentrations that match or exceed biophysical assays. Native PAGE preserves protein structure and allows for the separation of proteins that adopt multiple oligomeric states.[25,26] Ladder indicates expected movement through the 4-12% Tris Glycine gel in percentage as predicted by Thermo Fisher. Monomer MW = 20.1 kDa; dimer MW = 40.2 kDa. **A**) Image with normal exposure and **B**) high exposure. BPTF-PHD-BRD largely favors a single oligomeric state in solution that corresponds to a MW < 30 kDa.

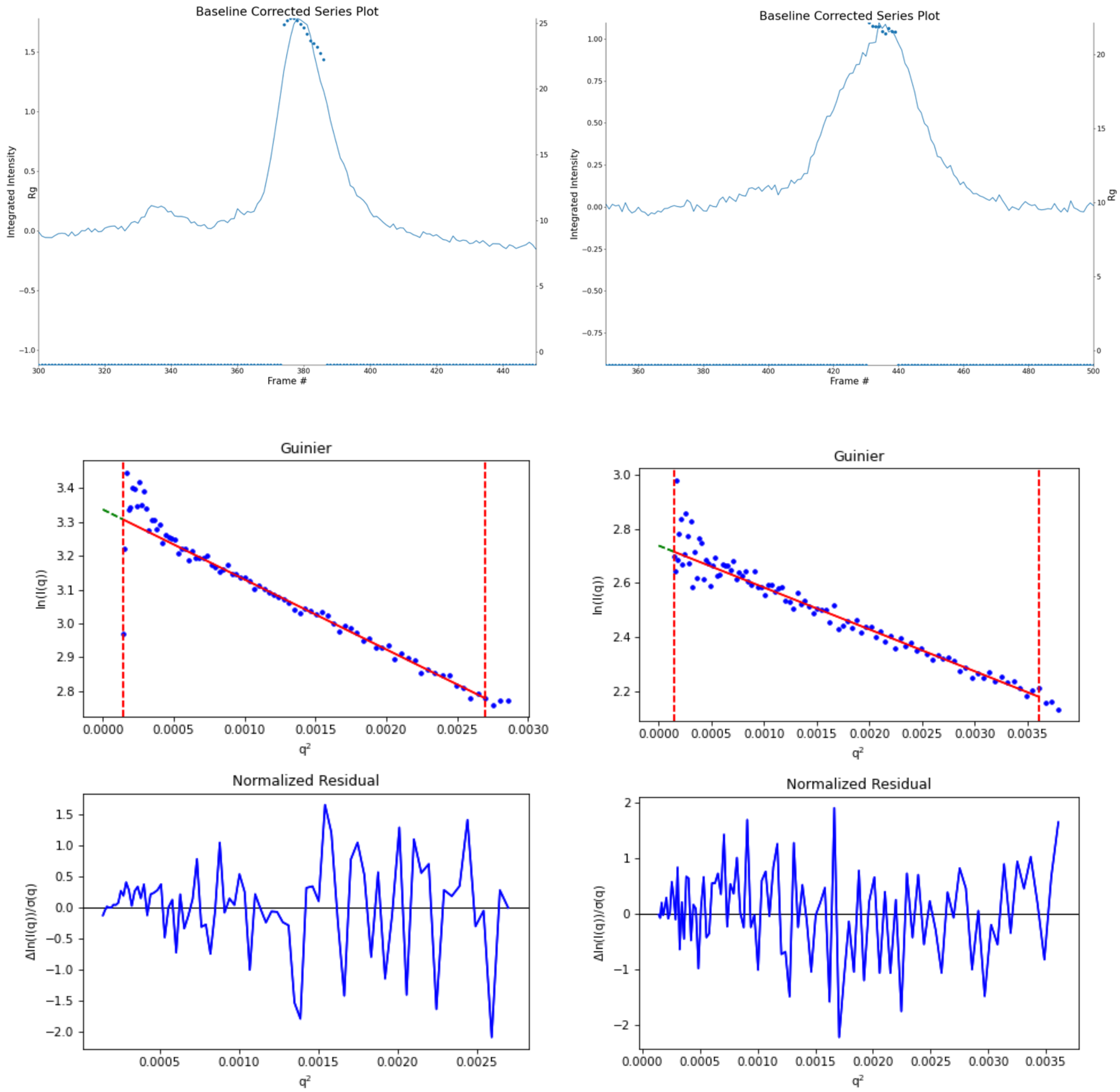

**Supplementary Figure S6.** SEC peaks and Guinier analyses of SAXS data without peptide 3 present (left) and peptide 3 bound (right).

**Supplementary Table 3.** Distribution of conformational ensembles produced by MultiFoXS with unliganded BPTF-PHD-BRD (left) and bound to peptide 3 (right).

| # States | Radius of Gyration | Proportion | % of Population < 22 Å | % of Population > 22 Å |
|---|---|---|---|---|
| One | 22.1 Å | 100% | 0% | 100% |
| Two | 24.3 Å | 57% | 43% | 57% |
| | 19.5 Å | 43% | | |
| Three | 21.4 Å | 20% | 48% | 52% |
| | 24.3 Å | 52% | | |
| | 19.2 Å | 28% | | |
| Four | 20.7 Å | 34% | 53% | 47% |
| | 24.3 Å | 31% | | |
| | 19.2 Å | 19% | | |
| | 26.2 Å | 16% | | |

| # States | Radius of Gyration | Proportion | % of Population < 22 Å | % of Population > 22 Å |
|---|---|---|---|---|
| One | 19.4 Å | 100% | 100% | 0% |
| Two | 18.2 Å | 72% | 72% | 28% |
| | 24.1 Å | 28% | | |
| Three | 18.2 Å | 36% | 75% | 25% |
| | 19.0 Å | 39% | | |
| | 28.3 Å | 25% | | |
| Four | 18.2 Å | 44% | 75% | 25% |
| | 19.0 Å | 31% | | |
| | 24.1 Å | 8% | | |
| | 28.3 Å | 17% | | |

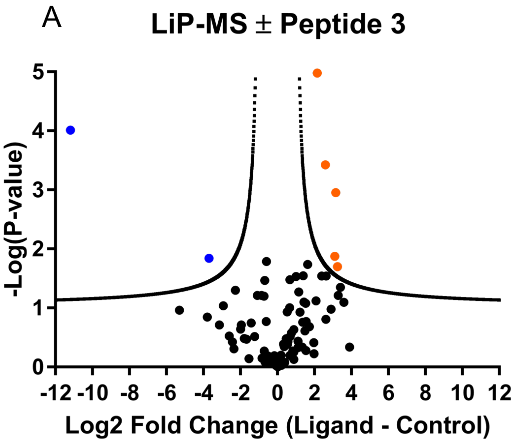

| Peptide | Fold Change | -Log(P-value) |
|---|---|---|
| LEPVDPNDAPDYYGVIKEPMDLATM | -11.2 | 4.013 |
| LEPVDPNDAPDYYGVIKEPMDLATME | -3.70 | 1.843 |
| CQSTEDAMTVLTPLTEKDYEGLK | +3.25 | 1.703 |
| EAELIDEYVCPQCQSTEDAMTVLTPLTEKDYEGLKR | +2.59 | 3.427 |
| PNDAPDYYGVIKEPMDLATMEER | +3.16 | 2.955 |
| QSEAELIDEYVCPQCQSTEDAMTVLTPLTEKDYEGLKR | +3.10 | 1.876 |
| STEDAMTVLTPLTEKDYEGLKR | +2.16 | 4.982 |

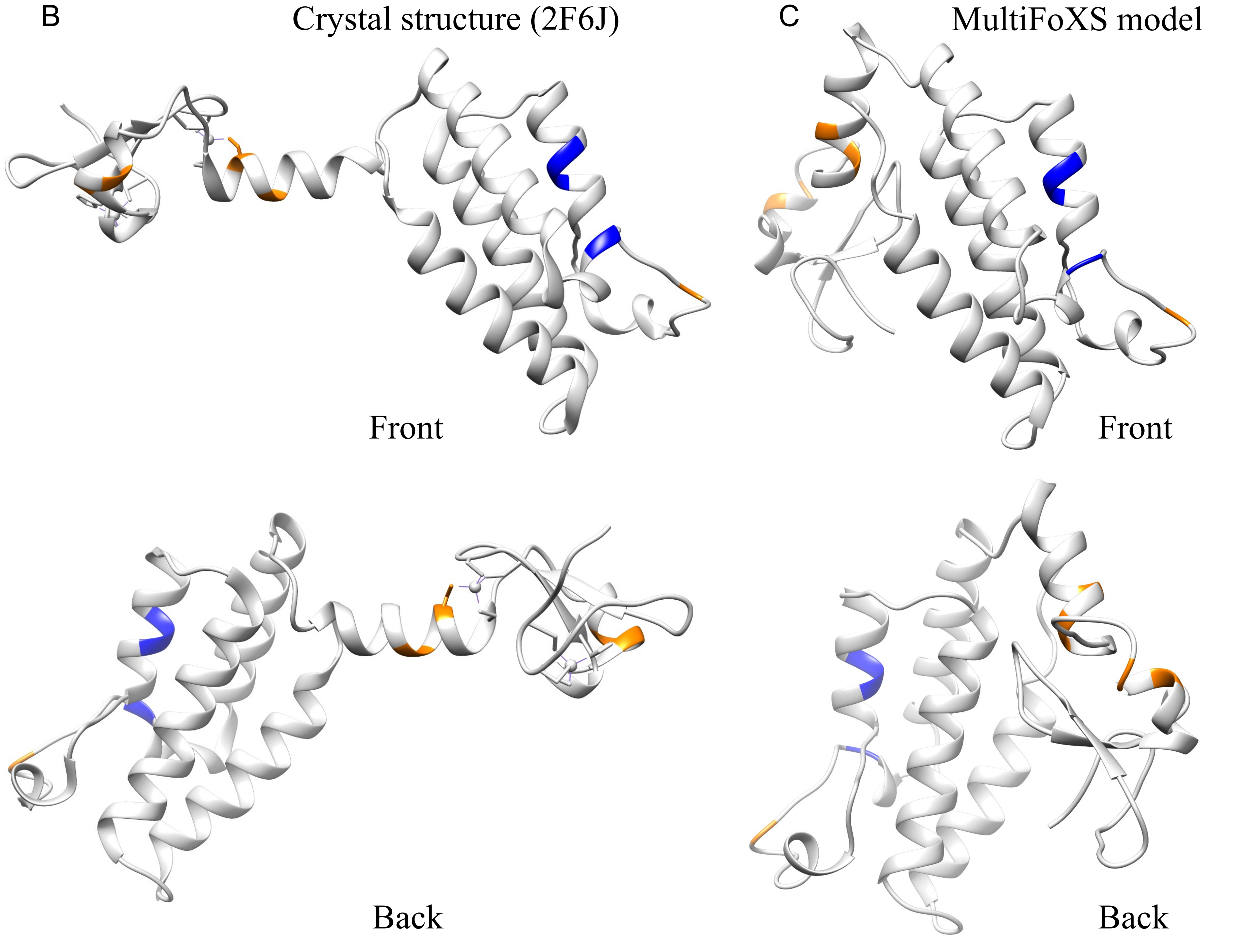

**Supplementary Figure S7.** A.) Volcano plot (left) and table (right) of conformotypic peptides downregulated (blue) or upregulated (right) upon treatment with **3**. Downregulated (blue) and upregulated (orange0 proteinase K cleavage sites mapped to an existing BPTF-PHD-BRD crystal structure (B) and to the MultiFoXS model (C) generated from the peptide bound SAXS data ($R_g$ = 19.0 Å)

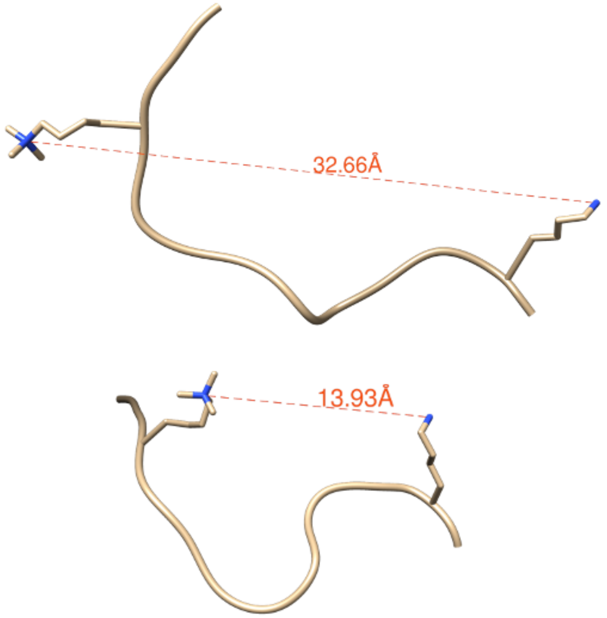

| ID | Atom 1 | Atom 2 | Distance |
|---|---|---|---|
| 1 | #0.1 LYS 14.B NZ | #0.1 M3L 4.B NZ | 32.02Å |
| 2 | #0.10 M3L 4.B NZ | #0.10 LYS 14.B NZ | 21.31Å |
| 3 | #0.11 M3L 4.B NZ | #0.11 LYS 14.B NZ | 24.24Å |
| 4 | #0.12 LYS 14.B NZ | #0.12 M3L 4.B NZ | 17.65Å |
| 5 | #0.13 LYS 14.B NZ | #0.13 M3L 4.B NZ | 20.74Å |
| 6 | #0.14 M3L 4.B NZ | #0.14 LYS 14.B NZ | 23.89Å |
| 7 | #0.15 LYS 14.B NZ | #0.15 M3L 4.B NZ | 26.53Å |
| 8 | #0.16 LYS 14.B NZ | #0.16 M3L 4.B NZ | 23.66Å |
| 9 | #0.17 LYS 14.B NZ | #0.17 M3L 4.B NZ | 13.93Å |
| 10 | #0.18 LYS 14.B NZ | #0.18 M3L 4.B NZ | 30.45Å |
| 11 | #0.19 LYS 14.B NZ | #0.19 M3L 4.B NZ | 17.99Å |
| 12 | #0.2 M3L 4.B NZ | #0.2 LYS 14.B NZ | 19.19Å |
| 13 | #0.20 LYS 14.B NZ | #0.20 M3L 4.B NZ | 29.15Å |
| 14 | #0.3 LYS 14.B NZ | #0.3 M3L 4.B NZ | 29.81Å |
| 15 | #0.4 M3L 4.B NZ | #0.4 LYS 14.B NZ | 28.79Å |
| 16 | #0.5 LYS 14.B NZ | #0.5 M3L 4.B NZ | 32.66Å |
| 17 | #0.6 LYS 14.B NZ | #0.6 M3L 4.B NZ | 26.24Å |
| 18 | #0.7 LYS 14.B NZ | #0.7 M3L 4.B NZ | 24.20Å |
| 19 | #0.8 LYS 14.B NZ | #0.8 M3L 4.B NZ | 26.98Å |
| 20 | #0.9 M3L 4.B NZ | #0.9 LYS 14.B NZ | 17.55Å |

**Supplementary Figure S8.** Distances (Å) measured between H3K4me$_3$ and H3K14 amine side chains from 20 individual poses modeled in histone H3(1-15)K4me$_3$ peptide NMR solution structure when bound to BPTF PHD (PDB ID 2FUU), measured using the “distances” tool in UCSF Chimera.

**BPTF-PHD-BRD**

A

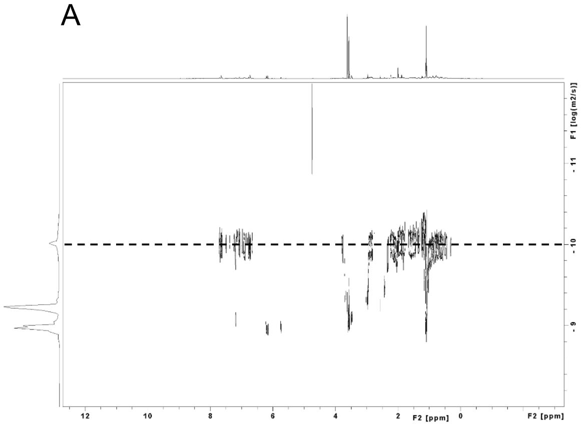

**5**

B

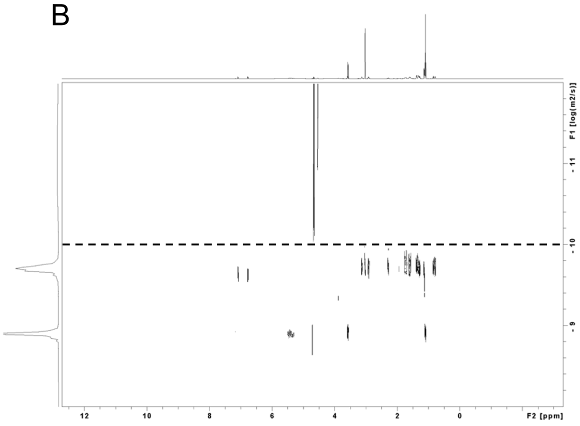

C **1:1 BPTF-PHD-BRD:5**

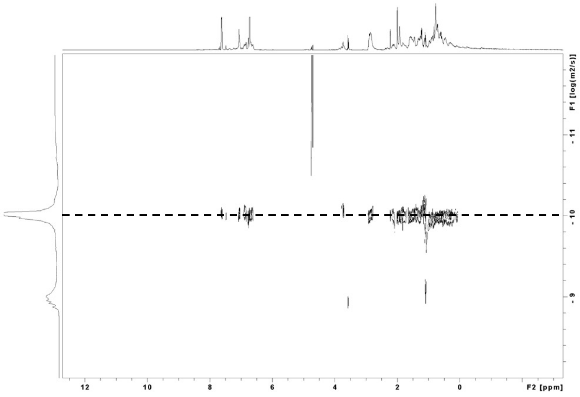

D **3**

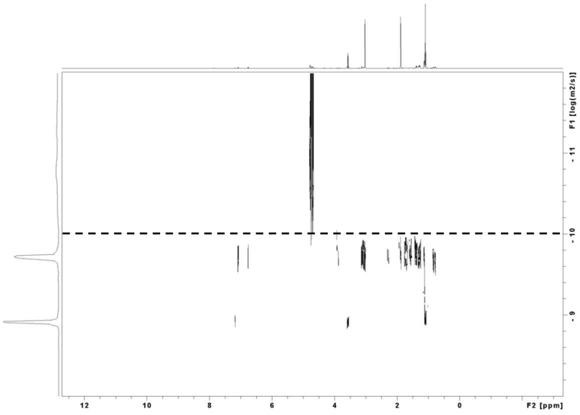

**1:1 BPTF-PHD-BRD:3**

E

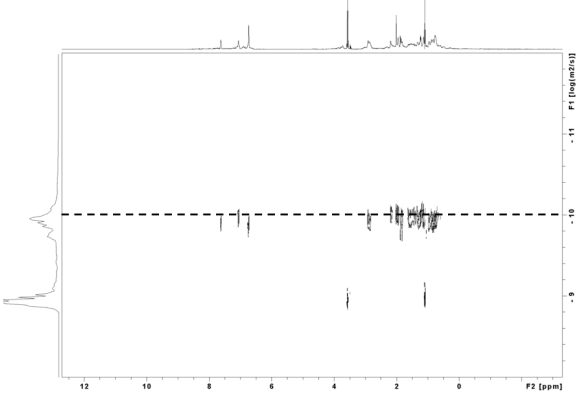

**Larger radius, lower *D***

**Smaller radius, higher *D***

| **Condition** | **Diffusion Coefficient (*D*) (x$10^{-10}m^2/s$)** |
|---|---|
| BPTF-PHD-BRD Apo | 3.4 |
| **5** Apo | 19 |
| **3** Apo | 11 |
| 1:1 BPTF-PHD-BRD:**5** | 2.2 |
| 1:1 BPTF-PHD-BRD:**3** | 4.8 |

**Supplementary Figure S9**. DOSY NMR spectra with BPTF-PHD-BRD alone (**A**), peptides **5** and **3** alone (**B and D**, respectively), and BPTF-PHD-BRD bound to peptide **5** (**C**) and bound to peptide **3** (**E**). DOSY can provide information on different oligomeric and conformational states of protein complexes by calculation of diffusion coefficients (*D*), which are inversely related to hydrodynamic radii.[27,28] Binding of peptide **3** results in a larger *D* value relative to the unbound BPTF-PHD-BRD ($D$ = 4.8 x $10^{-10}$ $m^2/s$ versus 3.4 x $10^{-10}$ $m^2/s$, respectively), suggesting this protein-peptide complex adopts a conformation that diffuses faster in solution and corresponds to a smaller hydrodynamic radius as changes in *D* that exceed 20% often reflect mass shifts and macromolecular structural rearrangement events.[29] This contradicts an expected decrease in *D* if a larger, dimeric species was forming upon peptide **3** addition.[30]

**Supplementary Table 4.** Chemical matter of PHD-finger containing proteins. Compounds selected from their respective campaigns have both $K_d$ and functional $IC_{50}$ values reported or are the most potent binder. Amiodarone and disulfiram exhibit broad spectrum, assay-dependent activity for many PHD-finger containing proteins including KDM5A PHD3, KDM7A/7B/7C, BPTF PHD.[4,31,32,33] MLD5 also inhibits the AIRE PHD ($IC_{50}$ = 77 µM; FP).

| Protein | Molecule | $K_d$ | $IC_{50}$ | Assay(s) |
|---|---|---|---|---|
| BAZ2A/2B | F23[34] | 7100 – 9,700 µM | 100 – 200 µM | HSQC NMR, AlphaLISA |
| UHRF1 PHD | MLD5[35] | - | 7.4 µM | FP, AlphaScreen |
| PYGO PHD | CF16[36] | - | 7300 µM | HSQC NMR |
| BPTF PHD | F4[4] | - | 130 µM | AlphaScreen |
| KDM5A PHD1 | B2[37] | - | 1300 µM | FP |
| KDM7B PHD | OC9[38] | 12 nM | 32 nM | ITC, AlphaScreen |
| TRIM24 PHD | PDC6[39] | 0.62 nM | 11 nM | SPR, AlphaScreen |

[38] Coleman, O. D.; Macdonald, J.; Thomson, B.; Ward, J. A.; Stubbs, C. J.; McAllister, T. E.; Clark, S.; Amin, S.; Cao, Y.; Abboud, M. I.; Zhang, Y.; Sanganee, H.; Huber, K. V. M.; Claridge, T. D. W.; Kawamura, A.

Cyclic Peptides Target the Aromatic Cage of a PHD-Finger Reader Domain to Modulate Epigenetic Protein Function. *Chem. Sci.* **2023**, *14* (26), 7136–7146.

[39] Platt, M. A.; Kot, E.; Martin, L. A. W.; Wallabrègue, A. L. D.; Song, L.; Boyd, A. M.; Koekemoer, L.; Hammond, E. M.; Conway, S. J. High-Affinity Peptide-Drug Conjugate Ligands for the TRIM24 PHD and Bromodomain. *Chem. –Eur. J.* **2025**, *31* (72), e03011.
